# Olfactory Spatial Attention: Neural Structural Connectivity Revealed

**DOI:** 10.1101/483446

**Authors:** Ganesh Elumalai, Harshita Chatterjee, Imran Shareef Syed, Valencia Brown, Lekesha Adele Sober, Nitya Akarsha Surya Venkata Ghanta, Pradeep Chandrasekhar, Anbarasan Venkatesan, Nadira Sewram, Nitesh Raj Muthukumaran, Golla Harkrishna, Nanduri Mojess Vamsi, Nneoma Somtochukwu Osakwe Team NeurON, Texila American University

## Abstract

Different aspects of spatial neglect, the disability to locate the position of the source of an external stimulus, has been studied frequently especially with regards to lack of response to visual stimuli. A modicum of papers fails to elaborate on that of the auditory field and even less conducted on the extraction of spatial information from olfactory stimuli. In this study we attempted to find a structural connection between the Olfactory Cortex (OC) and Temporoparietal Junction (TPJ), in order to suggest that processed olfactory information is carried from OC to TPJ, where the spatial information of the odor maybe acquired. Positive results were attained in all subjects with a lateralization observed in the right hemisphere.

**One Sentence Summary:** Existence and hemispherical dominance.

---

Olfactory identification involves the coalesced code molded by particular patterns of excitatory insert from olfactory receptor neurons. This combinatorial coding mechanism was postulated to be contributed by the spaciotemporal patterning of active neurons in olfactory centers *(1)*. The olfactory neural process underlying spatial recognition encompasses the computation of molecular information in constructing an image and possibly the location of a particular odorant *(2, 3)*. Information with regards to space may be configured by place cells when subject is situated in a specific environment, with or without visual cues. Non-visual modalities such as olfaction provide a constellation of the environment around to generate spatial recognition of odorants *(3)*. Documentation of the hippocampus following such is represented well, especially with experimentation involving the suppression of visual and auditory cues *(4)*. The temporal-parietal junction (TPJ) contributes to spatial recognition in the processing of visual, auditory and somatosensory stimuli, including information from the thalamus and the limbic system *(5).* Research done on monkeys revealed area 7 to be of high importance, being able to produce deficits in spatial attention. This area 7 in monkeys have been thought to correspond to angular gyrus (BA area 39) and supramarginal gyrus (BA area 40) in humans, along with superior temporal gyrus (BA area 22) which suggests visual perception and attention through fibers from the visual cortex and inferior parietal lobe *(5-7).* The right hemisphere of the cerebrum containing TPJ tend to be more specific for spatial intelligence and recognition, since spatial deficits are noted to be seen in patients with right TPJ lesions *(8-9)*. TPJ also functions in reorienting cortical response to task-oriented but currently unattended endogenous or exogenous stimuli *(8-13).* Two concepts are introduced in relation to olfactory attention and consciousness, ‘orienting’ and ‘detecting’. Orienting being the capability of aligning one’s focus to a particular sensory input (overt) or from internal memory (covert); and the detection of such stimuli has to be at a concentration where the nervous system is able to perceive it *(14-15).* Olfactory attention can be shifted in space by positioning the nose over odorants by allowing us to locate sources of odor and to track scents. It has been suggested that olfactory experience has a very simple spatial structure. Pathways from the piriform cortex to the orbitofrontal cortex is used to process odor information, but literature fails to mention projections to other brain regions as it relates to spatial recognition. Olfaction, in contrast to other sensory inputs do not pass through the thalamus to obtain such spatial processing. Although processing of odors in relation to spatial recognition commences at the olfactory bulb, the effect of a right resected temporal lobectomy demonstrated a decline in olfactory attention that is necessary for olfactory consciousness. This relation considers the probability that spatial recognition processing does occurs at the TPJ *(16).* Extensive studies done to analyze the response of TPJ to auditory and visual stimuli have been conducted, but not much work has been done in order to analyze the effect of olfactory stimuli on response generated by the TPJ in humans. It then puts into question what contributes to spatial attention concerning olfactory cues, further questioning whether the temporal-parietal junction is in fact playing a role in spatial recognition of olfaction as it does for other sensory inputs. The aim of this study is to provide a structural connection from the olfactory cortex (OC) to the TPJ. We obtained the structural Magnetic resonance imaging (sMRI) scans of 20 clinically normal subjects, including eleven females and ten males of various age groups (Table S1), and traced fibre tracts extending from the OC to the TPJ *(17).*

## Result

The results found have been divided in two sets. The first set includes comparison of the different attributes of the fibre tracts found in male and female subjects.

All the subjects displayed a wide-ranging number of tracts (Graph 1). Fig. S1 – Fig. S21 include images of tracts found in the subjects when the region merged Olfactory Cortex (mOC) was selected as the seeding region and the region merged Temporoparietal Junction (mTPJ) was selected as the ending region. These merged regions are a combination of the parts of the regions in the two different hemispheres, mOC is a combination of right hemisphere OC (rOC) and left hemisphere OC (lOC), similarly mTPJ is a combination of right hemisphere TPJ (rTPJ) and the left hemisphere TPJ (lTPJ). In the first set of fibre tracing, fibres only extending between merged regions have been considered, since a hemispherical comparison is not being made, and a primary association is being established before making any classifications.

Table S2 and Graph S1 compare the number of tracts found in the merged regions in male and female subjects. The significance of the difference in the average of the two samples (male and female) was assessed through a t-test calculation. The results were recorded as - mean number or tracts in females (Mf) = 40.5, degree of freedom in females (dff) = 9. Mean number of tracts in males (Mm) = 62.1, degree of freedom in males (dfm) = 9. The t-value is −0.78723. The p-value is 0.441389. The result is not significant at p < .05. This result maintains consistency with several other studies conducted stating that there are no significant differences between the sexes when response to stimuli is being compared. In this study, we suggest that the number of fibres may be directly proportional to the amount of attention drawn towards an olfactory stimulus.

Table S3, Graph S2 and Graph S3 compare the mean length of tracts found in the merged regions in male and female subjects. The results of significance of the difference in the average were recorded as – Mf = 98.19 mm, dff = 9. Mm = 98.9 mm, dfm = 9. The t-value is −0.19732. The p-value is 0.845788. The result is not significant at p < .05. The length of the fibres also show no major difference in the sexes, suggesting that the regions traversed by them is similar in the sexes, following a similar route.

Table S4, Graph S4 and Graph S5 compare the volume of tracts found in the merged regions in male and female subjects. The results of significance of the difference in the average were recorded as – Mf = 4083.08 mm3, dff = 9. Mm = 3980.48 mm3, dfm = 9. The t-value is 0.0748. The p-value is 0.941198. The result is not significant at p < .05. Higher the number of fibres more is the volume of the fibre tracts. Therefore, the result observed in this comparison maintains consistence with the result obtained from the comparison of number of tracts between the sexes.

The second set includes a hemispherical comparison of the fibres as seen collectively and in the male and female subjects.

This set of results were derived from tracing fibres extending from mOC to lTPJ and rTPJ (Fig. S21 - S40).

Table S5, Graph S6 and Graph S7 compare the number of tracts found ending in lTPJ and rTPJ starting from mOC regions in subjects. The significance of the difference in the average of the two samples (left and right hemisphere) was calculated. The results were recorded as - mean number or tracts ending in lTPJ (Ml) = 10.1, degree of freedom (dfl) = 19. Mean number of tracts ending in rTPJ (Mr) = 41.2, degree of freedom (dfr) = 19. The t-value is −2.719. The p-value is .009814. The result is significant at p < .05. In 85.71% of the subjects, the rTPJ receives a number of fibers than the lTPJ, in 9.52% of the subject the lTPJ receives a greater number of fibres than the rTPJ and in 4.76% of subjects, the number of fibres received by lTPJ and rTPJ is equal. Most of the fibres received in either TPJ are ipsilateral in nature with as few as 1-5 fibres originating in OC of the opposite side, among those subjects that displayed this feature, the fibres originating from an opposite OC were mostly from lOC, which then entered rOC and extended to rTPJ along with the bundle of fibres that originated in the rOC, no separate path taken by these fibres was noted. This suggests that information processed in rTPJ is affected by the stimuli processed in lOC, and damage or absence of this fibre may affect the output produced by rTPJ, and also lTPJ given that rTPJ and lTPJ directly interact.

Table S6, Graph S8 and Graph S9 compare the mean length of tracts found ending in lTPJ and rTPJ starting from mOC in subjects. The results were recorded as - Ml = 89.31 mm, dfl = 19. Mr = 96.42 mm, dfr = 19. The t-value is −0.77936. The p-value is 0.440593. The result is not significant at p < .05. This suggests that there is no variance seen in the aspect of attention produced in the hemispheres which is directed by the length of fibres.

Table S7, Graph S10 and Graph S11 compare the volume of tracts found ending in lTPJ and rTPJ starting from mOC in subjects. The results were recorded as - Ml = 1289.08 mm3, dfl = 19. Mr = 3023.49 mm3, dfr = 19. The t-value is −3.19782. The p-value is 0.00279. The result is significant at p < .05. This maintains uniformity with the comparison between number of fibres, and establishes that the right hemisphere receives more fibres in both sexes as compared to the left hemisphere, suggesting that right hemisphere may be more responsible for generation of attention in response to the stimuli.

Considering that we found significant differences in fibre number and tract volume in right and left hemisphere, we proceeded to find a pattern in the distribution of tract number and volume in the right and left hemispheres in male and female subjects (Table S8, Graph S12, Graph S13). In the female subjects in terms of tract numbers, we noted Mr = 29.8, dfr =9. Ml = 10.7, dfl = 9. The t-value is 1.95988. The p-value is 0.065679. The result is significant at p < .05. In male subjects, we noted Mr = 52.6, dfr =9. Ml = 9.5, dfl = 9. The t-value is 2.08221. The p-value is 0.051864. The result is not significant at p < .05. In terms of volume, in females, we noted Mr = 2768.51 mm3, dfr =9. Ml = 1567.35 mm3, dfl = 9. The t-value is 1.71731. The p-value is 0.103078. The result is significant at p < .05. In males we noted Mr = 3278.47 mm3, dfr =9. Ml = 1010.81 mm3, dfl = 9. The t-value is 2.67029. The p-value is.015604. The result is significant at p < .05. From this set of results, we concluded that the tract number and volume are both significantly different in the two hemispheres in females, with more fibres seen ending on the right hemisphere; in male subjects it was noted that while he volume of fibres ending is significantly higher in the right hemisphere, the number of fibres ending in the two hemispheres on TPJ are more or less similar. So, while we can say that the right hemisphere is dominant in production of attention in females, the question still remains in case of males.

In Table S9, Graph S14 and Graph S15, we have compared the number and volume of tracts in each hemisphere between male and female subjects. In terms of tract number in the right hemisphere, it was recorded that Mf = 29.8, dff = 9. Mm = 52.6, dfm = 9. The t-value is - 1.03103. The p-value is 0.316183. The result is not significant at p < .05. In the left hemisphere, it was recorded that Mf = 10.7, dff = 9. Mm = 9.5, dfm = 9. The t-value is 0.20458. The p-value is 0.840197. The result is not significant at p < .05. In terms of volume in right hemisphere, it was recorded that Mf = 95.26 mm3, dff = 9. Mm = 2975.87 mm3, dfm = 9. The t-value is - 3.51029. The p-value is 0.002499. The result is not significant at p < .05. In the left hemisphere, it was recorded that Mf = 1567.35 mm3, dff = 9. Mm = 1010.81 mm3, dfm = 9. The t-value is 1.12418. The p-value is 0.275707. The result is not significant at p < .05. After this set of results was obtained, we can conclude that there is no difference whatsoever in male in female subjects when it comes to hemispherical dominance.

Finally, we performed a comparison of tract numbers, mean length and volume between different age groups, and spotted a pattern that shows gradual increase and then decrease with progression in age (Graph S16, Graph S17, Graph S18), this suggests that the fibres traced show degeneration after a certain age, following which the attention drawn by olfactory stimuli may decrease considerably.

## Discussion

The present study performs tractography in order to establish a neural structural connectivity between the OC and TPJ *(Fig.1)*, and performs comparative analysis between sexes and hemispheres. Our initial hypothesis suggested that there exists a structural connectivity between the regions which facilitates orientation guided by spatial attention stimulated by odors. Our first set of results, upholds the same. We noted a vast range of fibres ranging from 2 to 256. We followed up our findings with a comparison between the sexes and found no significant difference between the number, mean length or volume of fibres running from the mOC to mTPJ. Our secondary hypothesis suggested that there would be a difference in the connectivity when compared between the hemispheres. This hypothesis stems from the fact that many studies conducted on the hemispherical comparison between the function of TPJ in spatial attention, especially guided by visual stimuli, has produced results suggesting that rTPJ plays a more prominent role in the same *(8, 9).* In the second set of results we observe that, a significant difference exists in the number and volume of tracts ending in the two hemispheres, favorably lateralized to the right hemisphere, this upholds our secondary hypothesis as well. Upon the confirmation of our secondary hypothesis, we proceeded with further comparison. As because we had not found any significant difference in mean length in the preceding analysis, we only considered the tract number and volume in the following comparisons. In women, the tract number difference in right and left hemisphere was significant, however, the difference was not significant in men. When the same was tested in terms of volume, the difference in the hemispheres was significant for both men and women. We then compared the number and volume of tracts in each hemisphere as noted in men vs women, only to find that there were no differences at all in any comparison done between men and women.

In studies conducted in the past it has been establishes that when spatial cues are applied in order to assess the attention drawn towards it in response, results are remarkably stable and no differences are observed when compared between sexes *(18).* And in some studies, assessing the behavioral response times to stimuli (mostly visual), although women were seen responding slower than men but the p value remained 0.05 or higher *(19-21)*, therefore the difference is insignificant. This could be parallel to spatial attention in response to olfactory stimuli as well.

Differences in cerebral hemispheres indicates different roles in relation to event order *(22).* Results show few fibres from the rTPJ decussating to the lTPJ. It was mentioned that TPJ is involved in lower-level computational processes/ spatial attention orientation, visual search duration, theory of the mind and empathy *(23-25).* Damage to this area affects awareness of one’s self and disorders associated with body knowledge including out of body experiences with respect to experienced unity, self-location, and egocentric visuospatial perspective *(23, 26, 27).* Left hemispatial neglect was seen in stroke patients with lesions to the right angular gyrus *(28).* This spatial overlap is crucial in interactions between word production and word perception *(29)* Due to this relation, the assumption that the rTPJ contributes to olfactory spatial attention of lTPJ is likely.

lTPJ comprises of the Wernicke’s area which is responsible for language comprehension, semantic knowledge of single words and pre-semantic processing of faces and names, low-level social perception and high-level social reasoning *(24, 29, 30).* Damage to lTPJ causes Wernicke’s aphasia and disrupts cognitive processes of mental state such as someone’s belief *(24,31).* lTPJ also encodes multisensorial (eg. visual and auditory) temporal order judgement (TOJ) serving as an integral part of the ‘when’ pathway. This allows an individual to process the timings of two spatially discrete stimuli. Conduction of this experiment is used to assess TOJ by allowing patients to perform two tasks by discriminating temporal order and spatial properties. This is useful in the sense of investigating perception for subjective simultaneity (bias) and noticeable difference (sensitivity). Known is the fact that temporal order judgment of visual stimuli is impacted by detection and identification expedited by attention. Planning of saccades to a specific location and a particular stimulus, as in dual task (DT) paradigm, prior influences increasing speed of perceptual processing. With this given information, one can deduce that perception processing of stimulus occurs first then temporal order judgement. This relates to the topic at hand for TOJ is influenced greatly by spatial attention *(24, 31, 32).* Results showing fibres decussating from the rTPJ to the lTPJ were hypothesized by the current researchers to be contributing to the ‘when’ pathway of olfactory stimuli. rTPJ spatial attentional activity have been noted to interfere with temporal order judgement producing biases, but does not reduce its performance. For an unbiased result in TOJ assessment, one is to withstand any spatially induced stimuli or attentional variables that may trigger the involvement of rTPJ. Research has indicated the ‘what’, ‘where’ and ‘when’ processing streams to be identical and follow the same organization *(23).*

## Materials and Methods

The present study aims to establish a structural connection between the Olfactory Cortex (OC) and Temporoparietal Junction (TPJ).

### Datasets Acquisition

The study used an open access, ultra-high b-value, various diffusion sensitizing directions (64,128 and 256) diffusion imaging datasets with the in-plane resolution and slice thickness of 1.5 mm. The data sets are originally developed and reposted by Massachusetts General Hospital – US Consortium Human Connectome Project (MGH-USC HCP) *(33,34).*

The present study involves Twenty healthy adult datasets (10 males and 10 Females, between the age 20–59 years old; mean age = 30.4). Given de-identification considerations, age information is provided in 5-year age bins. All participants gave their written informed consent, and the experiments were carried out with approval from the Institutional Review Board of Partners Healthcare of MGH-USC HCP project. The demographic details of the participants with gender and age are available in the data sharing repository *(35-53).*

### Data Pre-processing and Fibre tractography

The MGH-USC HCP team has completed their basic imaging data pre-processing, with software tools in Free surfer and FSL, which includes

i. Gradient nonlinearity correction,
ii. Motion correction,
iii. Eddy current correction,
iv. b-vectors.

#### Region selection

For initial tracing, to establish the existence, two regions were used, merged Olfactory Cortex (mOC) and merged Temporoparietal Cortex (mTPJ). Here, mOC was selected as the seeding region and mTPJ was selected as the ending region, with seeding limit set on 100,000. This implies that the fibre tracing would end once 100,00 fibres were found that begin running from mOC and terminate in mTPJ.

For the next set of tracing performed, to find a pattern in distribution of fibres between the hemispheres, three regions were used, mOC, left hemisphere temporoparietal cortex (lTPJ) and right hemisphere temporoparietal cortex (rTPJ). Here, mOC was selected as the seeding region, lTPJ and rTPJ were selected as the ending region, with seeding limit set on 100,000.

Table S10 contains region based statistical data for each region.

### Statistical Analysis

In order to analyze the significance of the difference between the mean of the two sets of data under each of the two categories, as seen in Table S1-S9, we performed a t-test calculation for two independent means, considering a two-tailed hypothesis.

The significance level was set on 0.05.

## Supplementary Text

Fiber tractography is a sophisticated method, which can be used to delineate individual fiber tracts from diffusion images. The main process of study, uses the “DSI-Studio” software tools for,

i. Complete preprocessing,
ii. Fiber tracking and
iii. Analysis.

The imaging data processing, helps to convert the pre-processed raw data to .src file format, which will be suitable for the further reconstruction process. The reconstruction of .src files achieved through software tools. It converts the .src imaging data to .fib file format. Only the .fib files are compatible for fiber tracking. To delineate individual fiber tracts from reconstructed diffusion images (.fib file), using a DSI studio software tools.

**Fig. S1.**
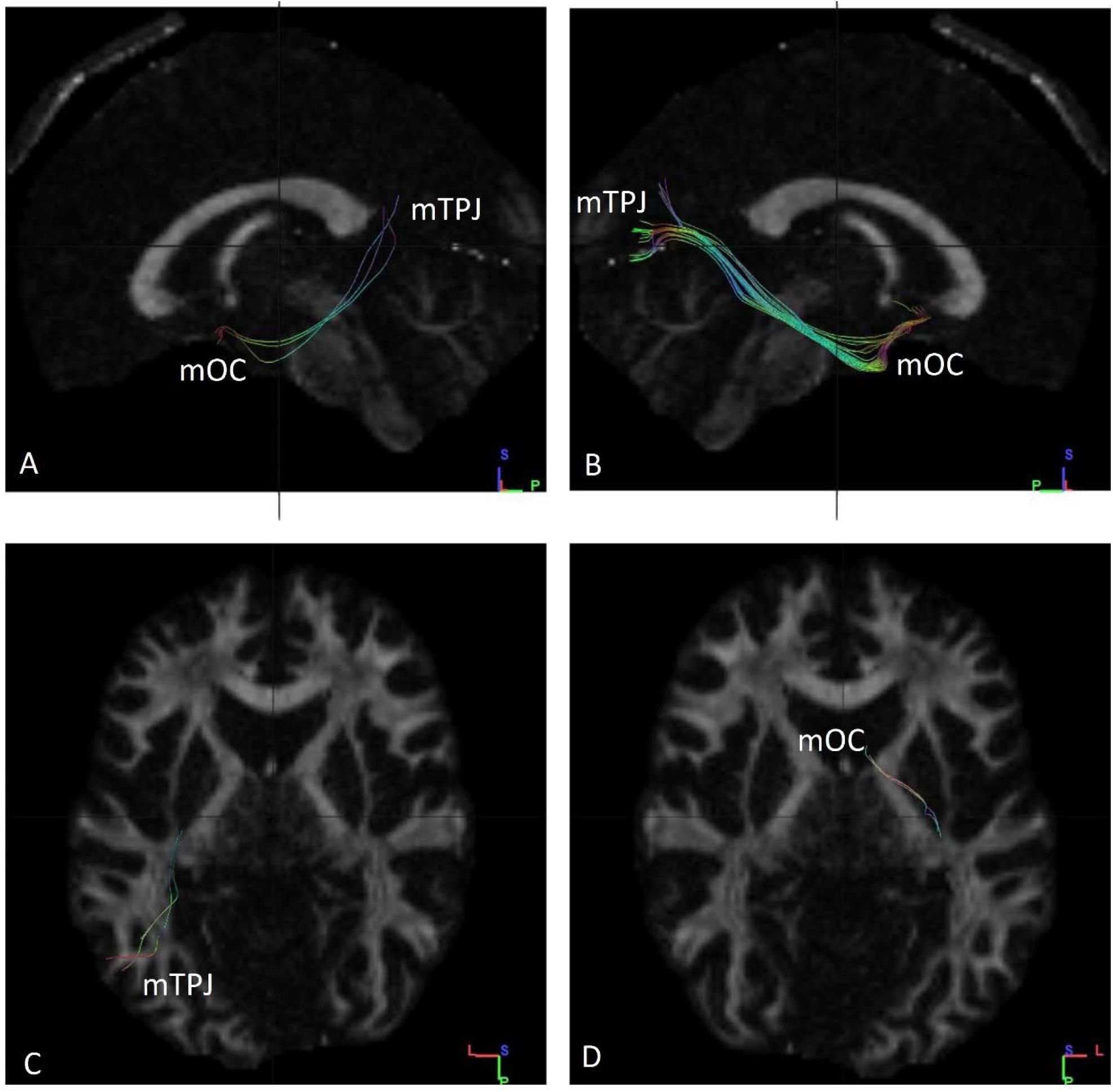
Fibres traced from mOC to mTPJ in subject no. 1. **(A)** Tracts observed from left sagittal view. **(B)** Tracts observed from right sagittal view. **(C)** Tracts ending in mTPJ as observed from superior axial view. **(D)** Tracts observed from inferior axial view. Some of the fibres are contralateral in nature. mTPJ – mergerd Temporoparietal Junction mOC – merged Olfactory Cortex

**Fig. S2.**
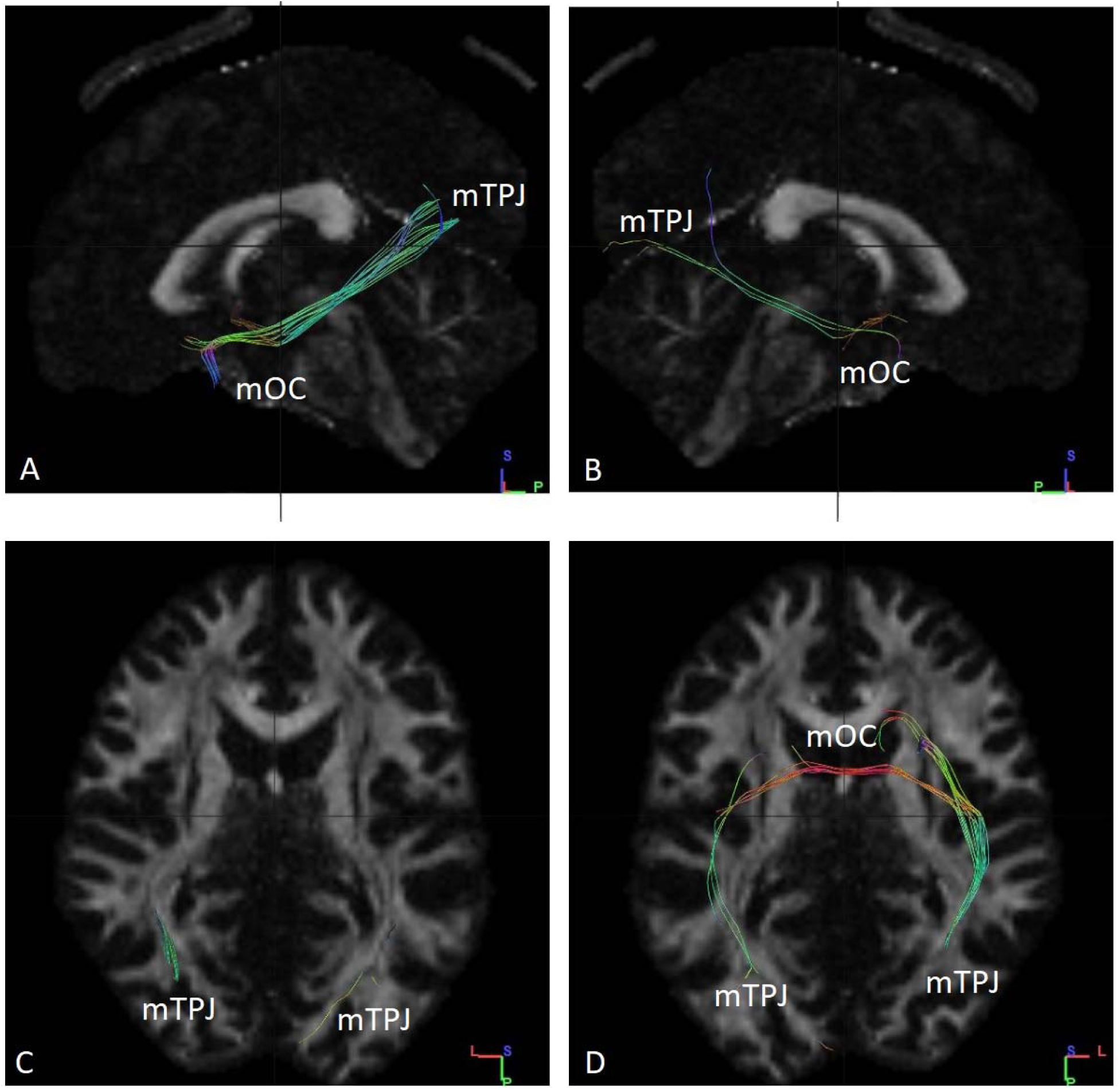
Fibres traced from mOC to mTPJ in subject no. 2. **(A)** Tracts observed from left sagittal view. **(B)** Tracts observed from right sagittal view. **(C)** Tracts observed from superior axial view. **(D)** Tracts ending in mTPJ as observed from an inferior axial view. Some of the fibres are contralateral in nature. mTPJ – mergerd Temporoparietal Junction mOC – merged Olfactory Cortex

**Fig. S3.**
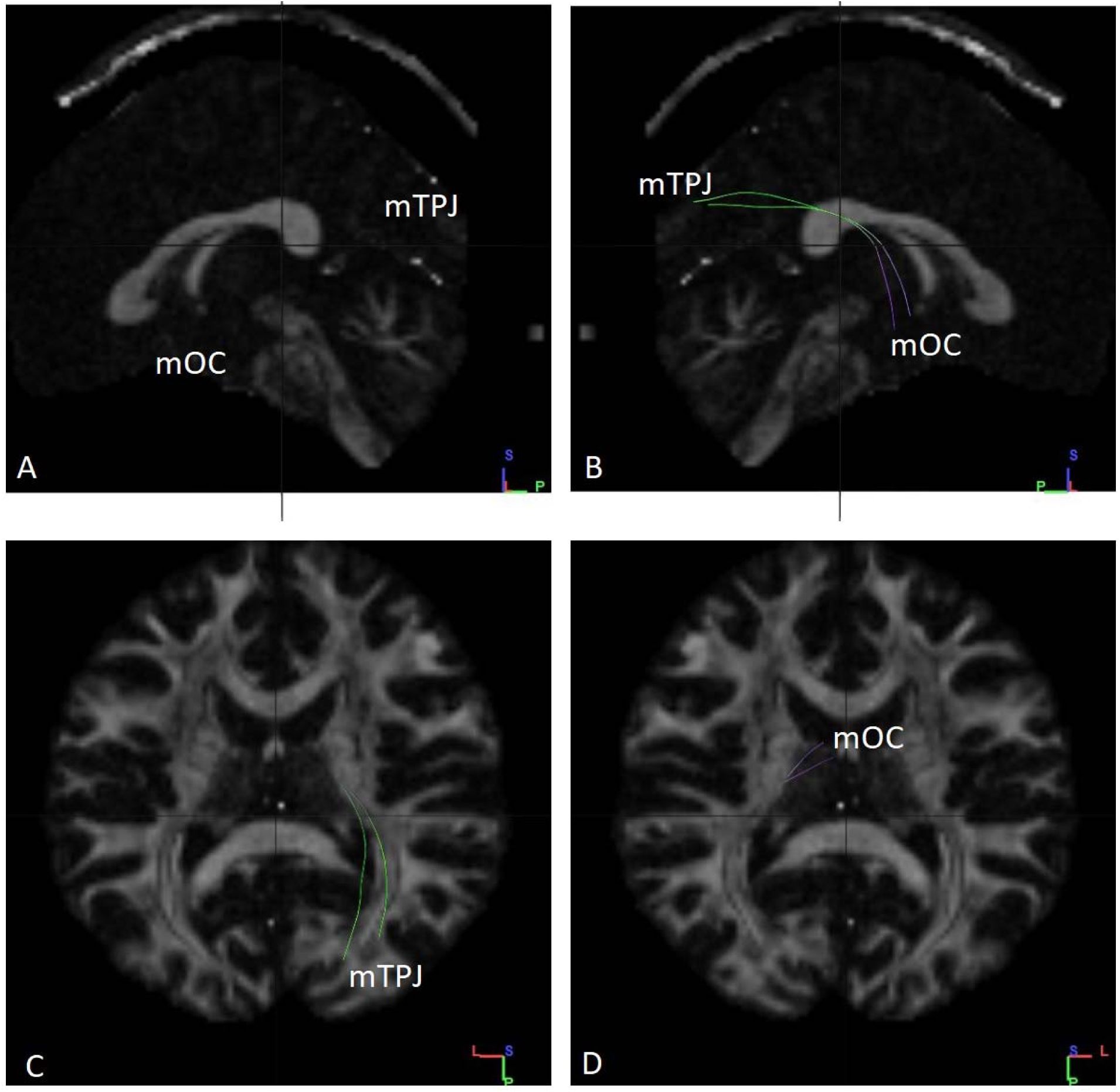
Fibres traced from mOC to mTPJ in subject no. 3. **(A)** No tracts observed from left sagittal view. **(B)** Tracts observed from right sagittal view. **(C)** Tracts ending in mTPJ as observed from the superior axial view. Tracts ending in mTPJ are ipsilateral in nature **(D)** Tracts observed from an inferior axial view. mTPJ – mergerd Temporoparietal Junction mOC – merged Olfactory Cortex

**Fig. S4.**
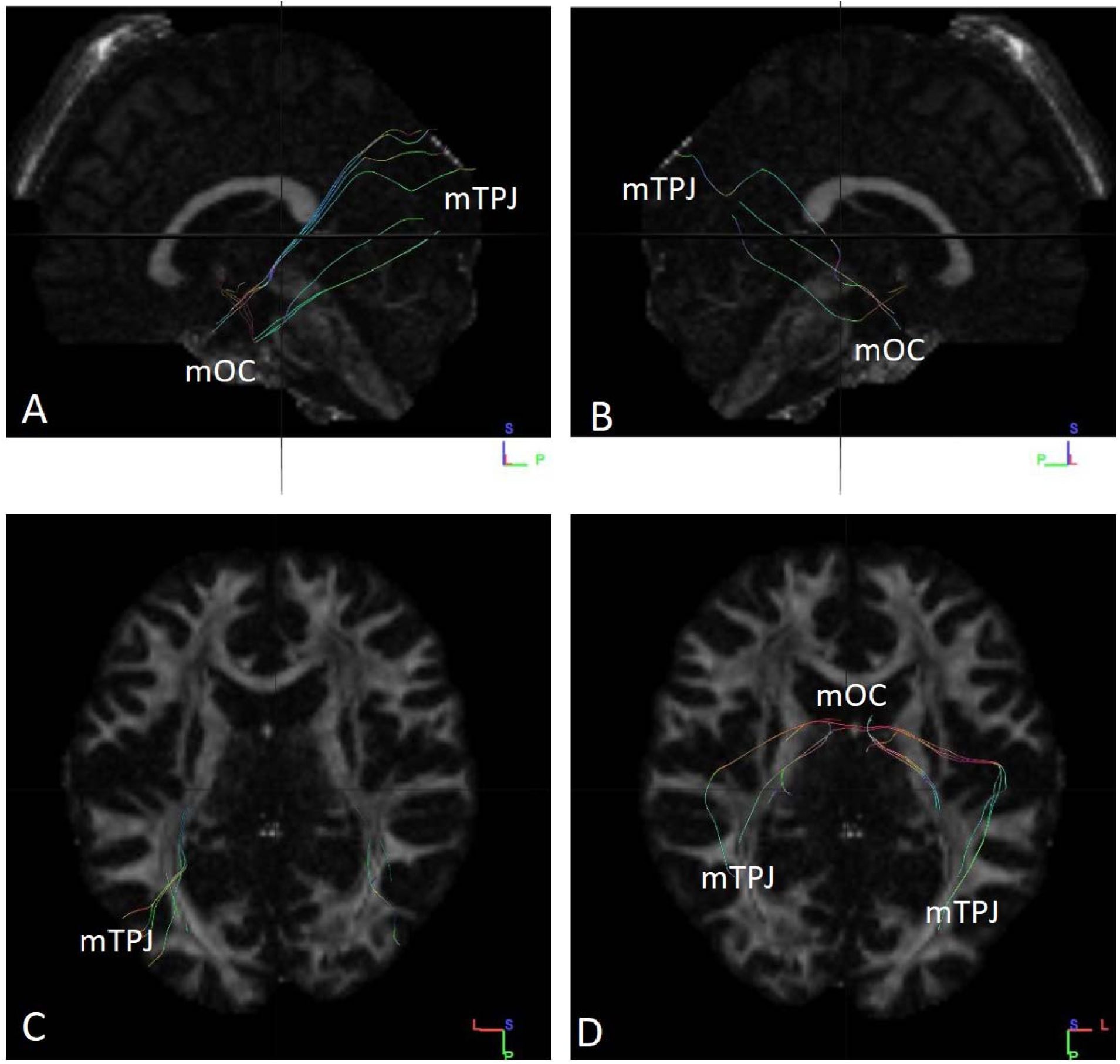
Fibres traced from mOC to mTPJ in subject no. 4. **(A)** Tracts observed from left sagittal view. **(B)** Tracts observed from right sagittal view. **(C)** Tracts ending in mTPJ as observed from superior axial view. **(D)** Tracts observed from an inferior axial view. Few of the fibres ending in the left side of the mTPJ arise from the right side of mOC. mTPJ – mergerd Temporoparietal Junction mOC – merged Olfactory Cortex

**Fig. S5.**
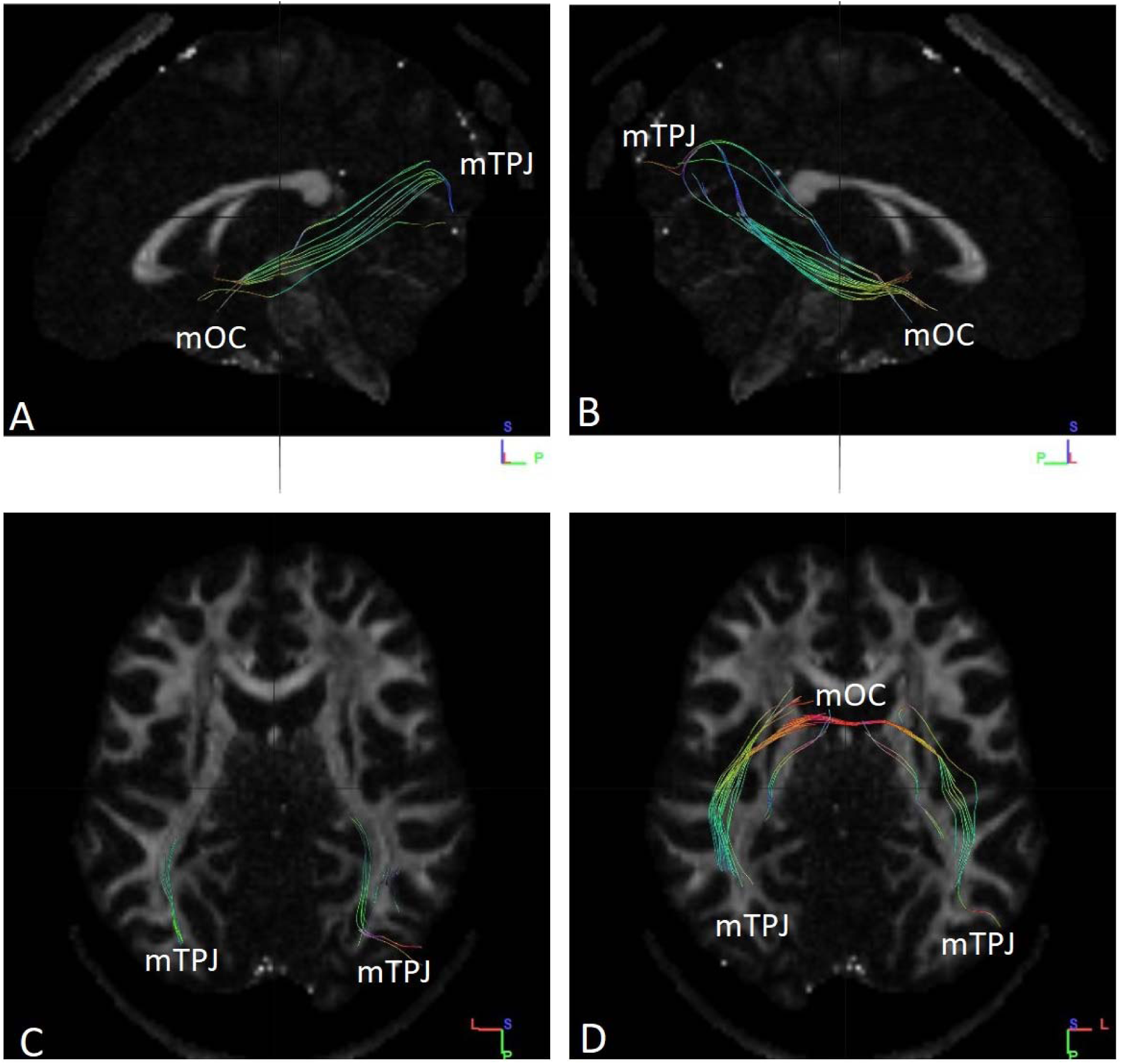
Fibres traced from mOC to mTPJ in subject no. 5. **(A)** Tracts observed from left sagittal view. **(B)** Tracts observed from right sagittal view. **(C)** Tracts ending in mTPJ as observed from superior axial view. **(D)** Tracts observed from an inferior axial view. Few of the fibres ending in the left side of the mTPJ arise from the right side of mOC. mTPJ – mergerd Temporoparietal Junction mOC – merged Olfactory Cortex

**Fig. S6.**
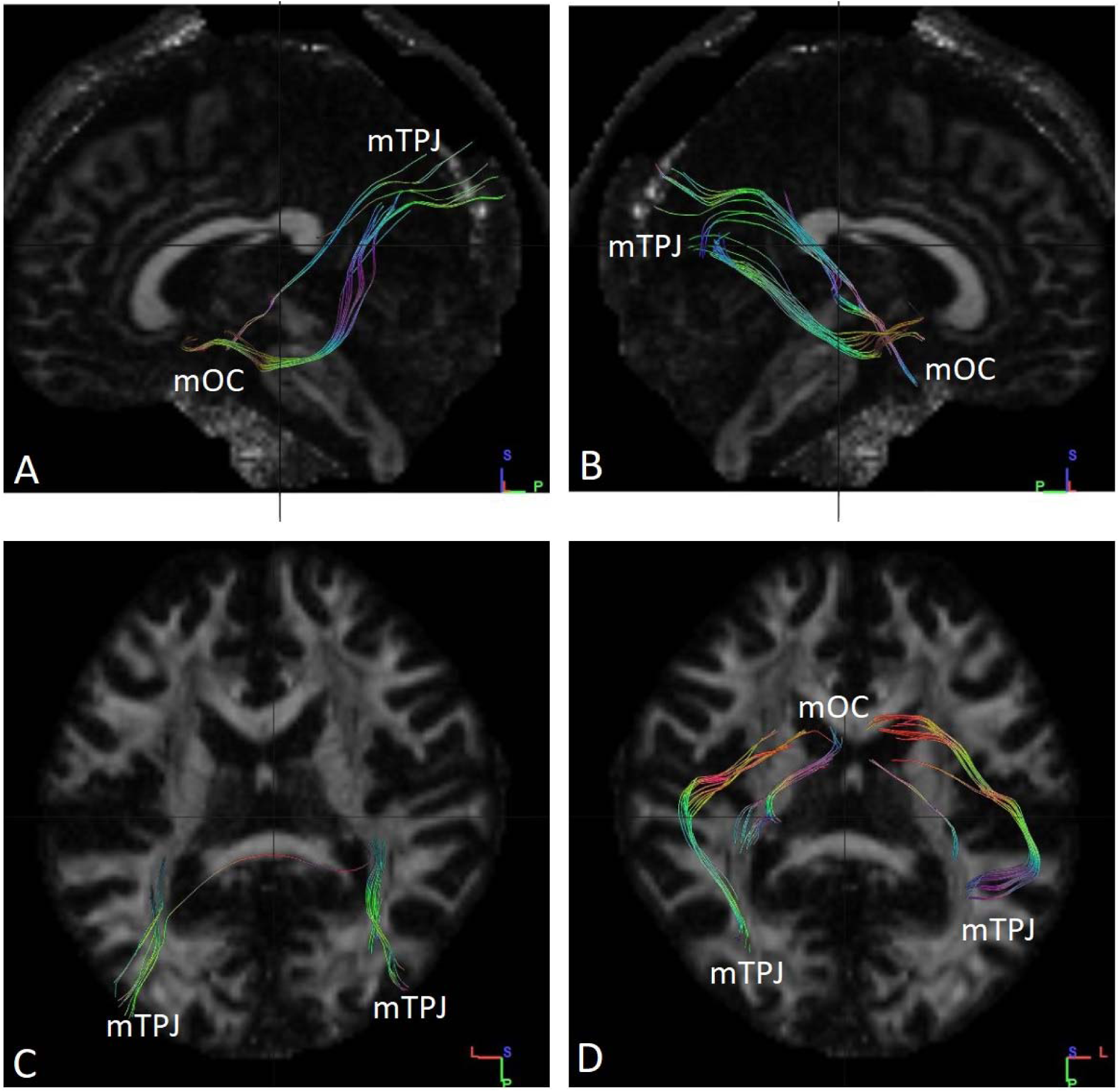
Fibres traced from mOC to mTPJ in subject no. 6. **(A)** Tracts observed from left sagittal view. **(B)** Tracts observed from right sagittal view. **(C)** Tracts ending in mTPJ as observed from superior axial view. **(D)** Tracts observed from an inferior axial view. Few of the fibres ending in the left side of the mTPJ arise from the right side of mOC. mTPJ – mergerd Temporoparietal Junction mOC – merged Olfactory Cortex

**Fig. S7.**
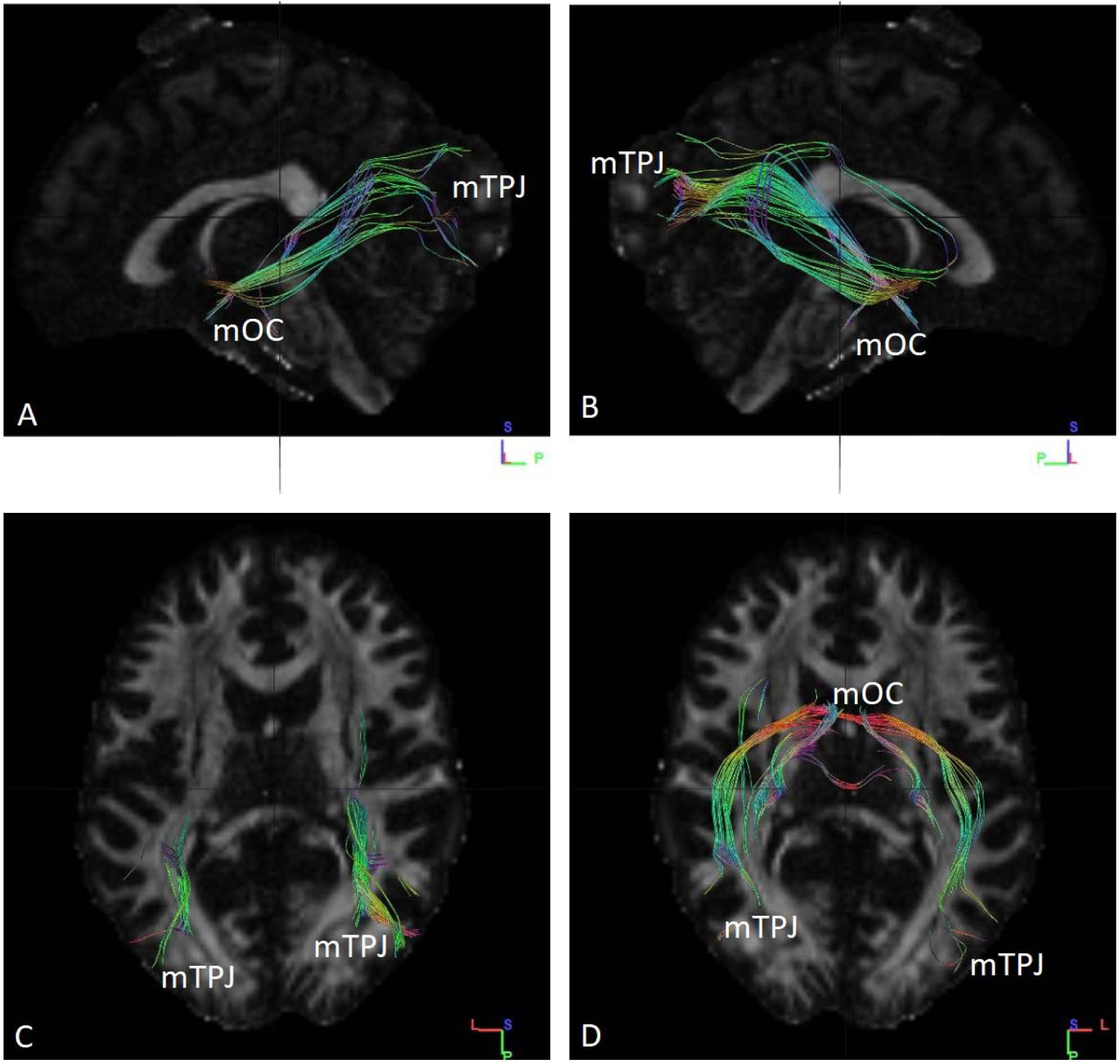
Fibres traced from mOC to mTPJ in subject no. 7. **(A)** Tracts observed from left sagittal view. **(B)** Tracts observed from right sagittal view. **(C)** Tracts ending in mTPJ observed from a superior axial view. **(D)** Tracts ending in mTPJ as observed from inferior axial view. Some of the fibres are contralateral in nature, rising from both right and left sife of the mOC. mTPJ – mergerd Temporoparietal Junction mOC – merged Olfactory Cortex

**Fig. S8.**
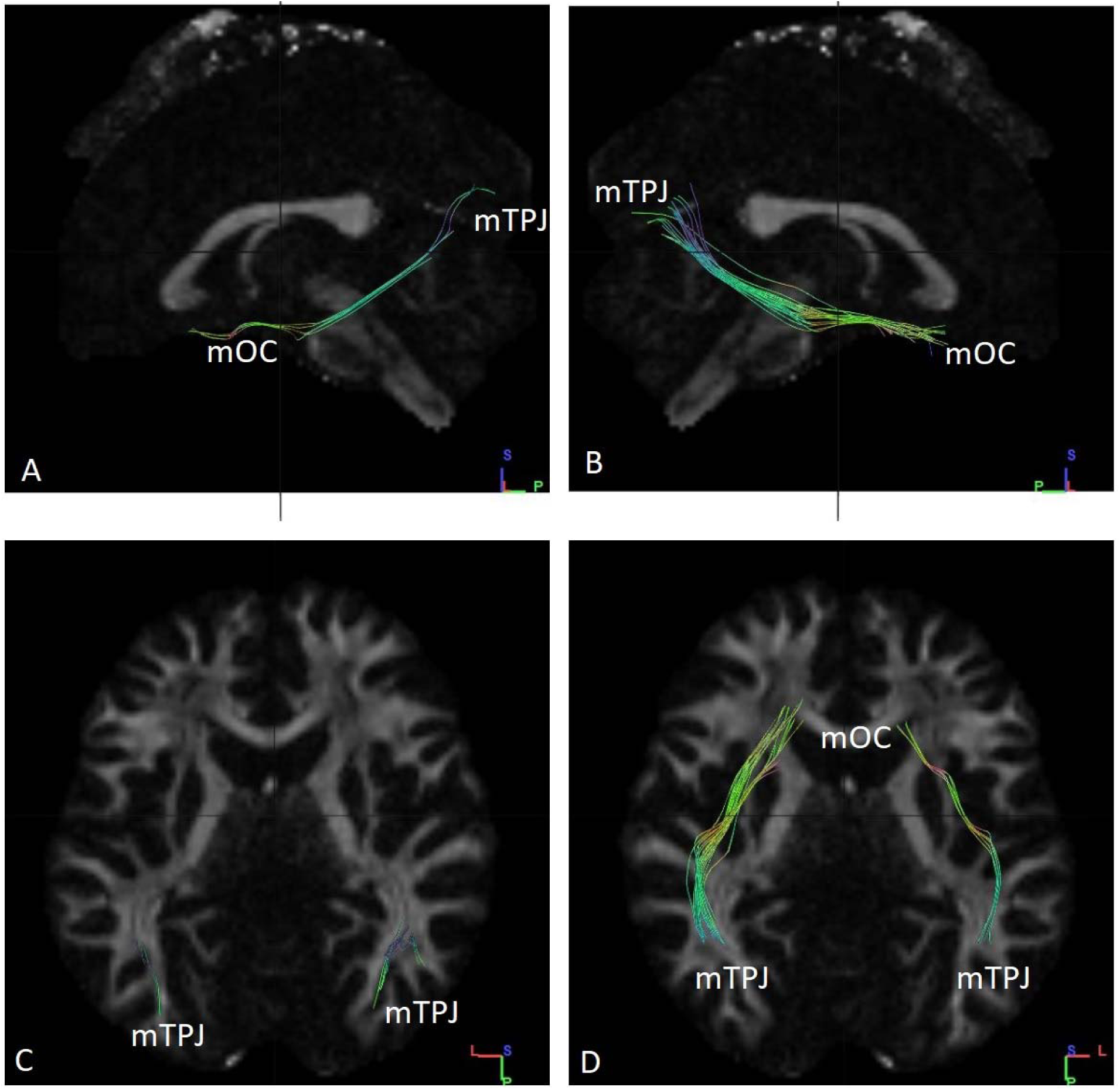
Fibres traced from mOC to mTPJ in subject no. 8. **(A)** Tracts observed from left sagittal view. **(B)** Tracts observed from right sagittal view. **(C)** Tracts ending in mTPJ observed from a superior axial view. **(D)** Tracts ending in mTPJ as observed from inferior axial view. All of the fibres are ipsilateral in nature. mTPJ – mergerd Temporoparietal Junction mOC – merged Olfactory Cortex

**Fig. S9.**
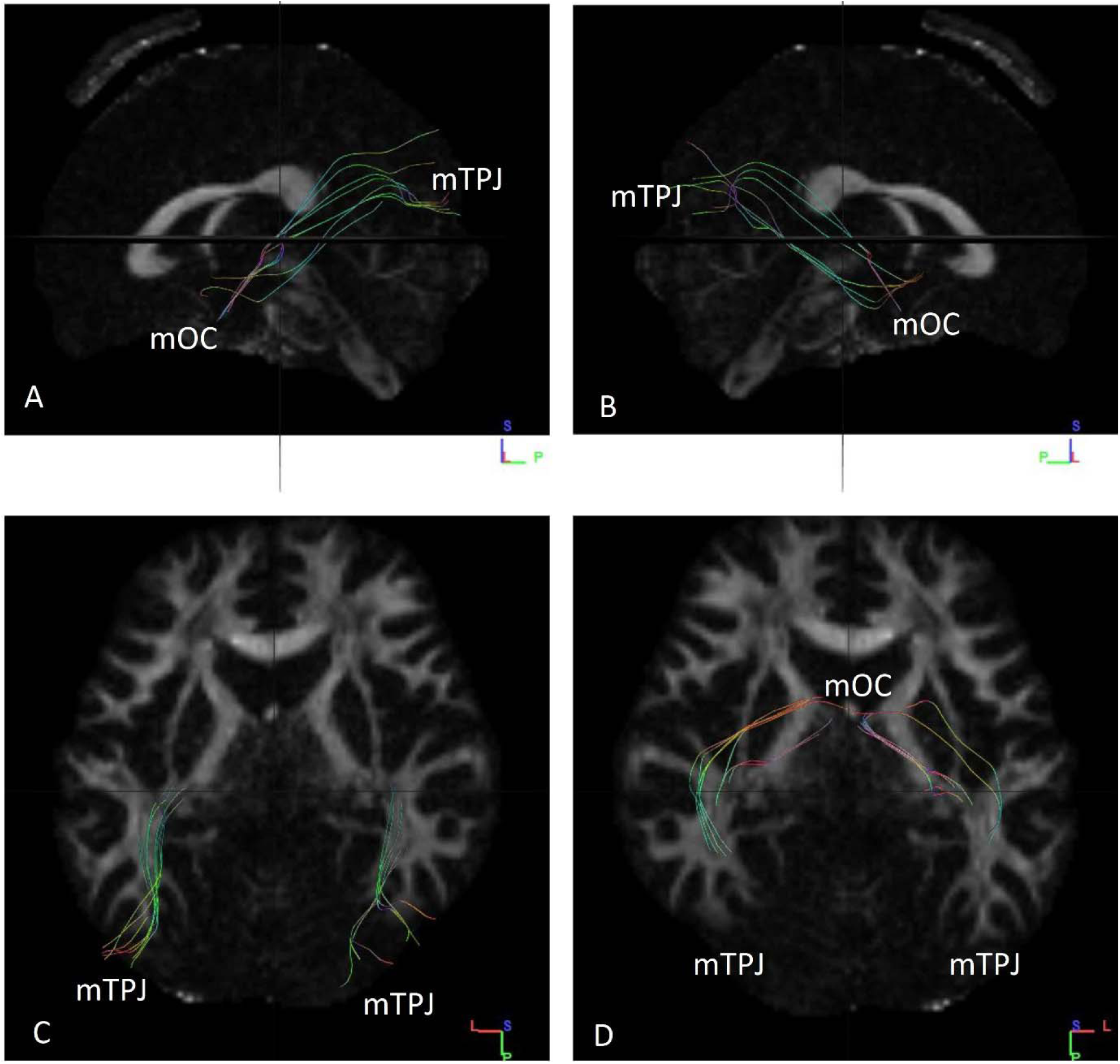
Fibres traced from mOC to mTPJ in subject no. 9. **(A)** Tracts observed from left sagittal view. **(B)** Tracts observed from right sagittal view. **(C)** Tracts ending in mTPJ observed from a superior axial view. **(D)** Tracts ending in mTPJ as observed from inferior axial view. All of the fibres are ipsilateral in nature. mTPJ – mergerd Temporoparietal Junction mOC – merged Olfactory Cortex

**Fig. S10.**
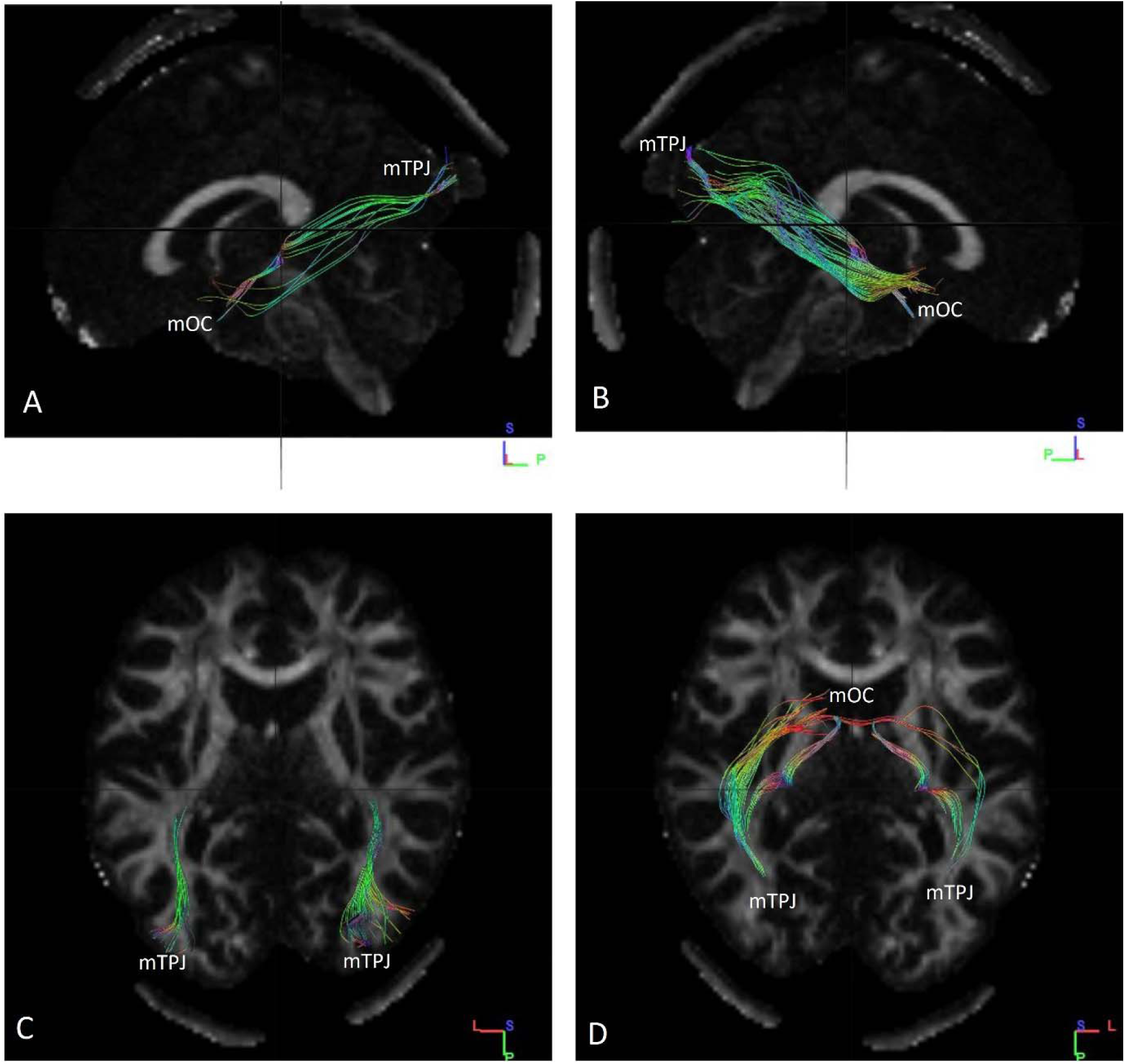
Fibres traced from mOC to mTPJ in subject no. 10. **(A)** Tracts observed from left sagittal view. **(B)** Tracts observed from right sagittal view. **(C)** Tracts ending in mTPJ observed from a superior axial view. **(D)** Tracts ending in mTPJ as observed from an inferior axial view. All of the fibres are ipsilateral in nature. mTPJ – mergerd Temporoparietal Junction mOC – merged Olfactory Cortex

**Fig. S11.**
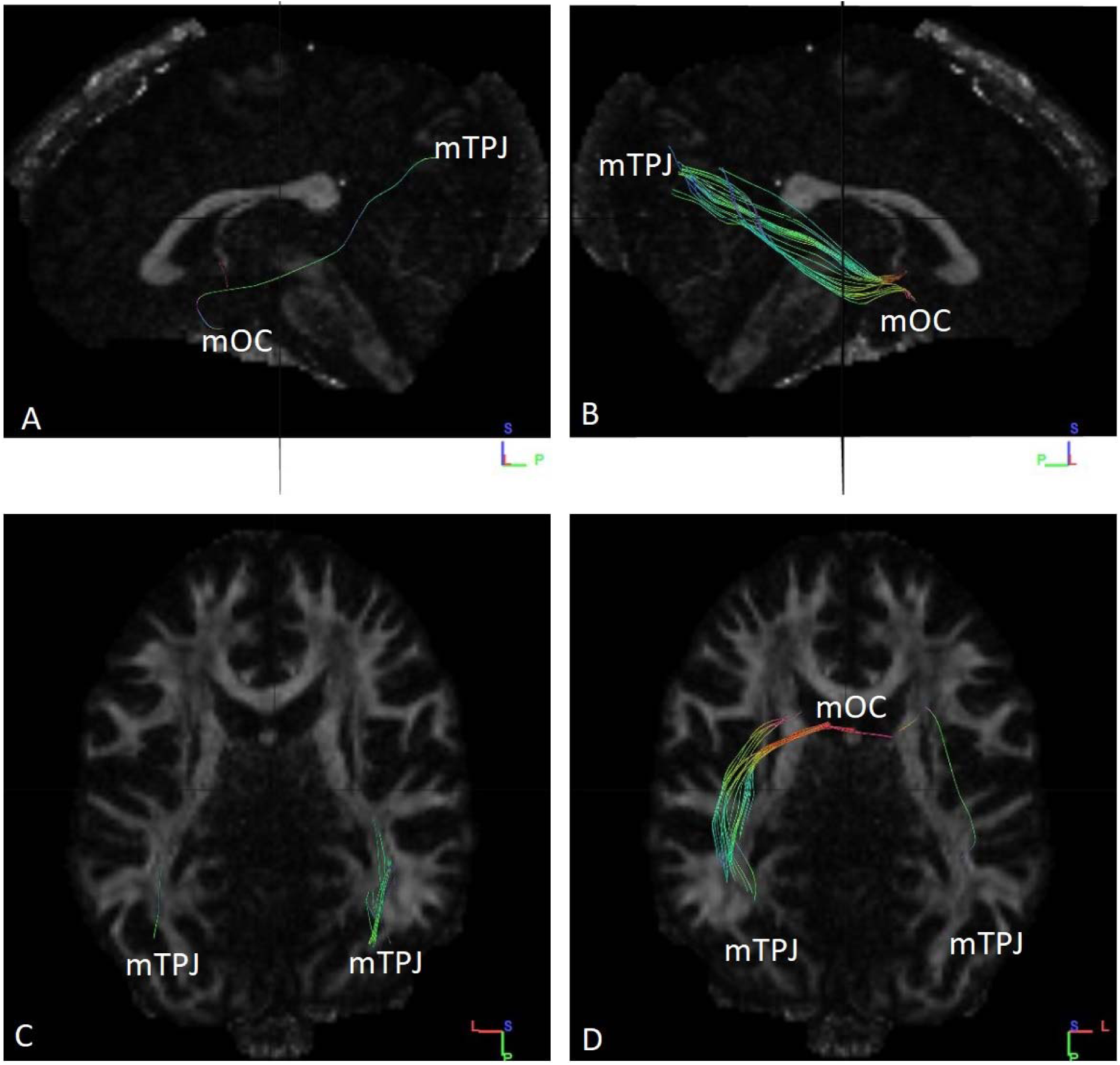
Fibres traced from mOC to mTPJ in subject no. 11. **(A)** Tracts observed from left sagittal view. **(B)** Tracts observed from right sagittal view. **(C)** Tracts ending in mTPJ as observed from superior axial view. **(D)** Tracts ending in mTPJ observed from an inferior axial view. Some of the fibres ending in the right side of mTPJ arise from the left side of mOC. mTPJ – mergerd Temporoparietal Junction mOC – merged Olfactory Cortex

**Fig. S12.**
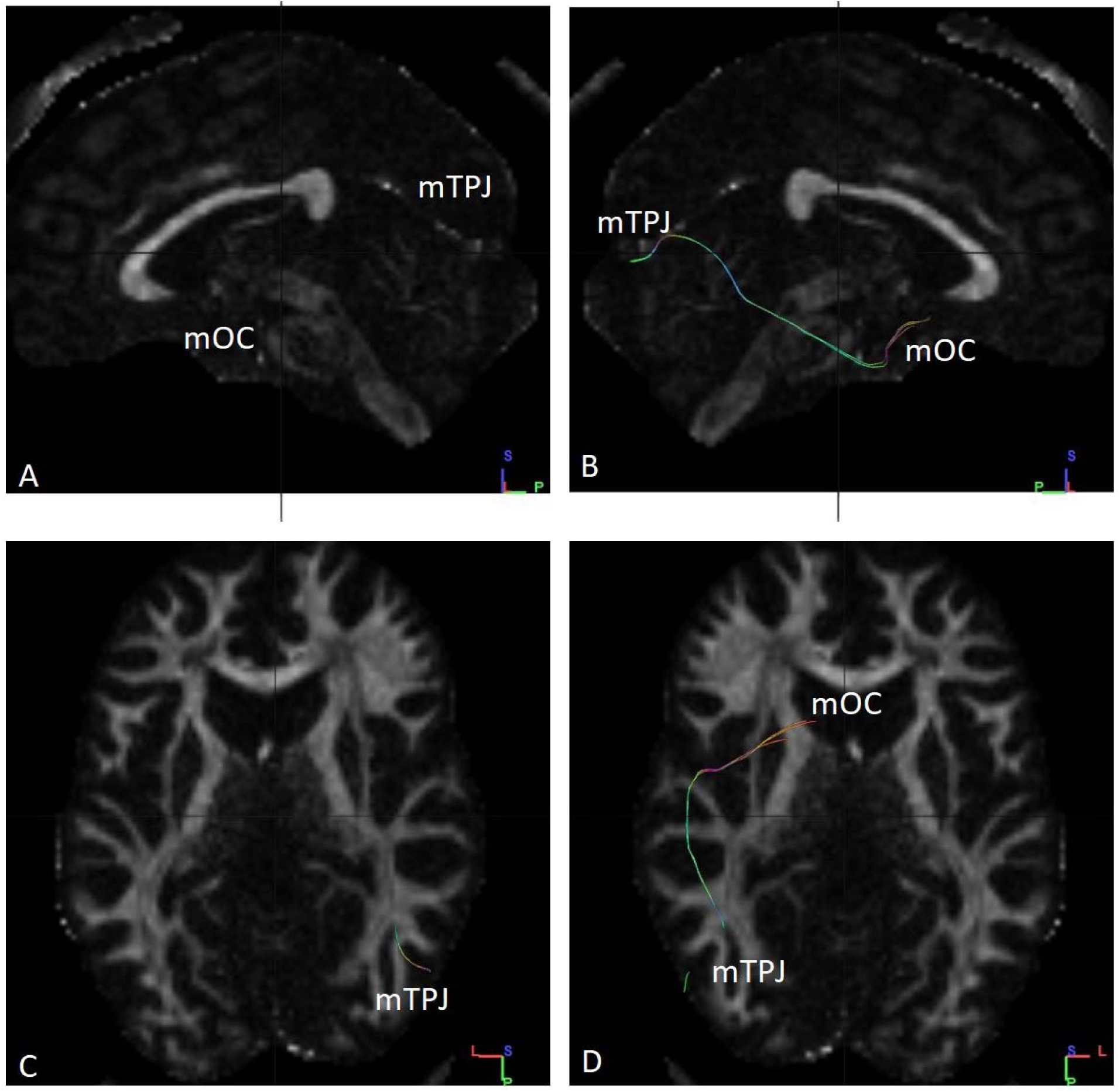
Fibres traced from mOC to mTPJ in subject no. 12. **(A)** No tracts were observed from left sagittal view. **(B)** Tracts observed from right sagittal view. **(C)** Tracts ending in mTPJ as observed from superior axial view. **(D)** Tracts ending in mTPJ observed from an inferior axial view. All the fibres seen are ipsilateral in nature. mTPJ – mergerd Temporoparietal Junction mOC – merged Olfactory Cortex

**Fig. S13.**
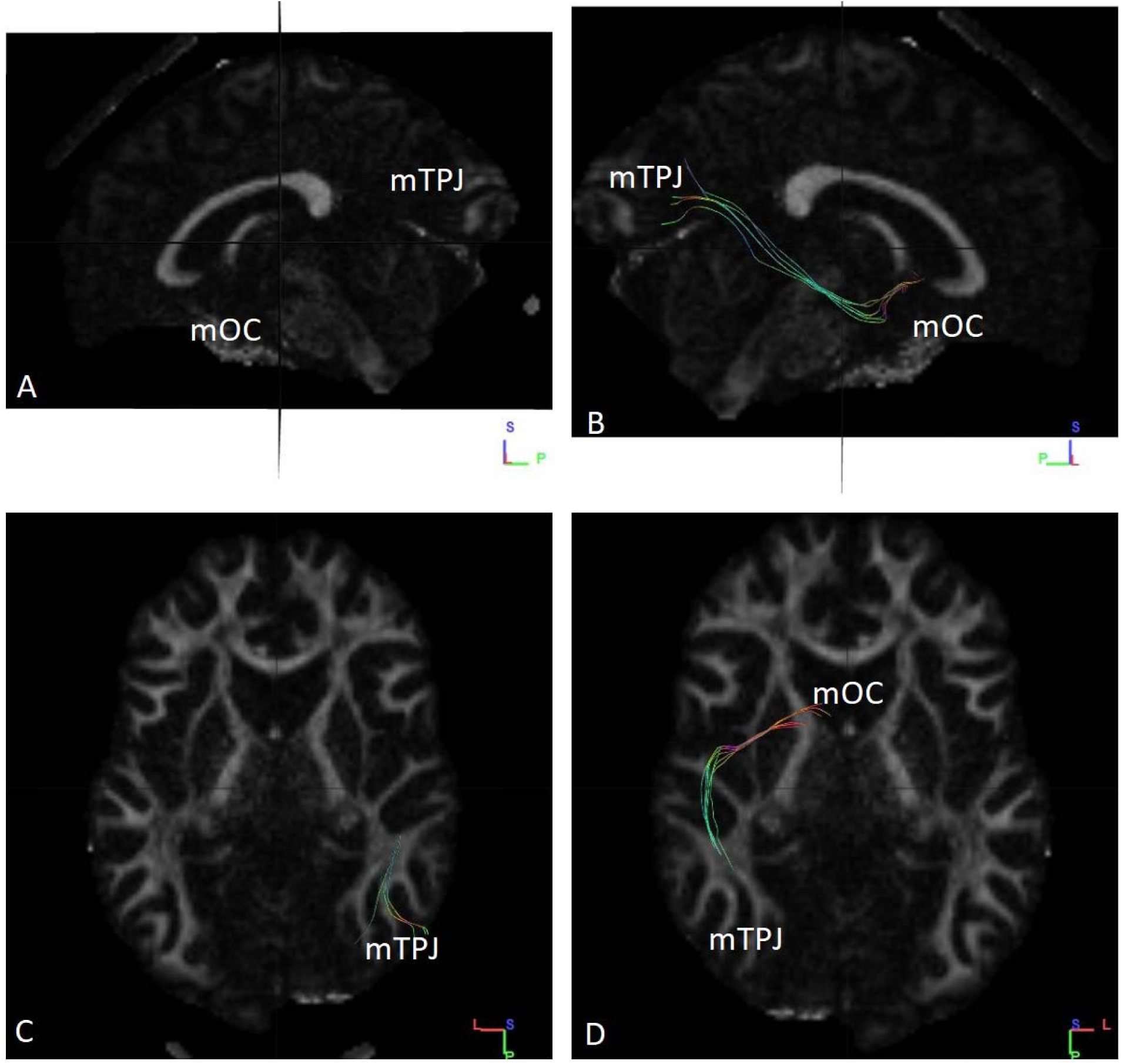
Fibres traced from mOC to mTPJ in subject no. 13. **(A)** No tracts observed from left sagittal view. **(B)** Tracts were observed from right sagittal view. **(C)** Tracts ending in mTPJ as observed from superior axial view. **(D)** Tracts ending in mTPJ observed from an inferior axial view. All the fibres seen are ipsilateral in nature. mTPJ – mergerd Temporoparietal Junction mOC – merged Olfactory Cortex

**Fig. S14.**
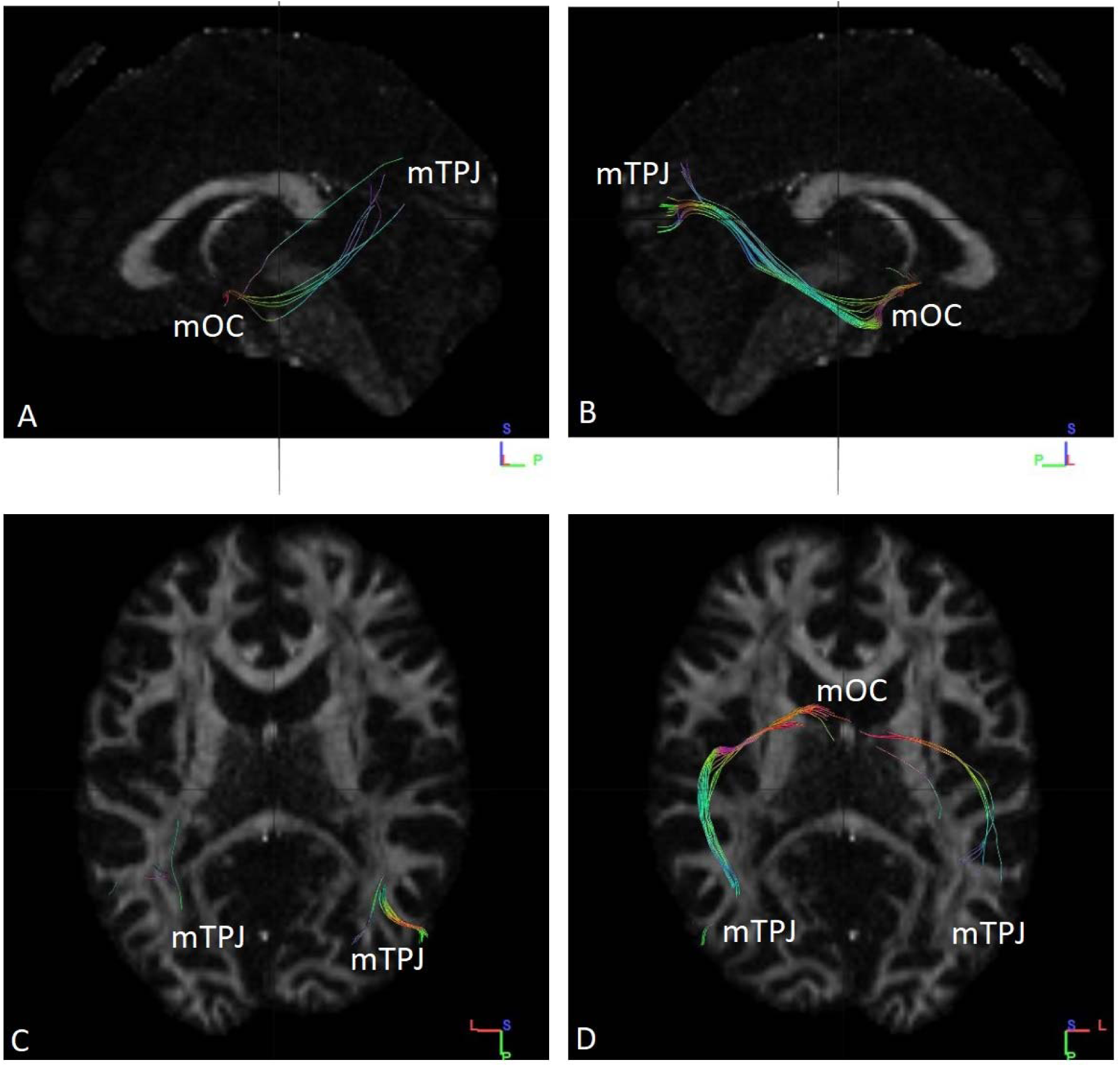
Fibres traced from mOC to mTPJ in subject no. 14. (A) Tracts observed from left sagittal view. (B) Tracts were observed from right sagittal view. (C) Tracts ending in mTPJ as observed from superior axial view. (D) Tracts ending in mTPJ observed from an inferior axial view. All the fibres seen are ipsilateral in nature. mTPJ – mergerd Temporoparietal Junction mOC – merged Olfactory Cortex

**Fig. S15.**
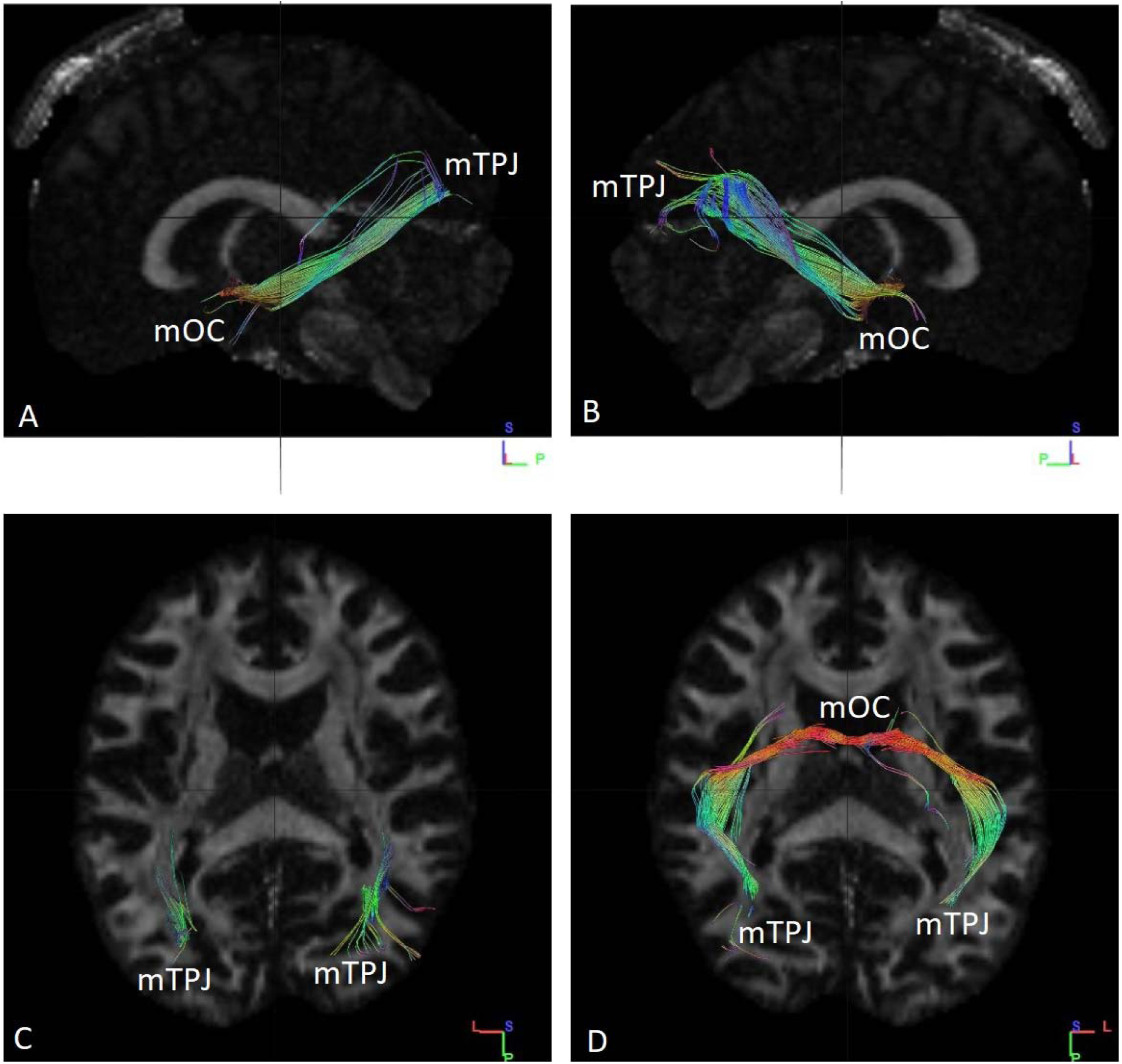
Fibres traced from mOC to mTPJ in subject no. 15. **(A)** Tracts were observed from left sagittal view. **(B)** Tracts observed from right sagittal view. **(C)** Tracts ending in mTPJ as observed from superior axial view. **(D)** Tracts ending in mTPJ observed from an inferior axial view. Some of the fibres ending in the right side of mTPJ arise from the left side of mOC. mTPJ – mergerd Temporoparietal Junction mOC – merged Olfactory Cortex

**Fig. S16.**
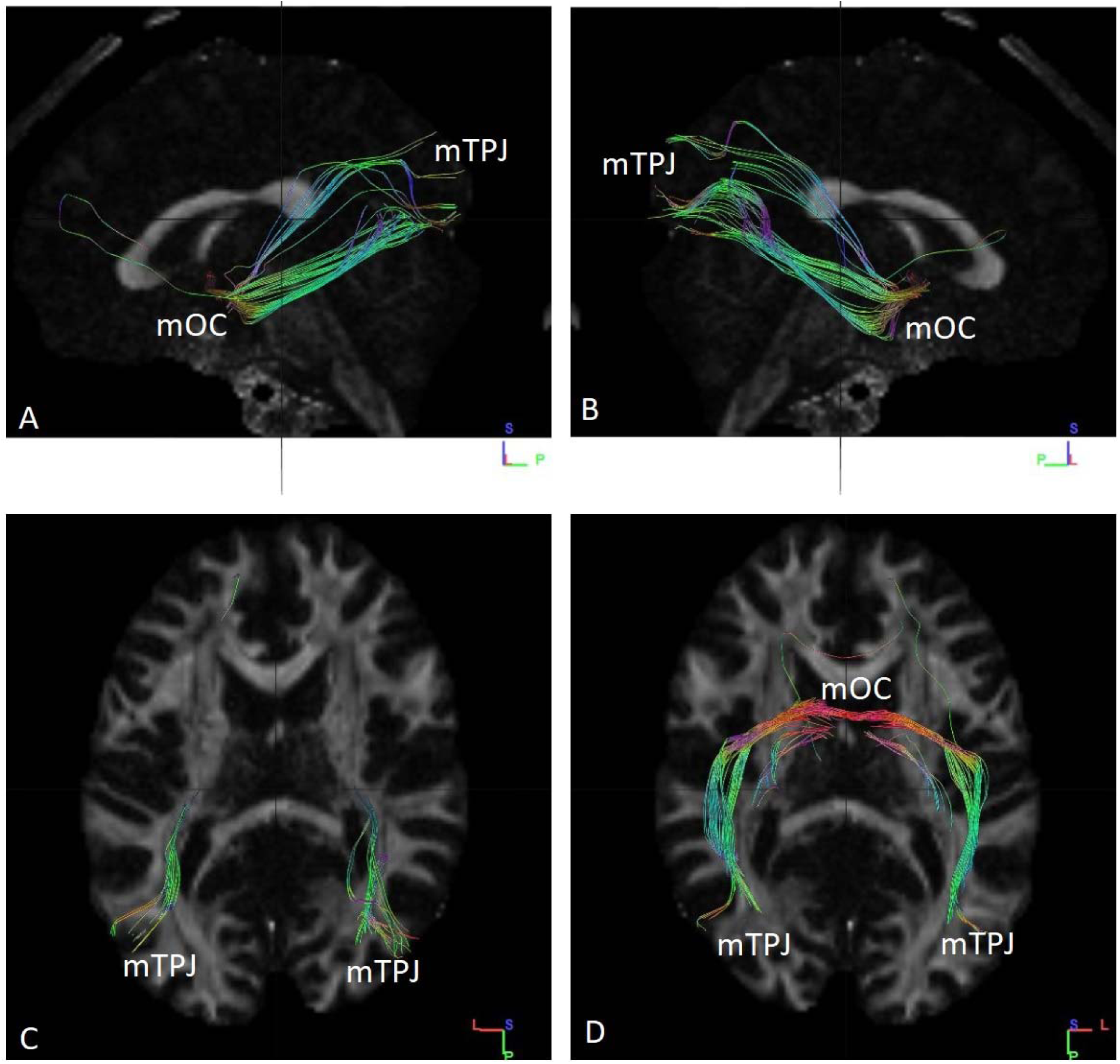
Fibres traced from mOC to mTPJ in subject no. 16. **(A)** Tracts were observed from left sagittal view**. (B)** Tracts observed from right sagittal view. **(C)** Tracts ending in mTPJ as observed from superior axial view. **(D)** Tracts ending in mTPJ observed from an inferior axial view. Some of the fibres ending in the mTPJ are contralateral in nature, rising from both right and left side of mOC. mTPJ – mergerd Temporoparietal Junction mOC – merged Olfactory Cortex

**Fig. S17.**
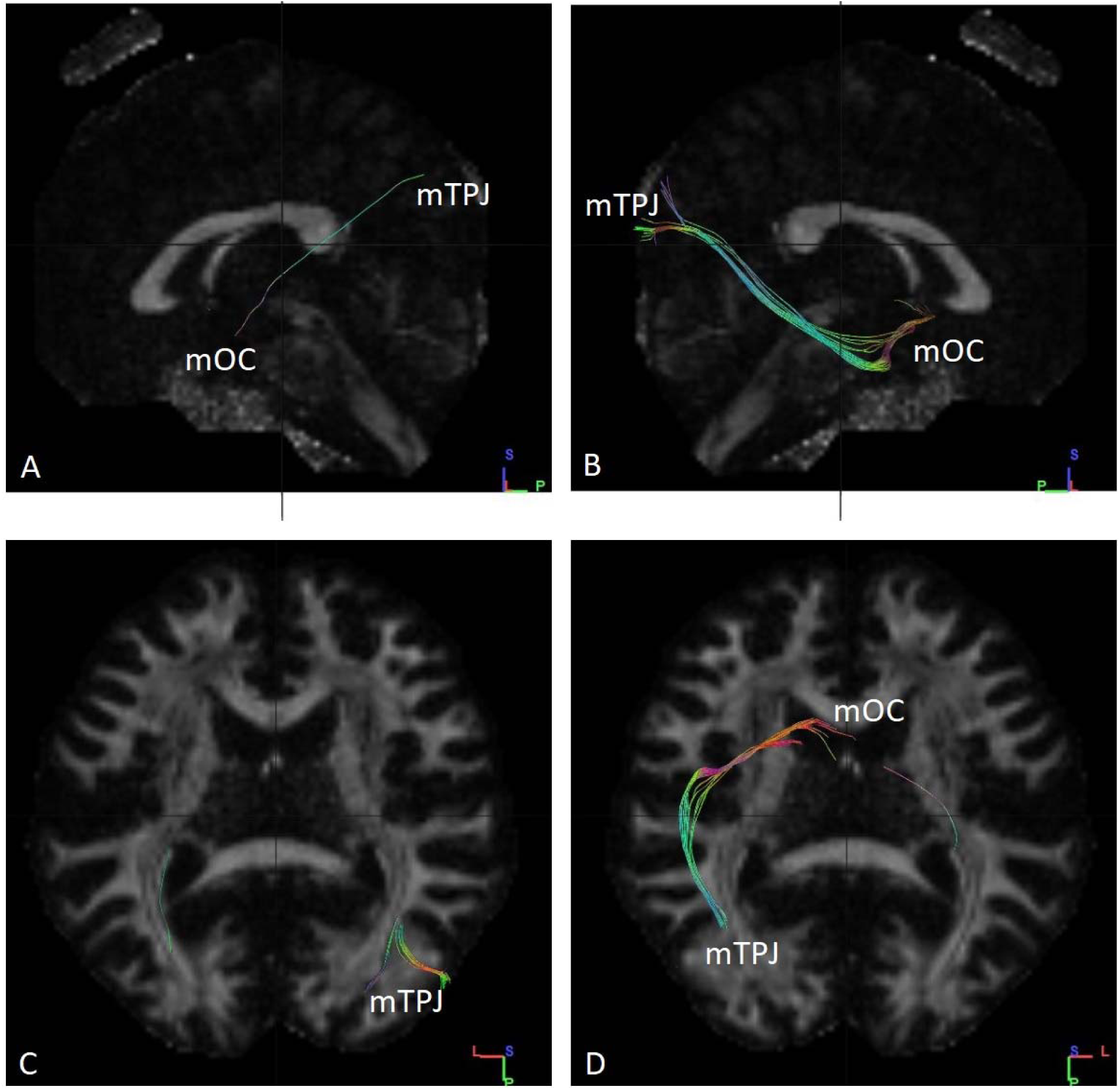
Fibres traced from mOC to mTPJ in subject no. 17. **(A)** No tracts were observed from left sagittal view. **(B)** Tracts observed from right sagittal view. **(C)** Tracts ending in mTPJ as observed from superior axial view. **(D)** Tracts ending in mTPJ observed from an inferior axial view. All the fibres are ipsilateral in nature. mTPJ – mergerd Temporoparietal Junction mOC – merged Olfactory Cortex

**Fig. S18.**
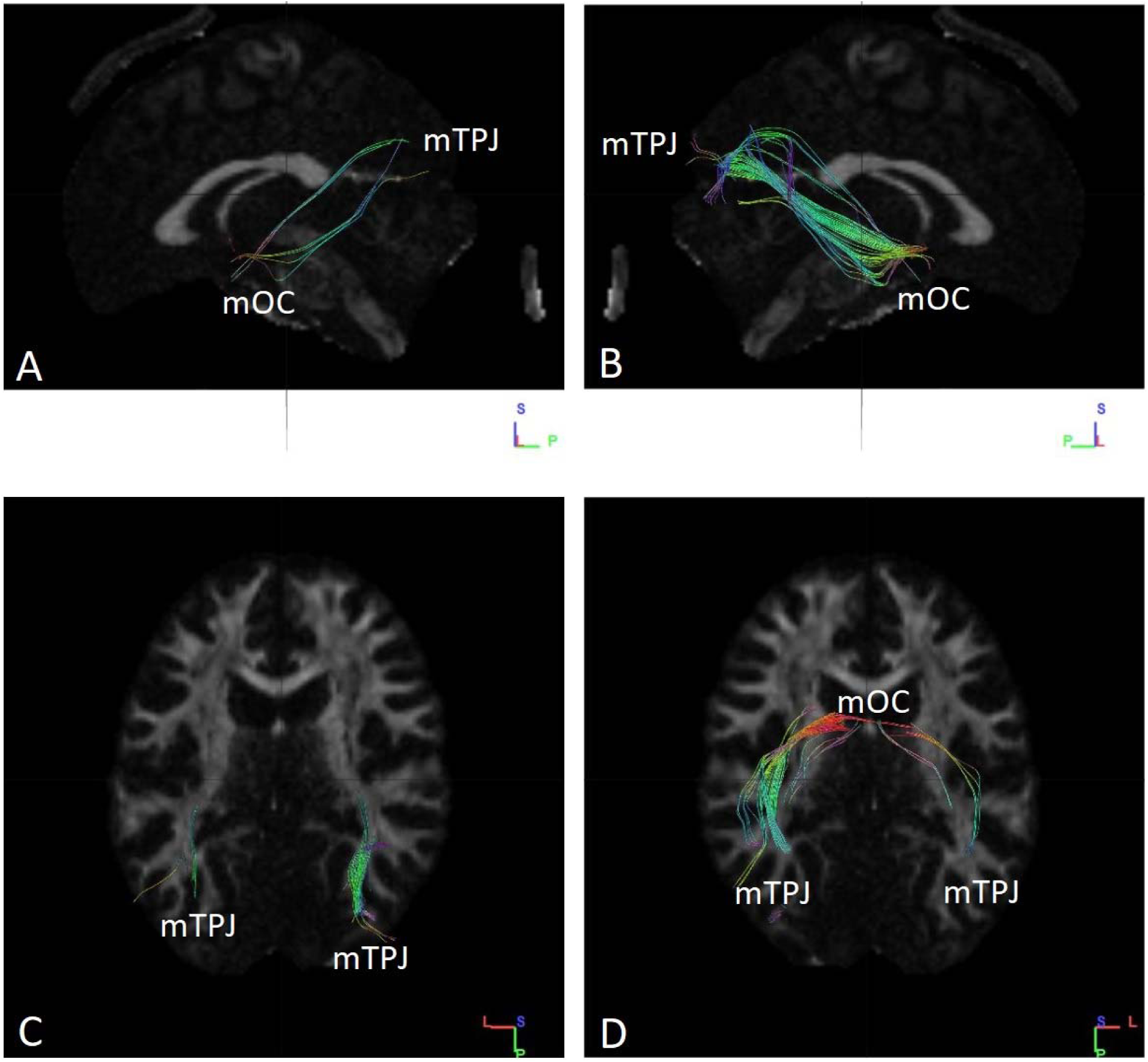
Fibres traced from mOC to mTPJ in subject no. 18. **(A)** Tracts observed from left sagittal view. **(B)** Tracts observed from right sagittal view. **(C)** Tracts ending in mTPJ as observed from superior axial view. **(D)** Tracts ending in mTPJ observed from an inferior axial view. Some of the fibres ending in the right side of mTPJ arise from the left side of mOC. mTPJ – mergerd Temporoparietal Junction mOC – merged Olfactory Cortex

**Fig. S19.**
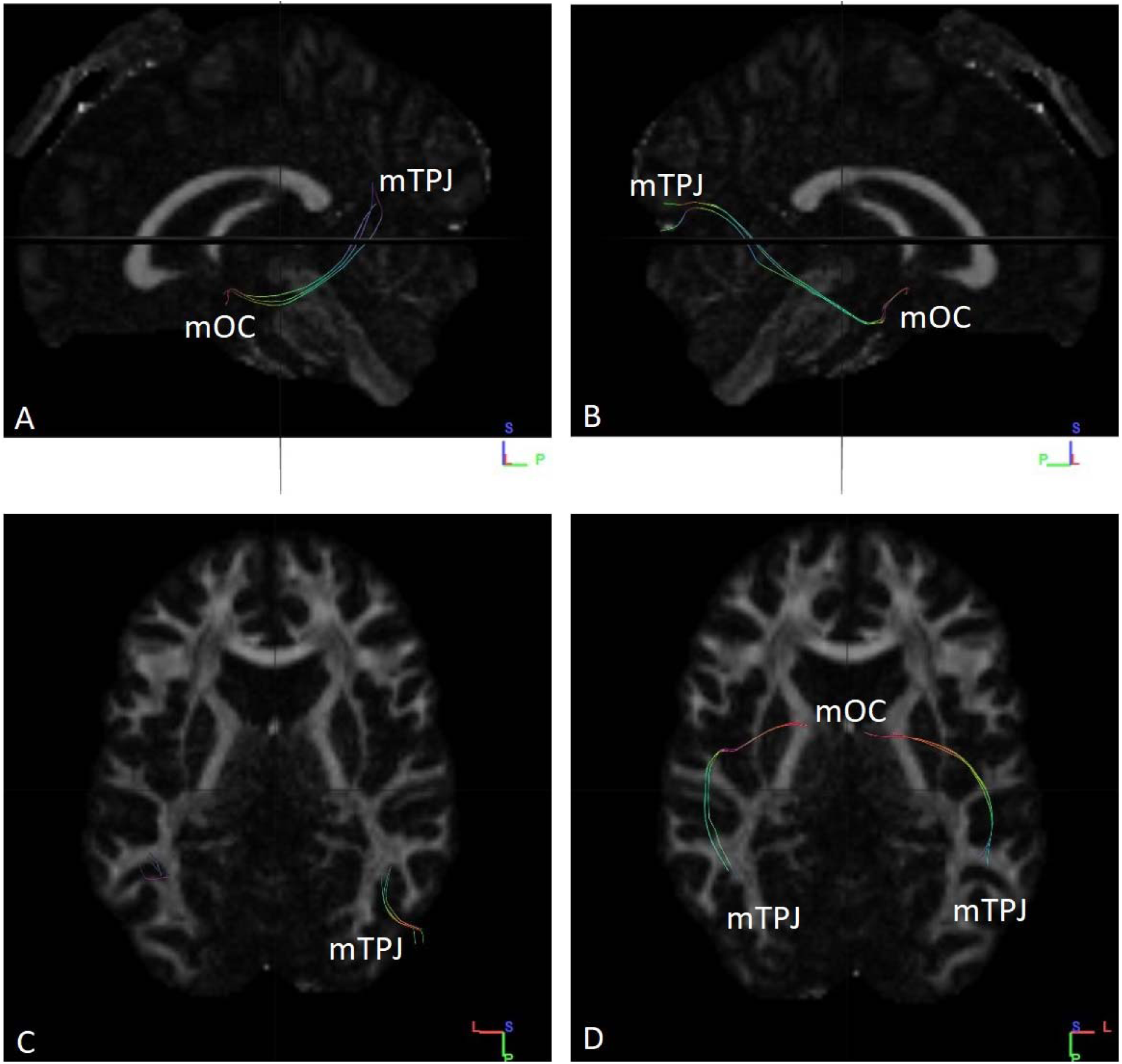
Fibres traced from mOC to mTPJ in subject no. 19. **(A)** Tracts observed from left sagittal view. **(B)** Tracts observed from right sagittal view. **(C)** Tracts ending in mTPJ as observed from superior axial view. **(D)** Tracts ending in mTPJ observed from an inferior axial view. All the fibres are ipsilateral in nature. mTPJ – mergerd Temporoparietal Junction mOC – merged Olfactory Cortex

**Fig. S20.**
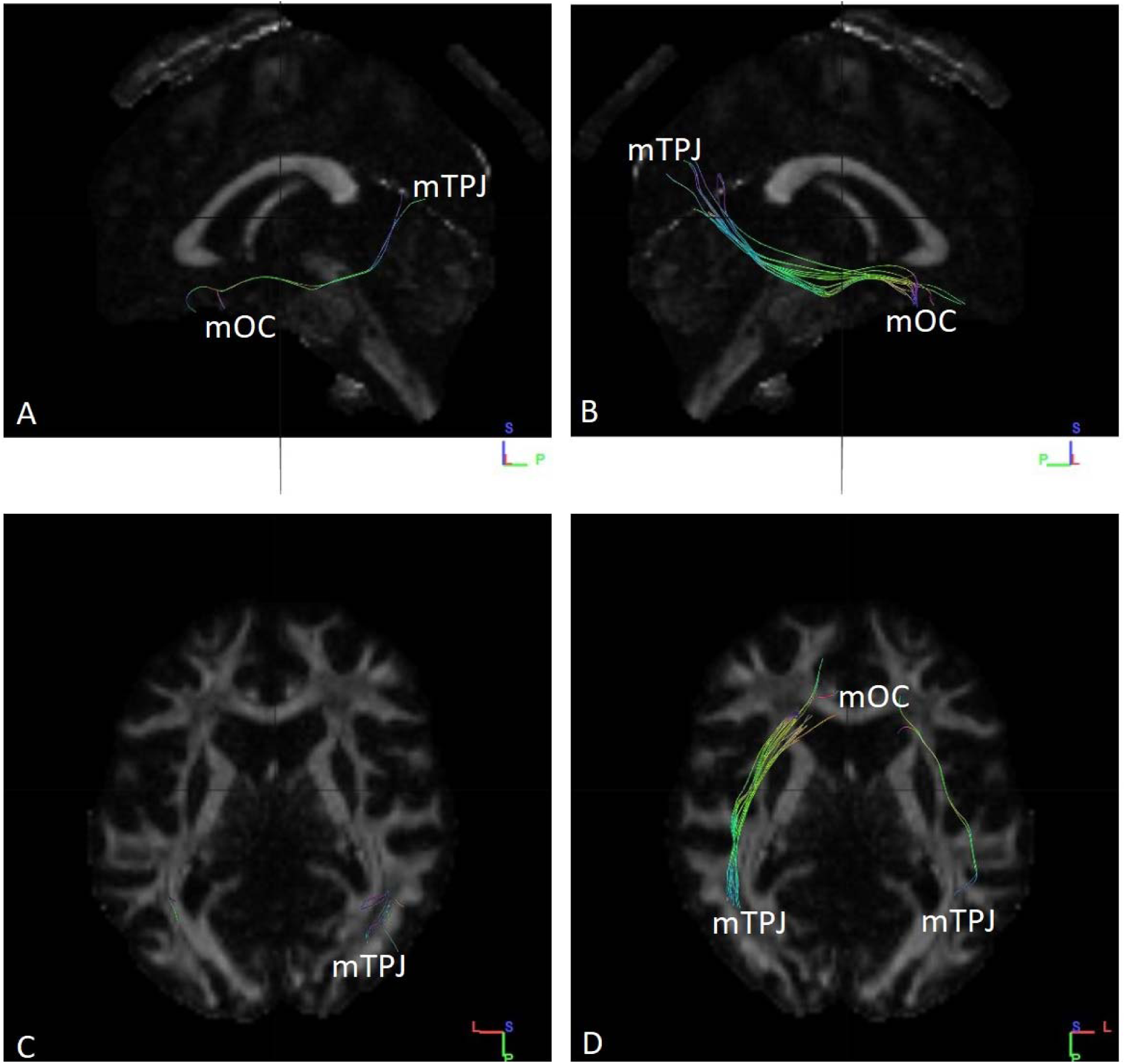
Fibres traced from mOC to mTPJ in subject no. 20. (A) Tracts observed from left sagittal view. (B) Tracts observed from right sagittal view. (C) Tracts ending in mTPJ as observed from superior axial view. (D) Tracts ending in mTPJ observed from an inferior axial view. All the fibres are ipsilateral in nature. mTPJ – mergerd Temporoparietal Junction mOC – merged Olfactory Cortex

**Fig. S21.**
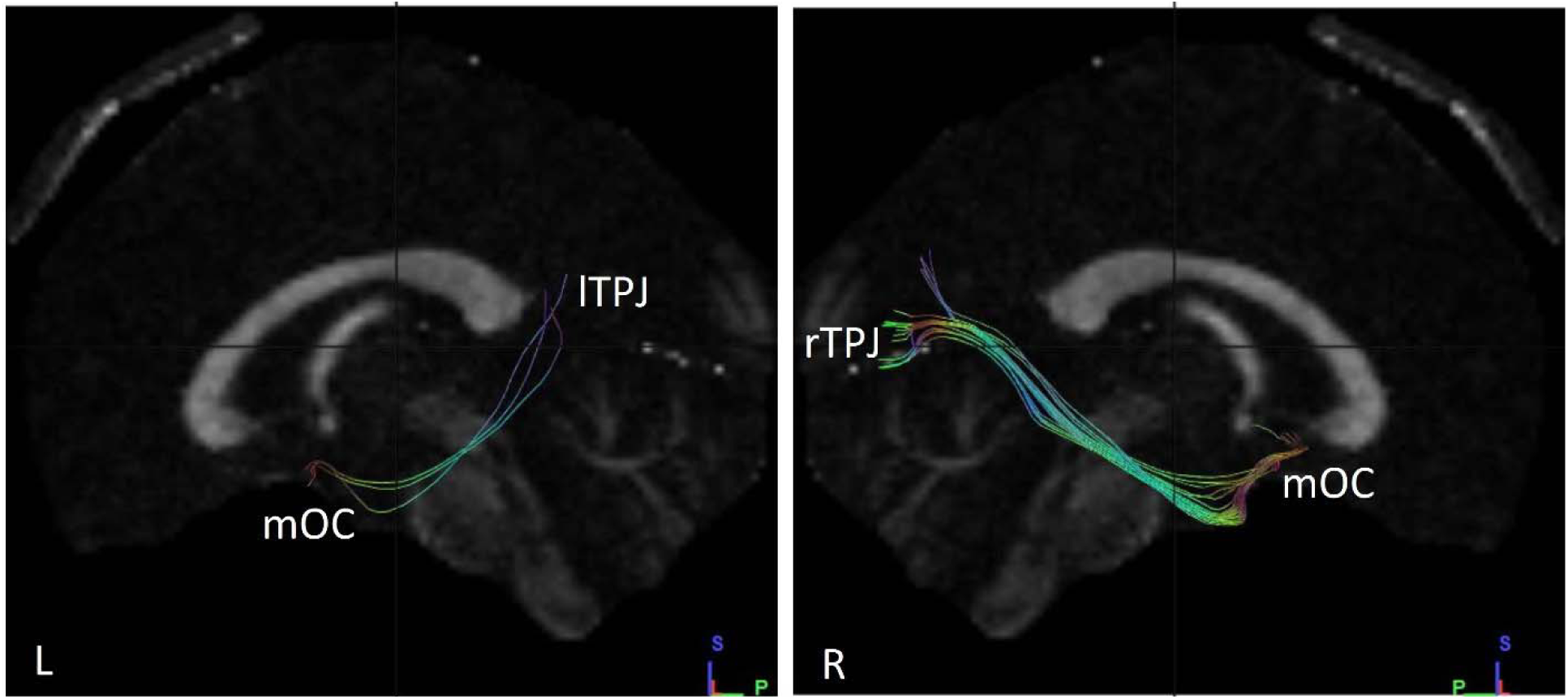
Fibres traced from mOC to lTPJ and rTPJ in subject no. 1. **(L)** Tracts observed from left sagittal view, when fibres traced from mOC to lTPJ. **(R)** Tracts observed from right sagittal view, when fibres traced from mOC to rTPJ. The number and volume of tracts are visibly low in the left hemisphere as compared to right. mOC – merged Olfactory Cortex lTPJ – left Temporoparietal Junction rTPJ – right Temporoparietal Junction

**Fig. S22.**
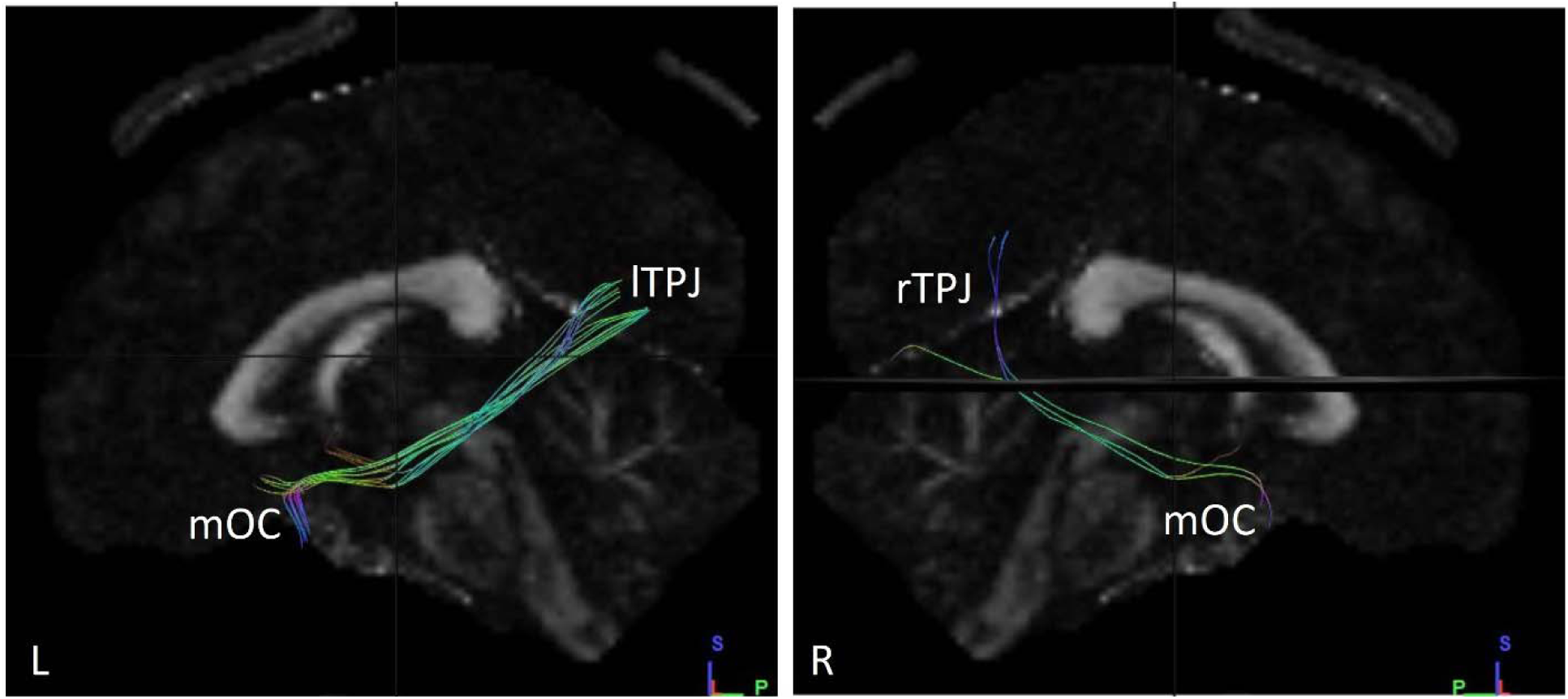
Fibres traced from mOC to lTPJ and rTPJ in subject no. 2. **(L)** Tracts observed from left sagittal view, when fibres traced from mOC to lTPJ. **(R)** Tracts observed from right sagittal view, when fibres traced from mOC to rTPJ. The number and volume of tracts do not appear significantly different between the two hemispheres. mOC – merged Olfactory Cortex lTPJ – left Temporoparietal Junction rTPJ – right Temporoparietal Junction

**Fig. S23.**
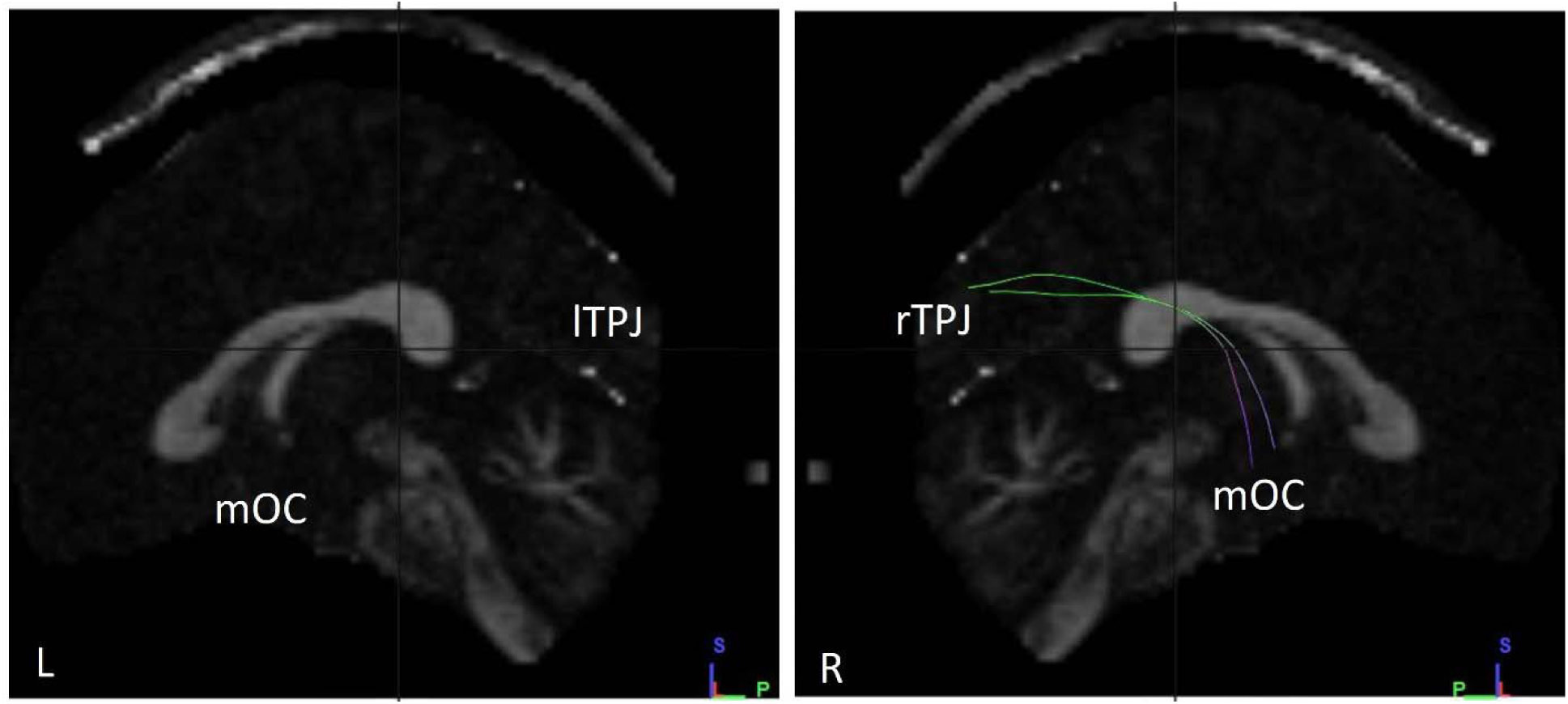
Fibres traced from mOC to lTPJ and rTPJ in subject no. 3. **(L)** No tracts observed from left sagittal view, when fibres traced from mOC to lTPJ. **(R)** Tracts observed from right sagittal view, when fibres traced from mOC to rTPJ. There is a significant difference in the observation between the hemispheres, arising from the complete lack of fibres in the left hemisphere. mOC – merged Olfactory Cortex lTPJ – left Temporoparietal Junction rTPJ – right Temporoparietal Junction

**Fig. S24.**
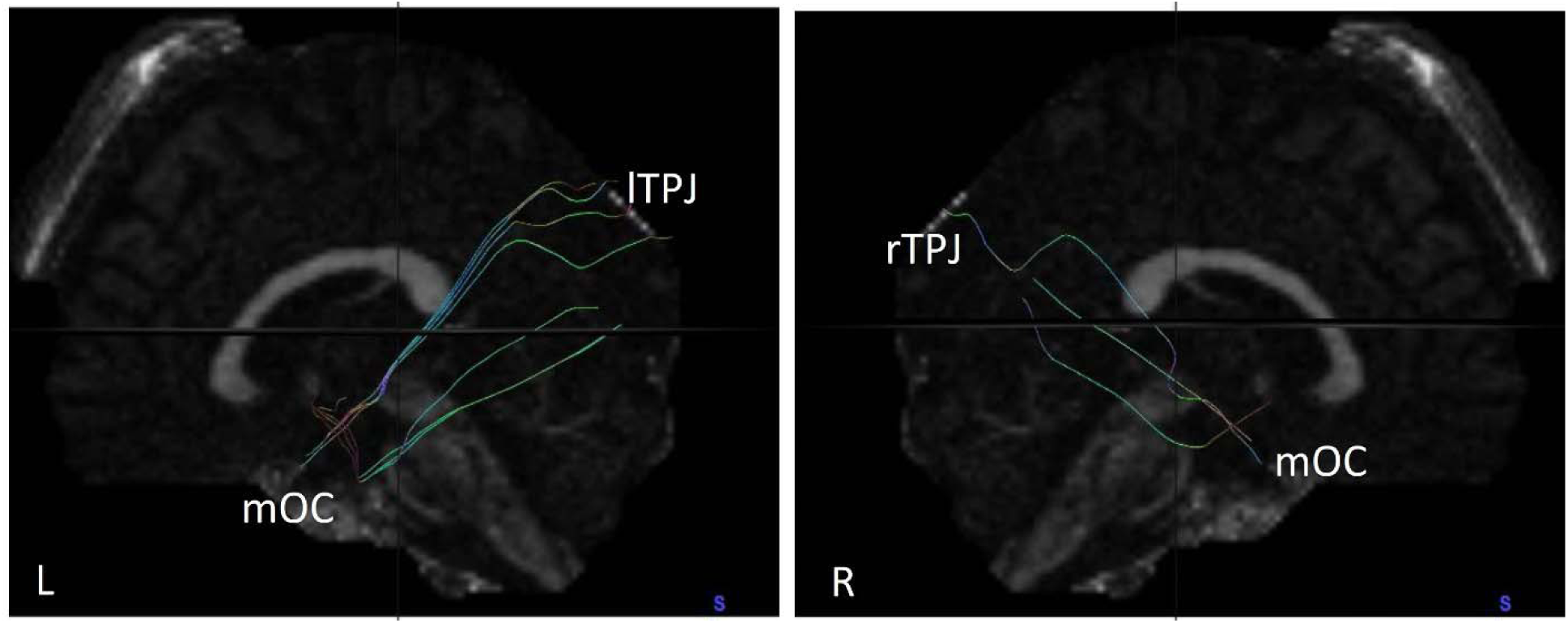
Fibres traced from mOC to lTPJ and rTPJ in subject no. 4. **(L)** Tracts observed from left sagittal view, when fibres traced from mOC to lTPJ. **(R)** Tracts observed from right sagittal view, when fibres traced from mOC to rTPJ. There is a subtle difference in number and volume of fibres observed between the hemispheres with a greater number of fibres noted in the left hemisphere. mOC – merged Olfactory Cortex lTPJ – left Temporoparietal Junction rTPJ – right Temporoparietal Junction

**Fig. S25.**
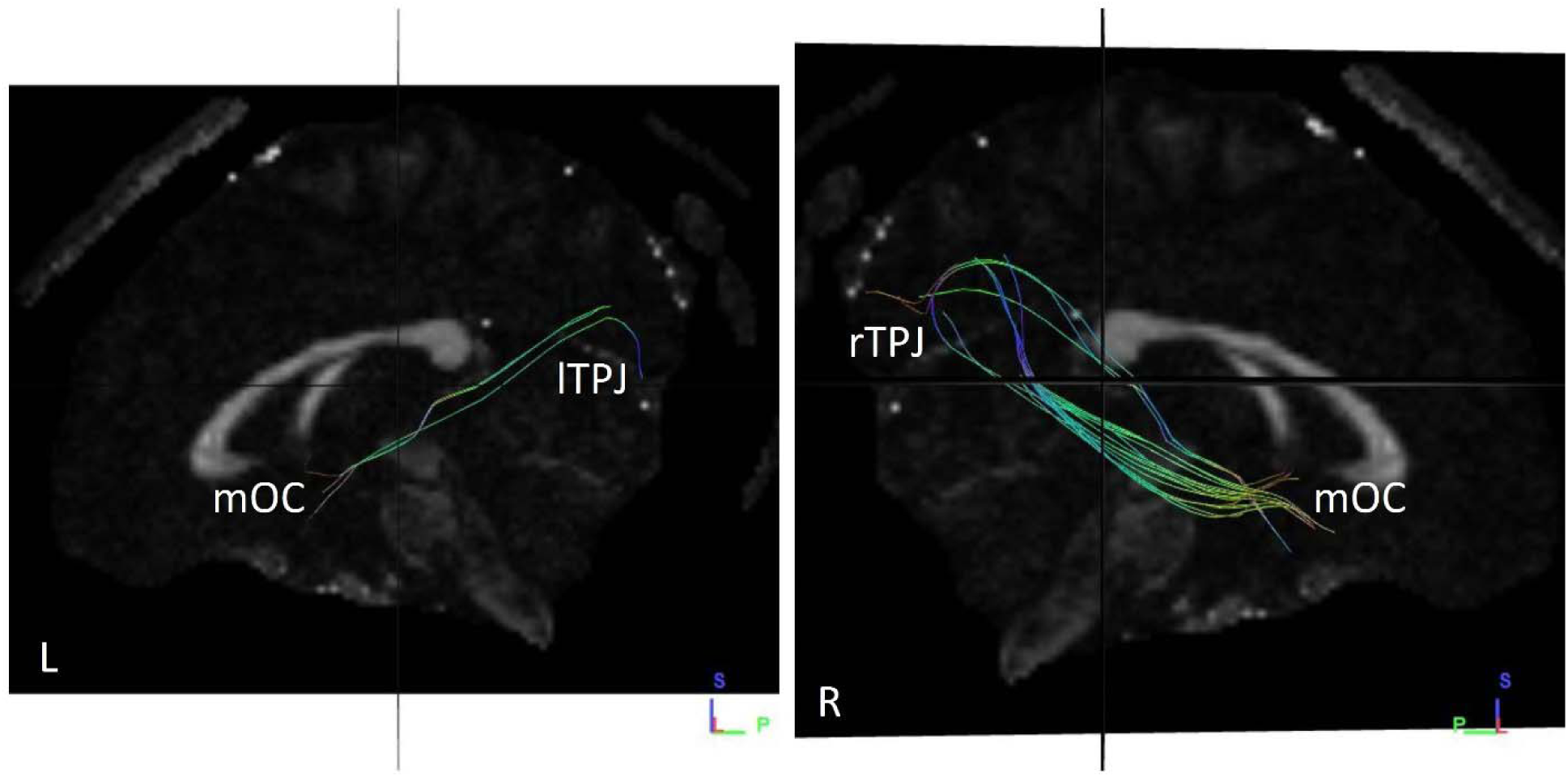
Fibres traced from mOC to lTPJ and rTPJ in subject no. 5. **(L)** Tracts observed from left sagittal view, when fibres traced from mOC to lTPJ. **(R)** Tracts observed from right sagittal view, when fibres traced from mOC to rTPJ. There is a significant difference in number and volume of tracts observed between the hemispheres, with more fibres observed in the right hemisphere. mOC – merged Olfactory Cortex lTPJ – left Temporoparietal Junction rTPJ – right Temporoparietal Junction

**Fig. S26.**
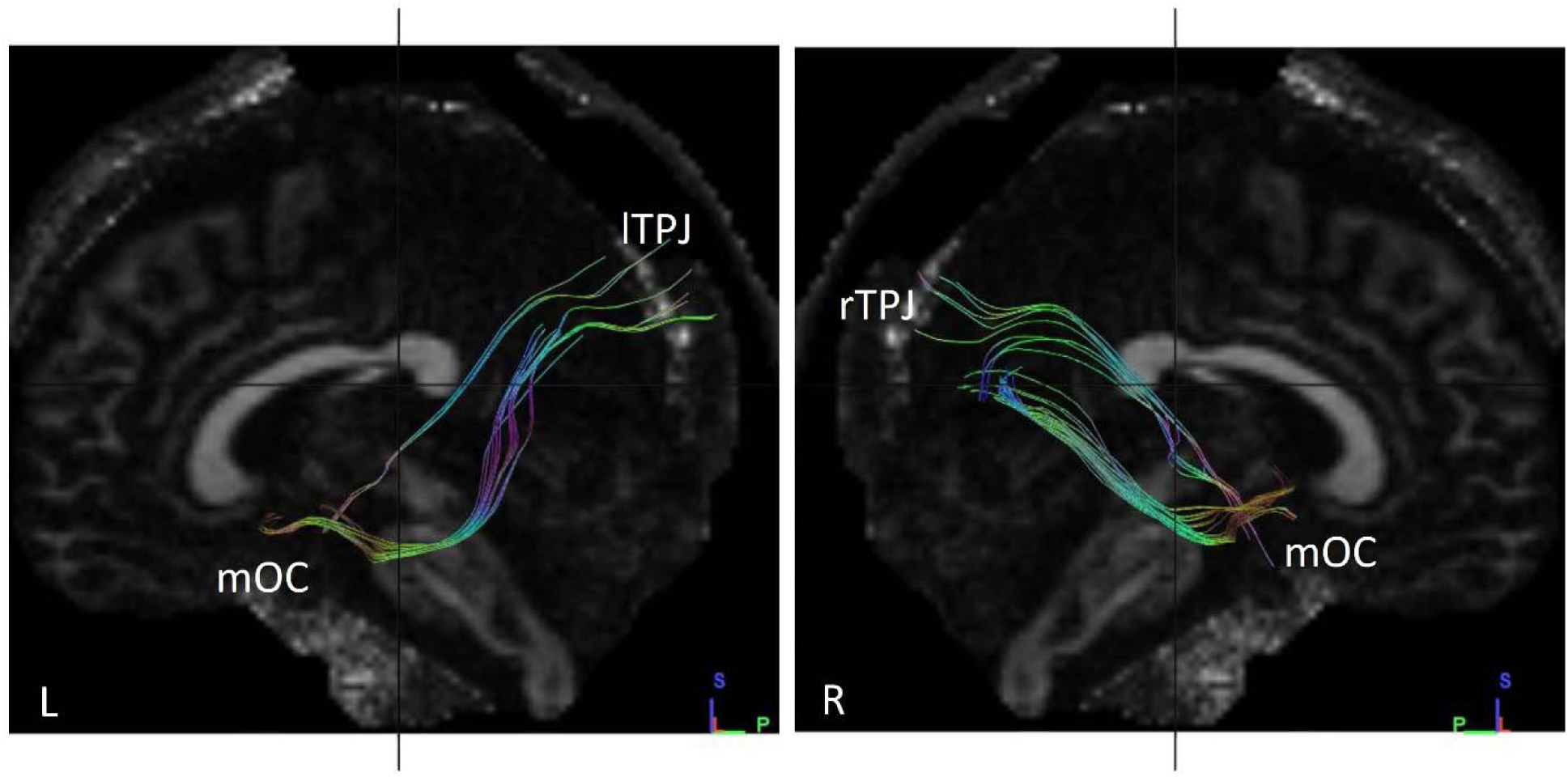
Fibres traced from mOC to lTPJ and rTPJ in subject no. 6. **(L)** Tracts observed from left sagittal view, when fibres traced from mOC to lTPJ. **(R)** Tracts observed from right sagittal view, when fibres traced from mOC to rTPJ. There is a significant difference in number and volume of tracts observed between the hemispheres, with more fibres observed in the right hemisphere. mOC – merged Olfactory Cortex lTPJ – left Temporoparietal Junction rTPJ – right Temporoparietal Junction

**Fig. S27.**
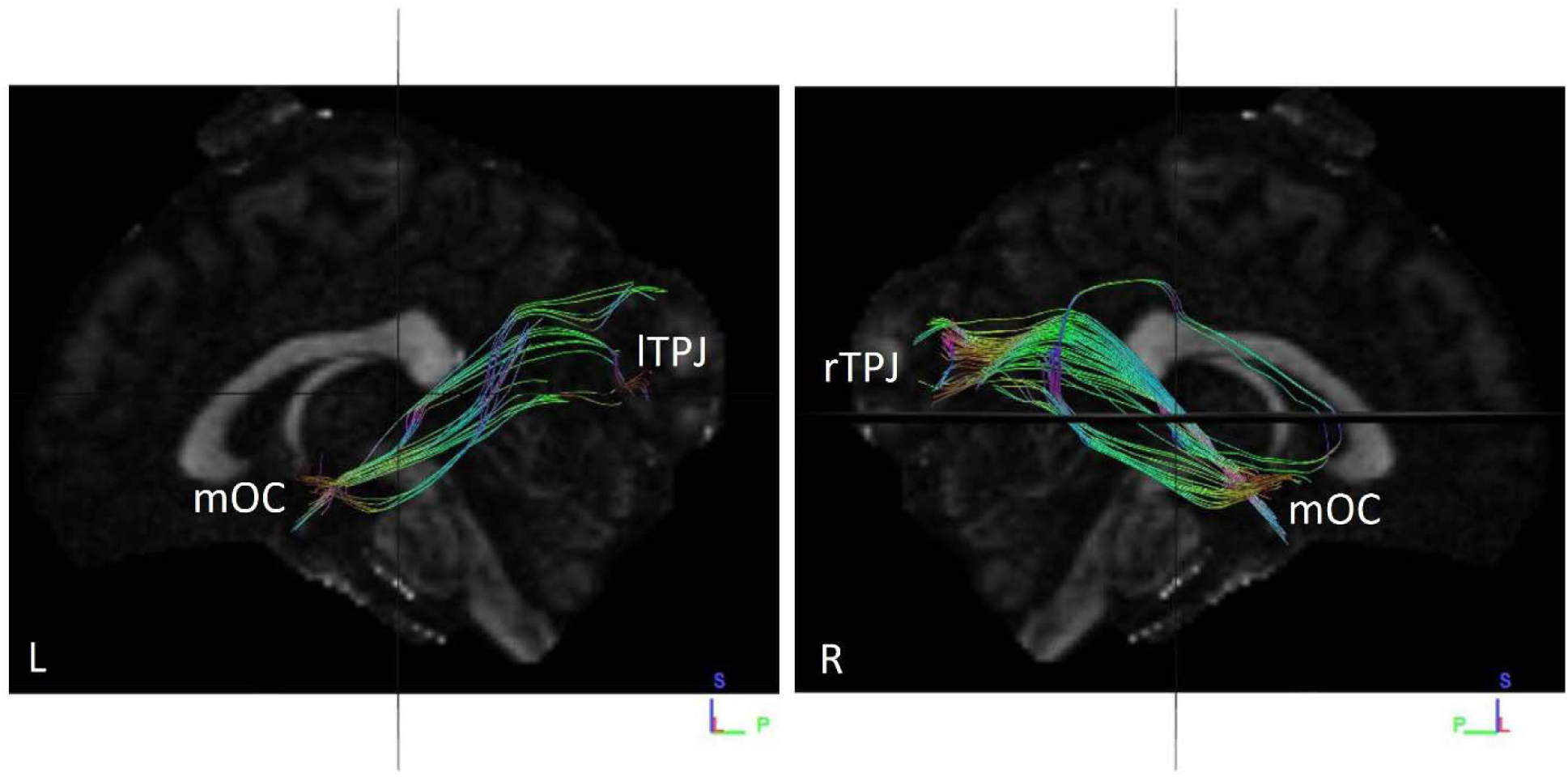
Fibres traced from mOC to lTPJ and rTPJ in subject no. 7. **(L)** Tracts observed from left sagittal view, when fibres traced from mOC to lTPJ. **(R)** Tracts observed from right sagittal view, when fibres traced from mOC to rTPJ. A significant difference in number and volume of tracts is not observed between the hemispheres. mOC – merged Olfactory Cortex lTPJ – left Temporoparietal Junction rTPJ – right Temporoparietal Junction

**Fig. S28.**
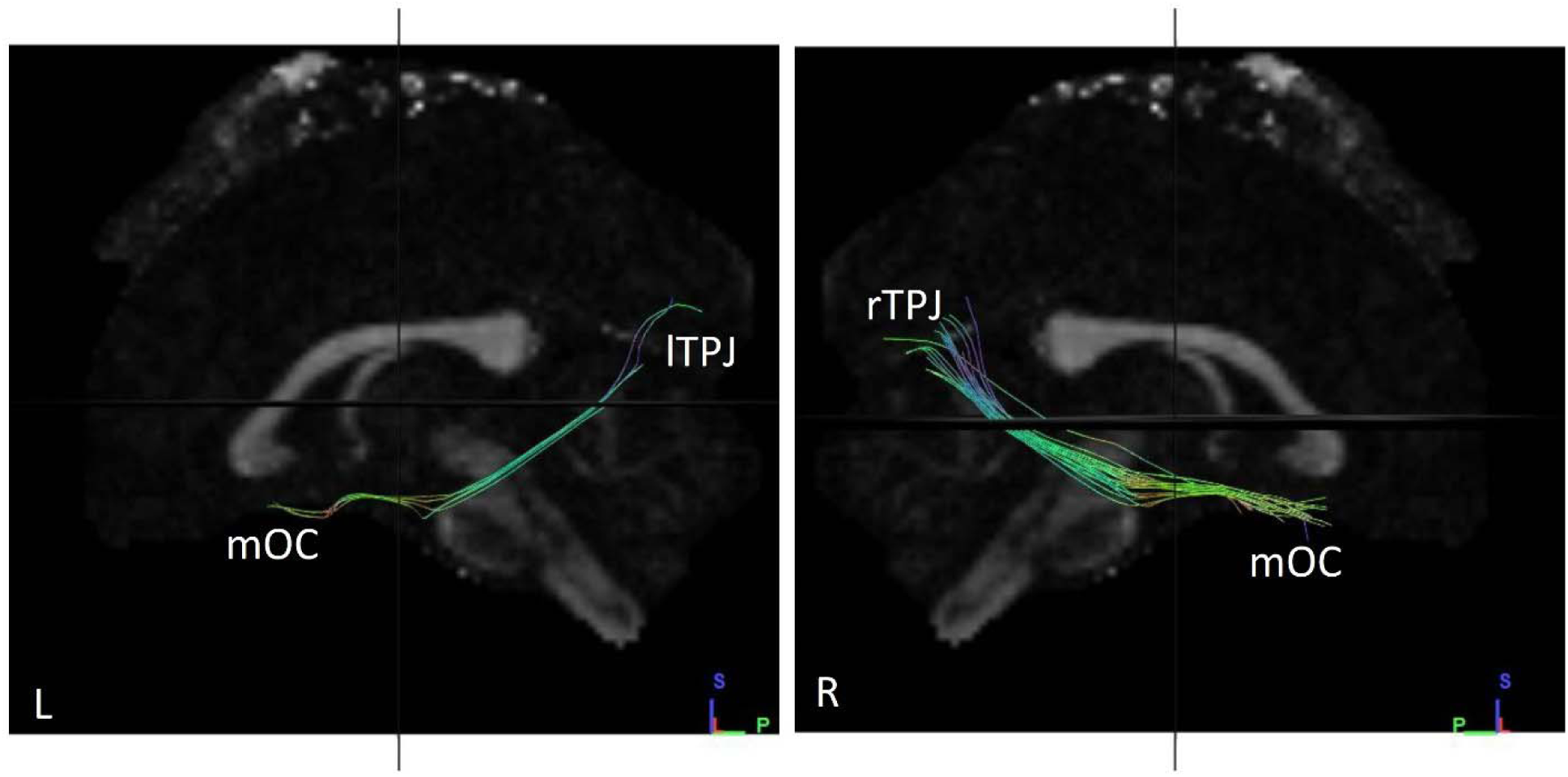
Fibres traced from mOC to lTPJ and rTPJ in subject no. 8. **(L)** Tracts observed from left sagittal view, when fibres traced from mOC to lTPJ. **(R)** Tracts observed from right sagittal view, when fibres traced from mOC to rTPJ. A significant difference in number and volume of tracts is observed between the hemispheres, with more fibres in the right hemisphere. mOC – merged Olfactory Cortex lTPJ – left Temporoparietal Junction rTPJ – right Temporoparietal Junction

**Fig. S39.**
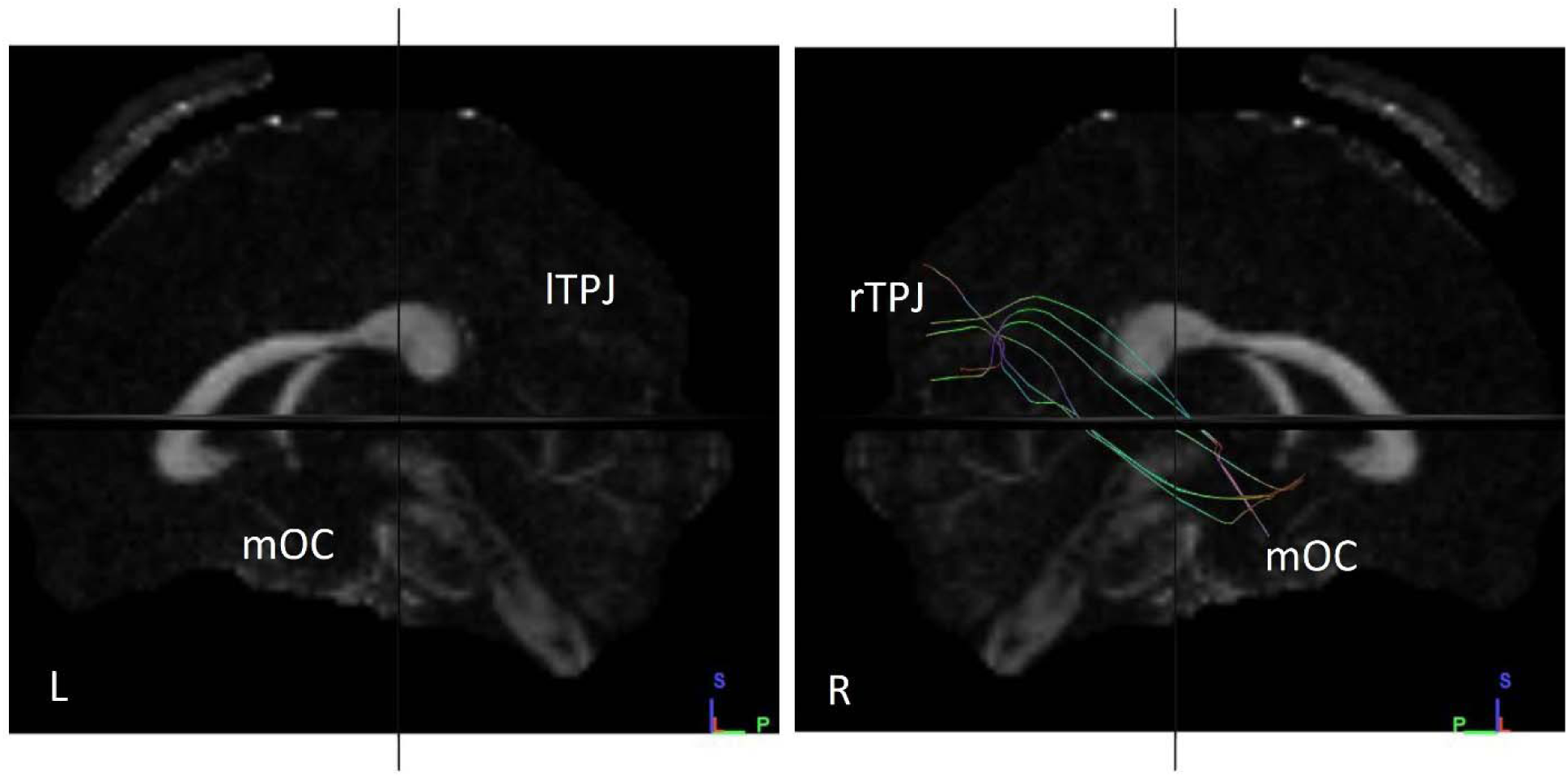
Fibres traced from mOC to lTPJ and rTPJ in subject no. 9. **(L)** Tracts observed from left sagittal view, when fibres traced from mOC to lTPJ. **(R)** Tracts observed from right sagittal view, when fibres traced from mOC to rTPJ. A significant difference in number and volume of tracts is not observed between the hemispheres. mOC – merged Olfactory Cortex lTPJ – left Temporoparietal Junction rTPJ – right Temporoparietal Junction

**Fig. S30.**
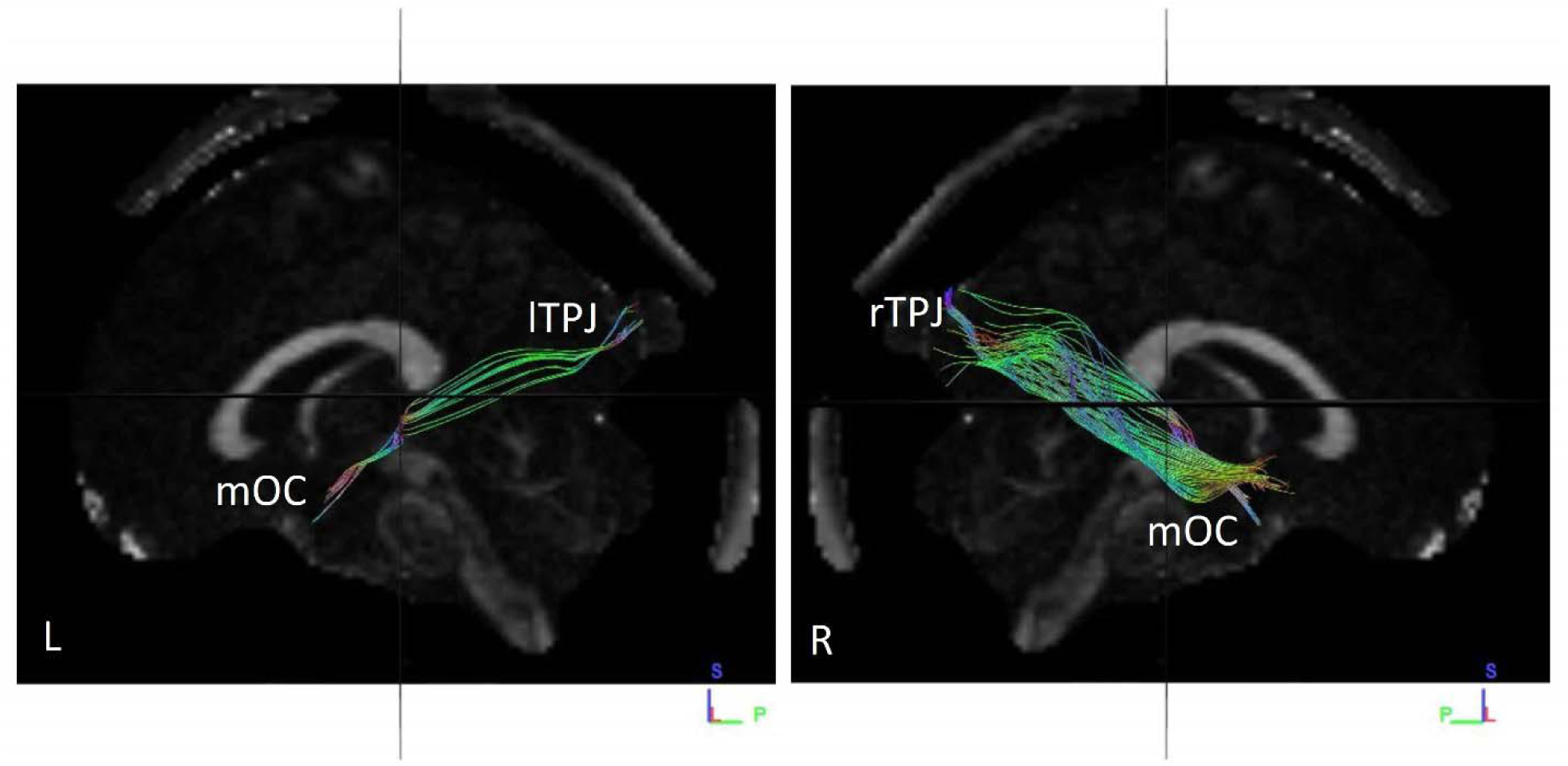
Fibres traced from mOC to lTPJ and rTPJ in subject no. 10. **(L)** Tracts observed from left sagittal view, when fibres traced from mOC to lTPJ. **(R)** Tracts observed from right sagittal view, when fibres traced from mOC to rTPJ. A significant difference in number and volume of tracts is observed between the hemispheres, with a greater number of fibres in the right hemisphere. mOC – merged Olfactory Cortex lTPJ – left Temporoparietal Junction rTPJ – right Temporoparietal Junction

**Fig. S31.**
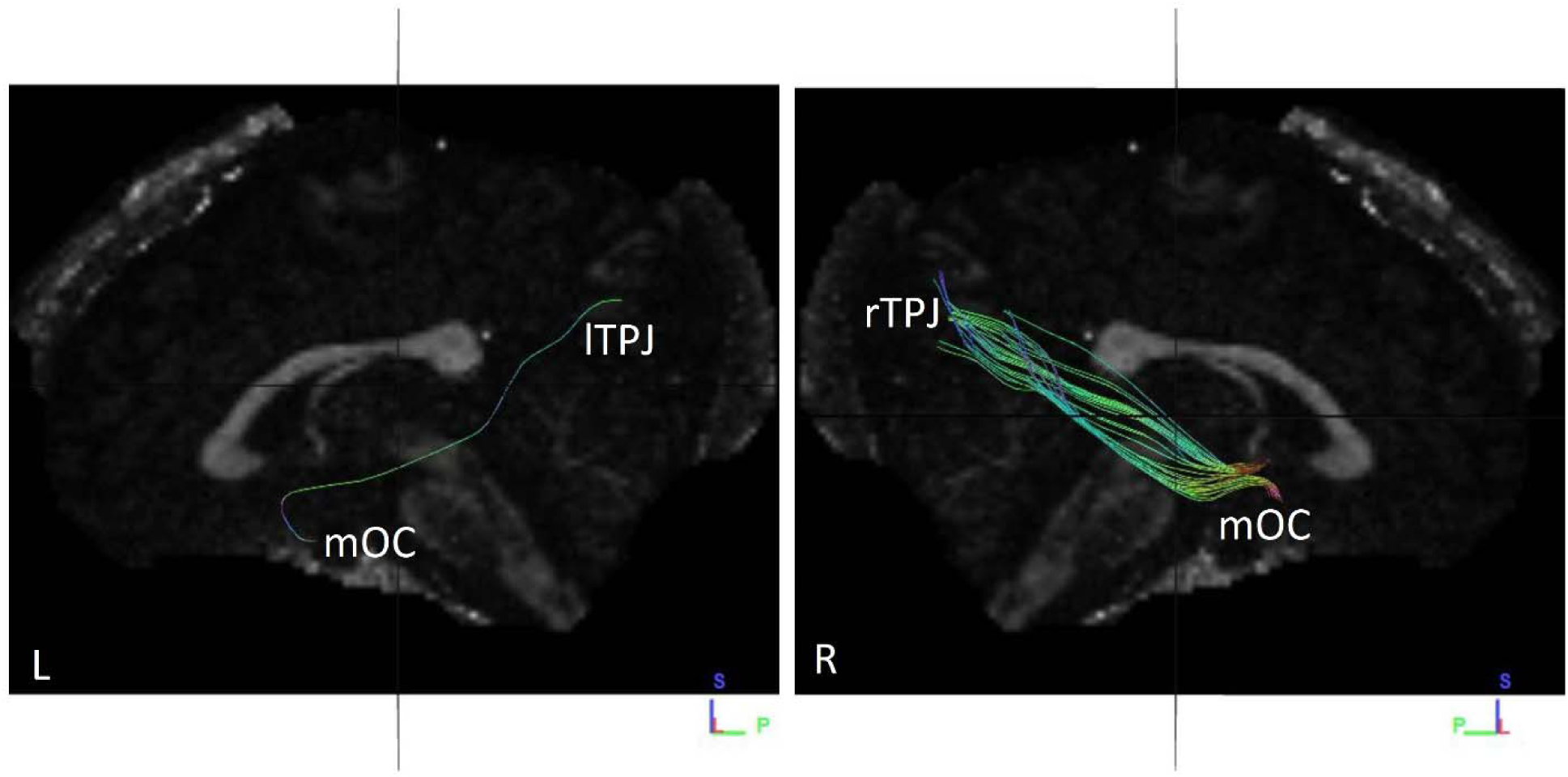
Fibres traced from mOC to lTPJ and rTPJ in subject no. 11. **(L)** Tracts observed from left sagittal view, when fibres traced from mOC to lTPJ. **(R)** Tracts observed from right sagittal view, when fibres traced from mOC to rTPJ. A significant difference in number and volume of tracts is observed between the hemispheres, with a greater number of fibres in the right hemisphere. mOC – merged Olfactory Cortex lTPJ – left Temporoparietal Junction rTPJ – right Temporoparietal Junction

**Fig. S32.**
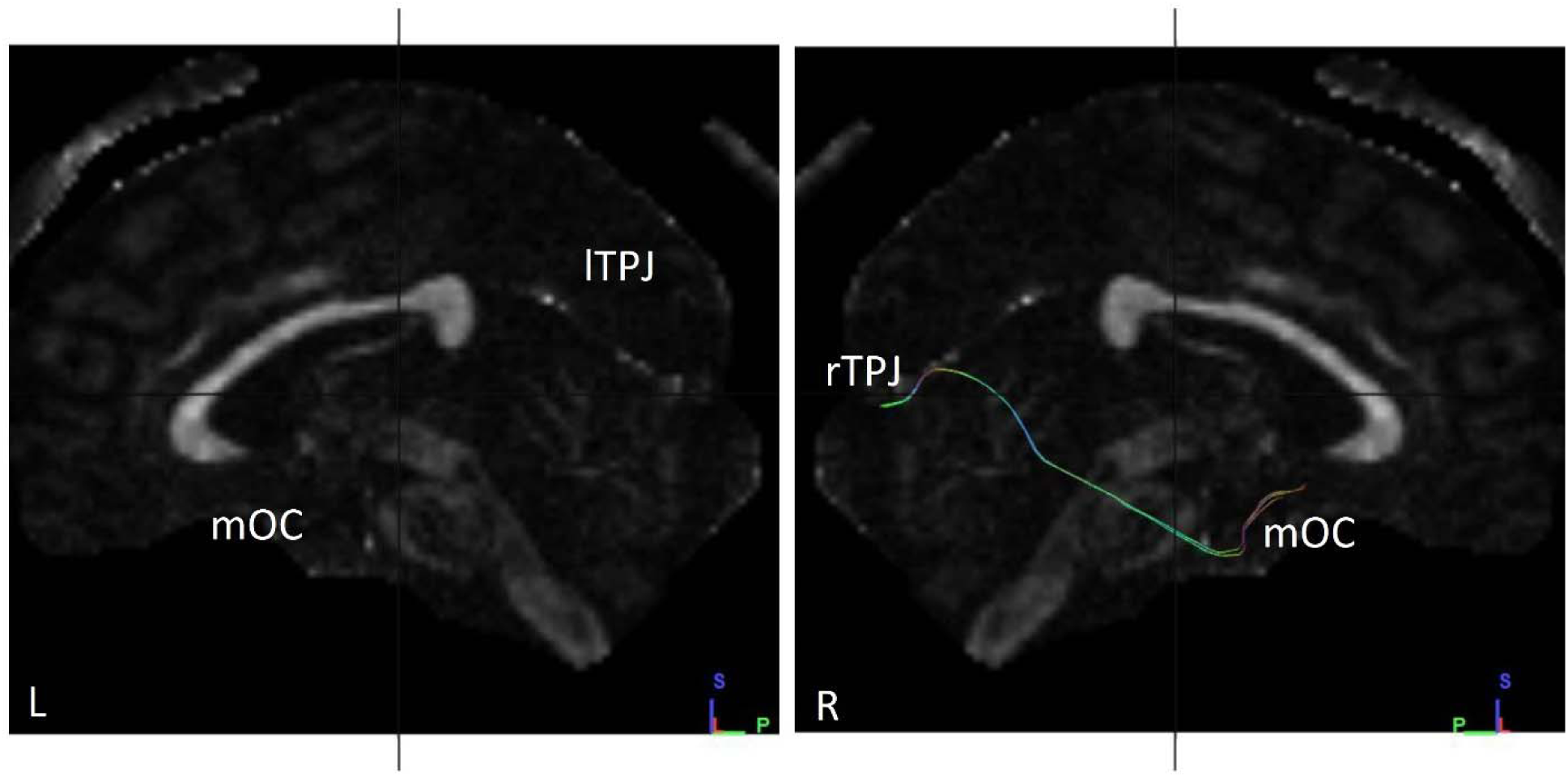
Fibres traced from mOC to lTPJ and rTPJ in subject no. 12. **(L)** No tracts observed from left sagittal view, when fibres traced from mOC to lTPJ. **(R)** Tracts observed from right sagittal view, when fibres traced from mOC to rTPJ. A significant difference in number and volume of tracts is observed between the hemispheres arising from the total lack of fibres in the left hemisphere. mOC – merged Olfactory Cortex lTPJ – left Temporoparietal Junction rTPJ – right Temporoparietal Junction

**Fig. S33.**
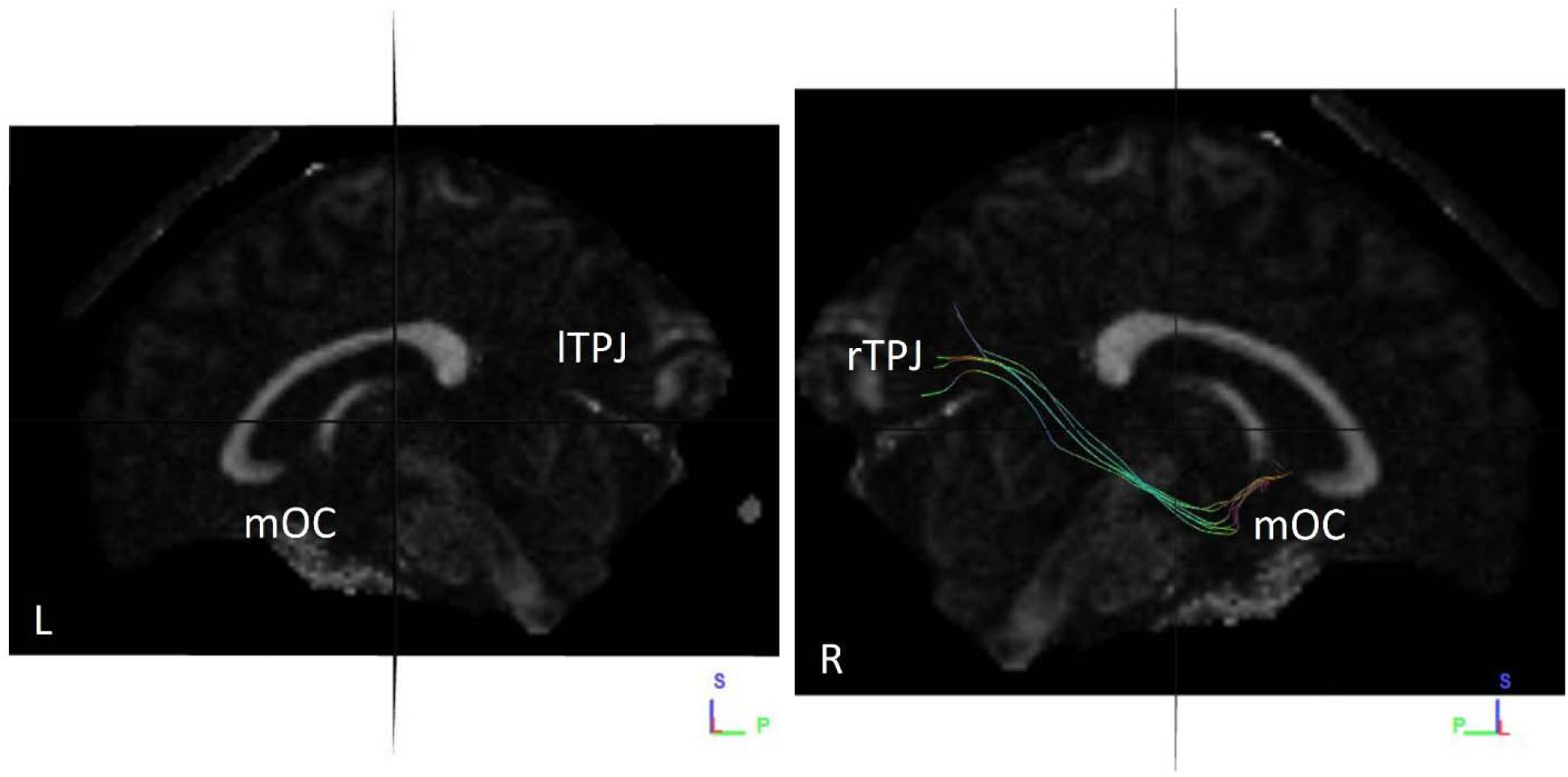
Fibres traced from mOC to lTPJ and rTPJ in subject no. 13. **(L)** No tracts observed from left sagittal view, when fibres traced from mOC to lTPJ. **(R)** Tracts observed from right sagittal view, when fibres traced from mOC to rTPJ. A significant difference in number and volume of tracts is observed between the hemispheres arising from the total lack of fibres in the left hemisphere. mOC – merged Olfactory Cortex lTPJ – left Temporoparietal Junction rTPJ – right Temporoparietal Junction

**Fig. S34.**
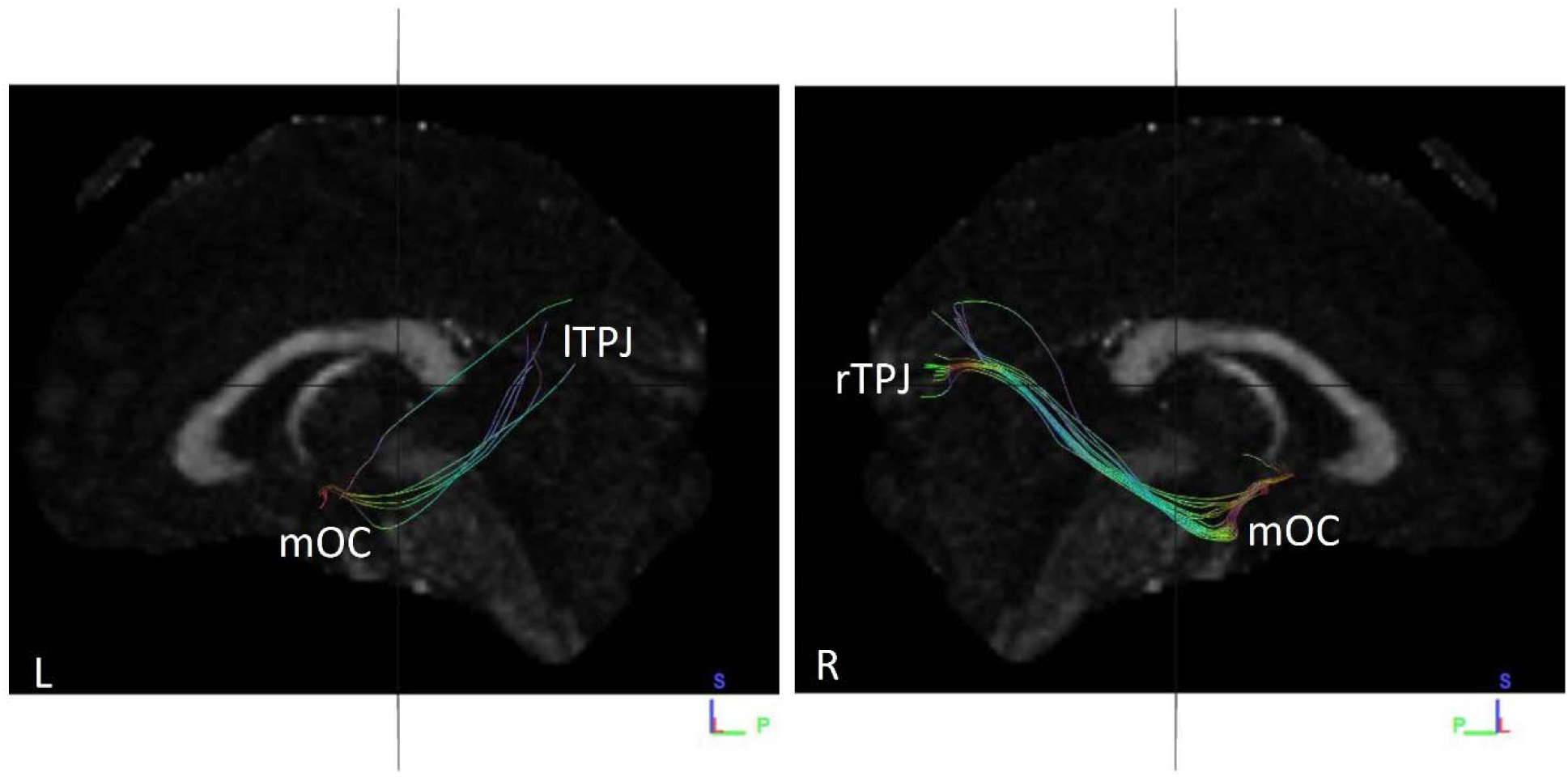
Fibres traced from mOC to lTPJ and rTPJ in subject no. 14. **(L)** Tracts observed from left sagittal view, when fibres traced from mOC to lTPJ. **(R)** Tracts observed from right sagittal view, when fibres traced from mOC to rTPJ. A significant difference in number and volume of tracts is observed between the hemispheres, with right hemisphere having a greater number of fibres. mOC – merged Olfactory Cortex lTPJ – left Temporoparietal Junction rTPJ – right Temporoparietal Junction

**Fig. S35.**
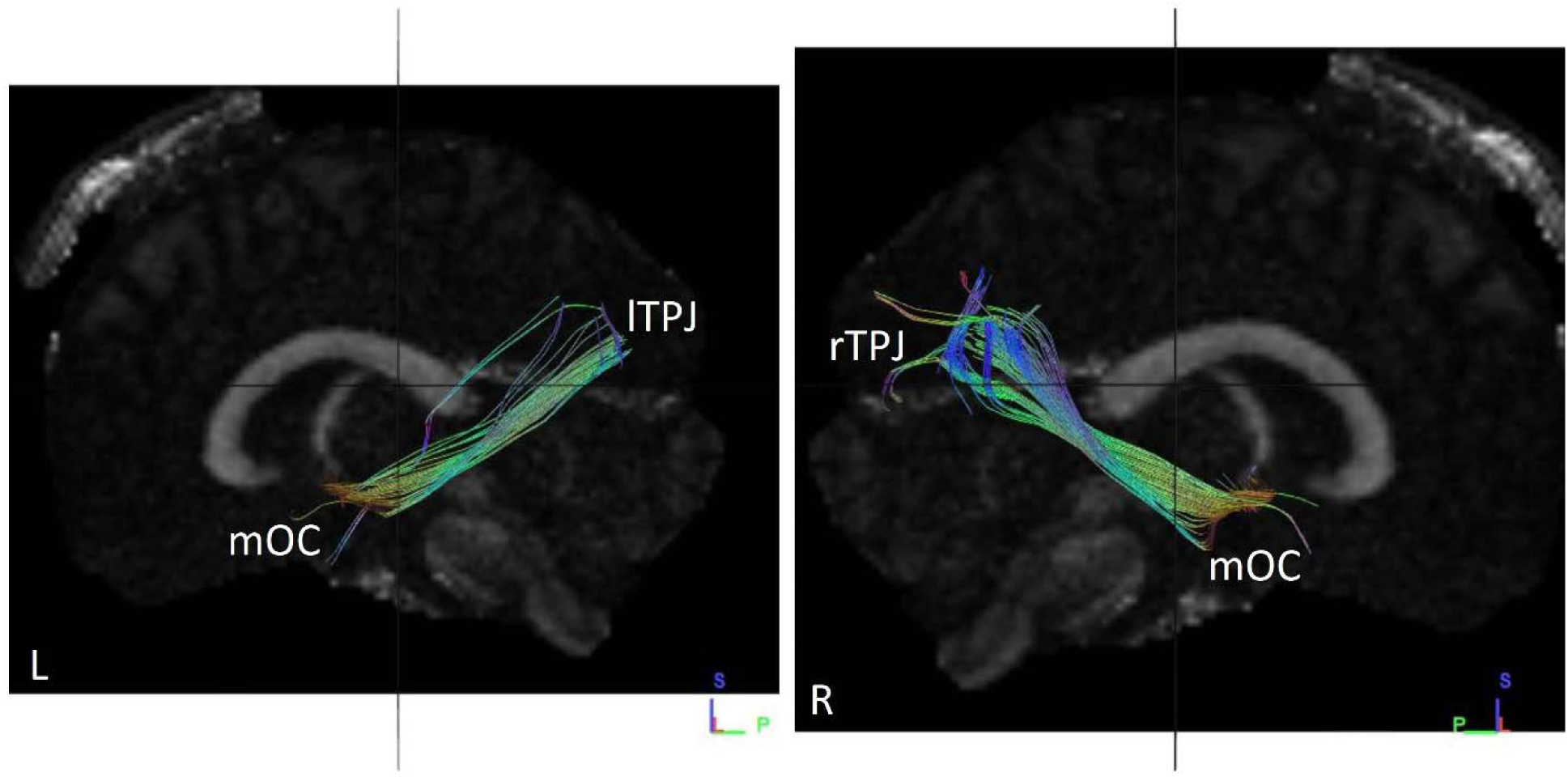
Fibres traced from mOC to lTPJ and rTPJ in subject no. 15. **(L)** Tracts observed from left sagittal view, when fibres traced from mOC to lTPJ. **(R)** Tracts observed from right sagittal view, when fibres traced from mOC to rTPJ. A significant difference in number and volume of tracts is not observed between the hemispheres. mOC – merged Olfactory Cortex lTPJ – left Temporoparietal Junction rTPJ – right Temporoparietal Junction

**Fig. S36.**
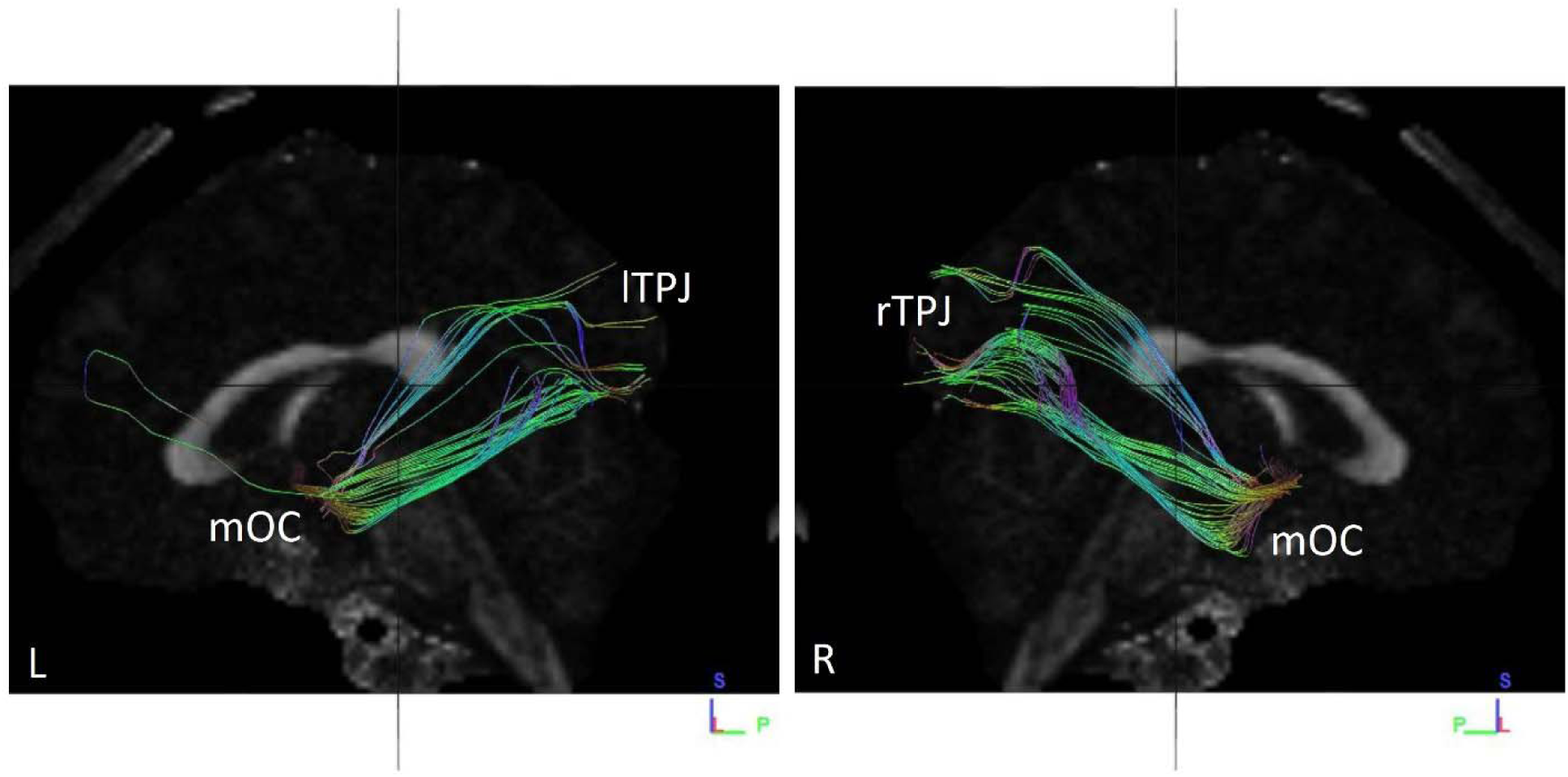
Fibres traced from mOC to lTPJ and rTPJ in subject no. 16. **(L)** Tracts observed from left sagittal view, when fibres traced from mOC to lTPJ. **(R)** Tracts observed from right sagittal view, when fibres traced from mOC to rTPJ. A significant difference in number and volume of tracts is not observed between the hemispheres mOC – merged Olfactory Cortex lTPJ – left Temporoparietal Junction rTPJ – right Temporoparietal Junction

**Fig. S37.**
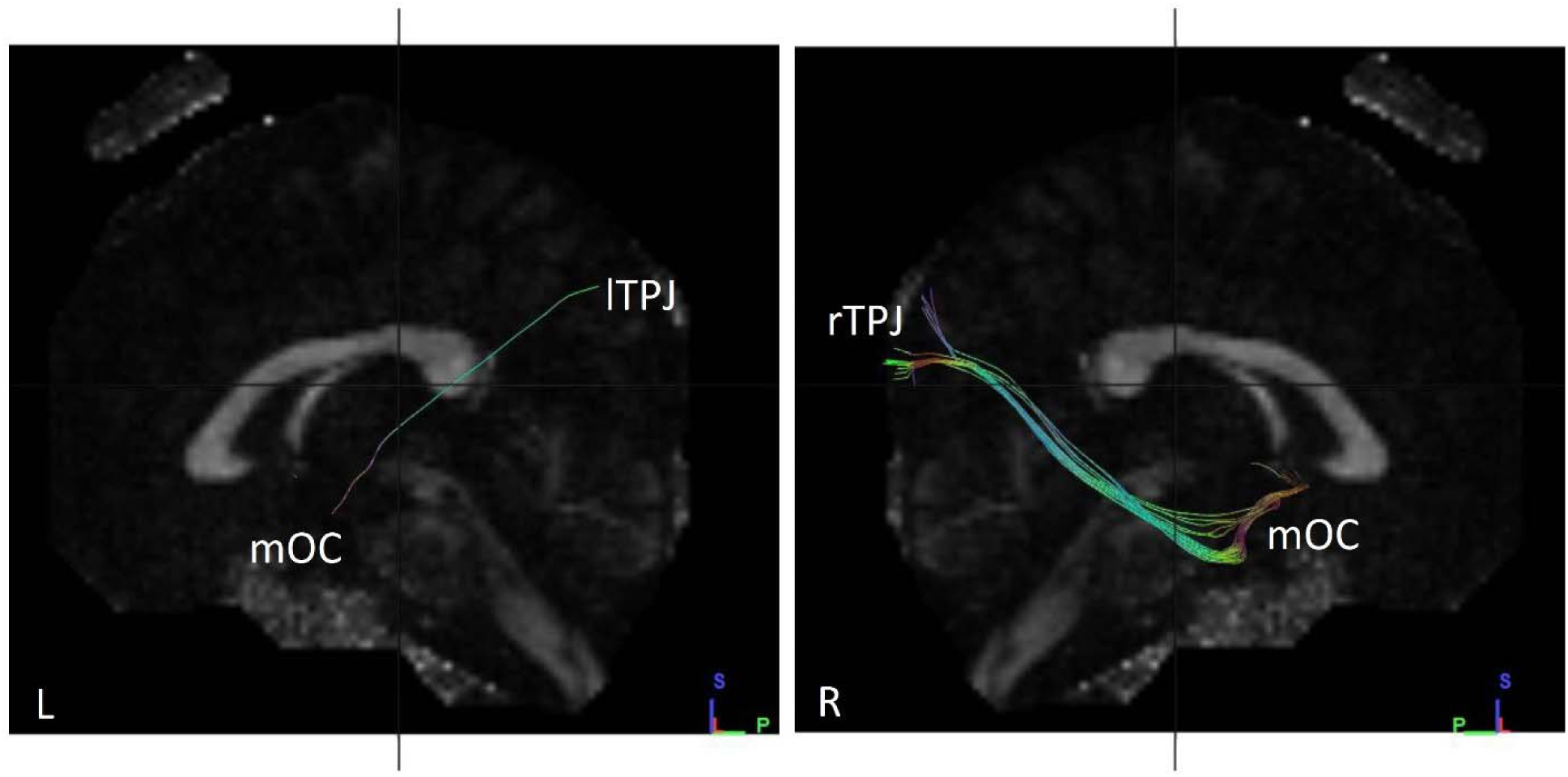
Fibres traced from mOC to lTPJ and rTPJ in subject no. 17. **(L)** Tracts observed from left sagittal view, when fibres traced from mOC to lTPJ. **(R)** Tracts observed from right sagittal view, when fibres traced from mOC to rTPJ. A significant difference in number and volume of tracts is observed between the hemispheres, with a much greater number of fibres observed in the right hemisphere. mOC – merged Olfactory Cortex lTPJ – left Temporoparietal Junction rTPJ – right Temporoparietal Junction

**Fig. S38.**
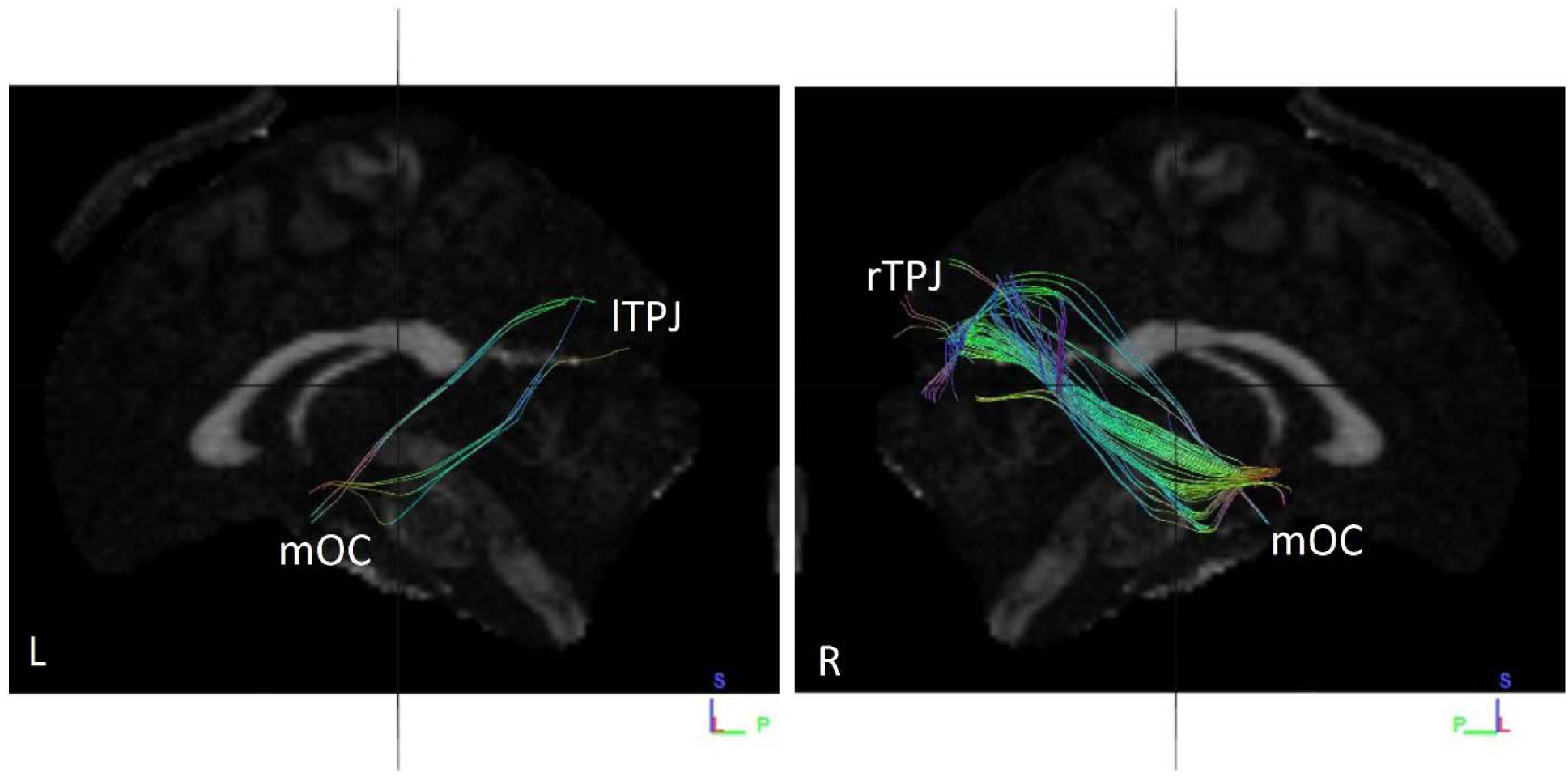
Fibres traced from mOC to lTPJ and rTPJ in subject no. 18. **(L)** Tracts observed from left sagittal view, when fibres traced from mOC to lTPJ. **(R)** Tracts observed from right sagittal view, when fibres traced from mOC to rTPJ. A significant difference in number and volume of tracts is observed between the hemispheres, with a much greater number of fibres observed in the right hemisphere. mOC – merged Olfactory Cortex lTPJ – left Temporoparietal Junction rTPJ – right Temporoparietal Junction

**Fig. S39.**
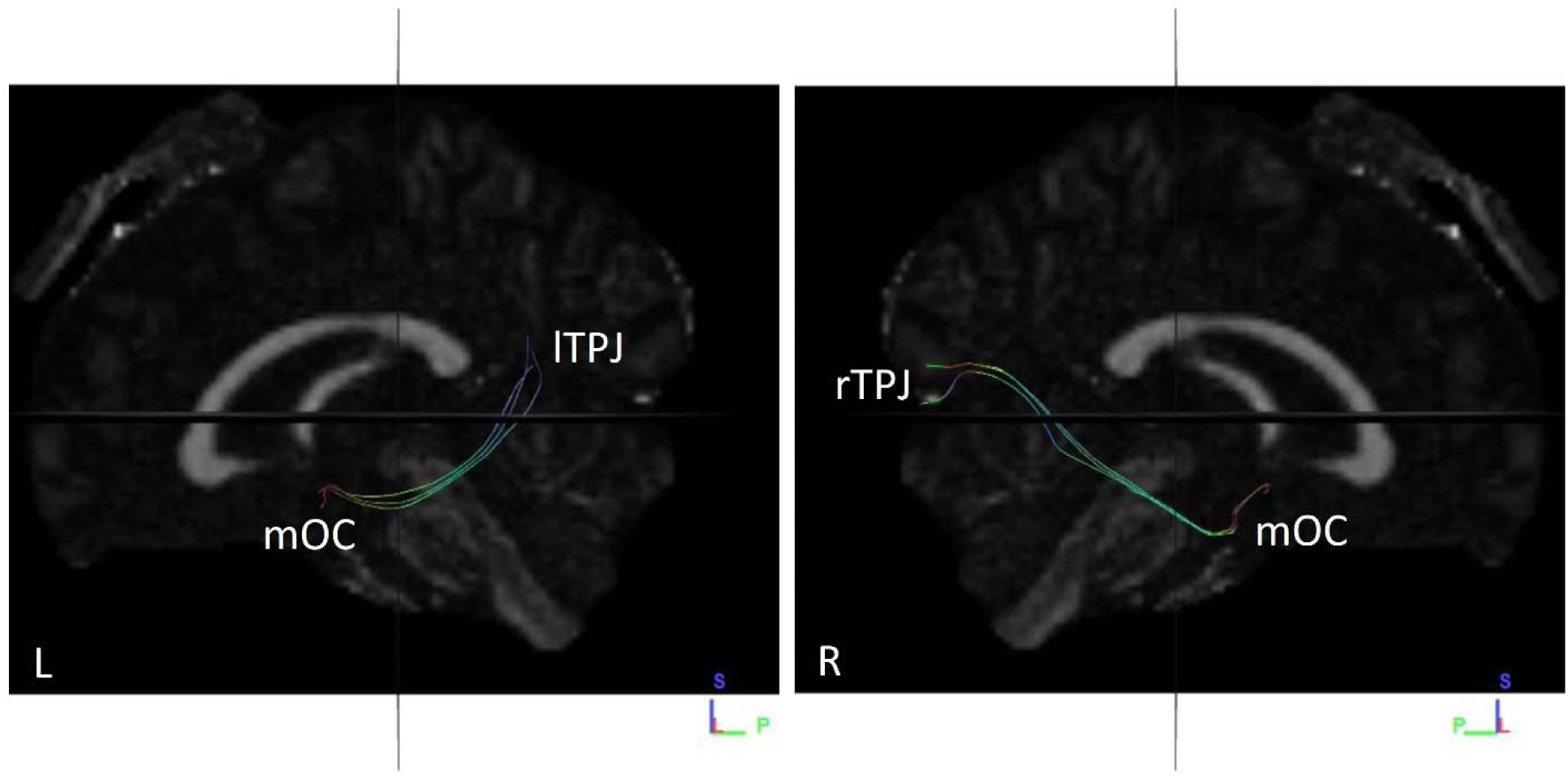
Fibres traced from mOC to lTPJ and rTPJ in subject no. 19. **(L)** Tracts observed from left sagittal view, when fibres traced from mOC to lTPJ. **(R)** Tracts observed from right sagittal view, when fibres traced from mOC to rTPJ. A significant difference in number and volume of tracts is not observed between the hemispheres. mOC – merged Olfactory Cortex lTPJ – left Temporoparietal Junction rTPJ – right Temporoparietal Junction

**Fig. S40.**
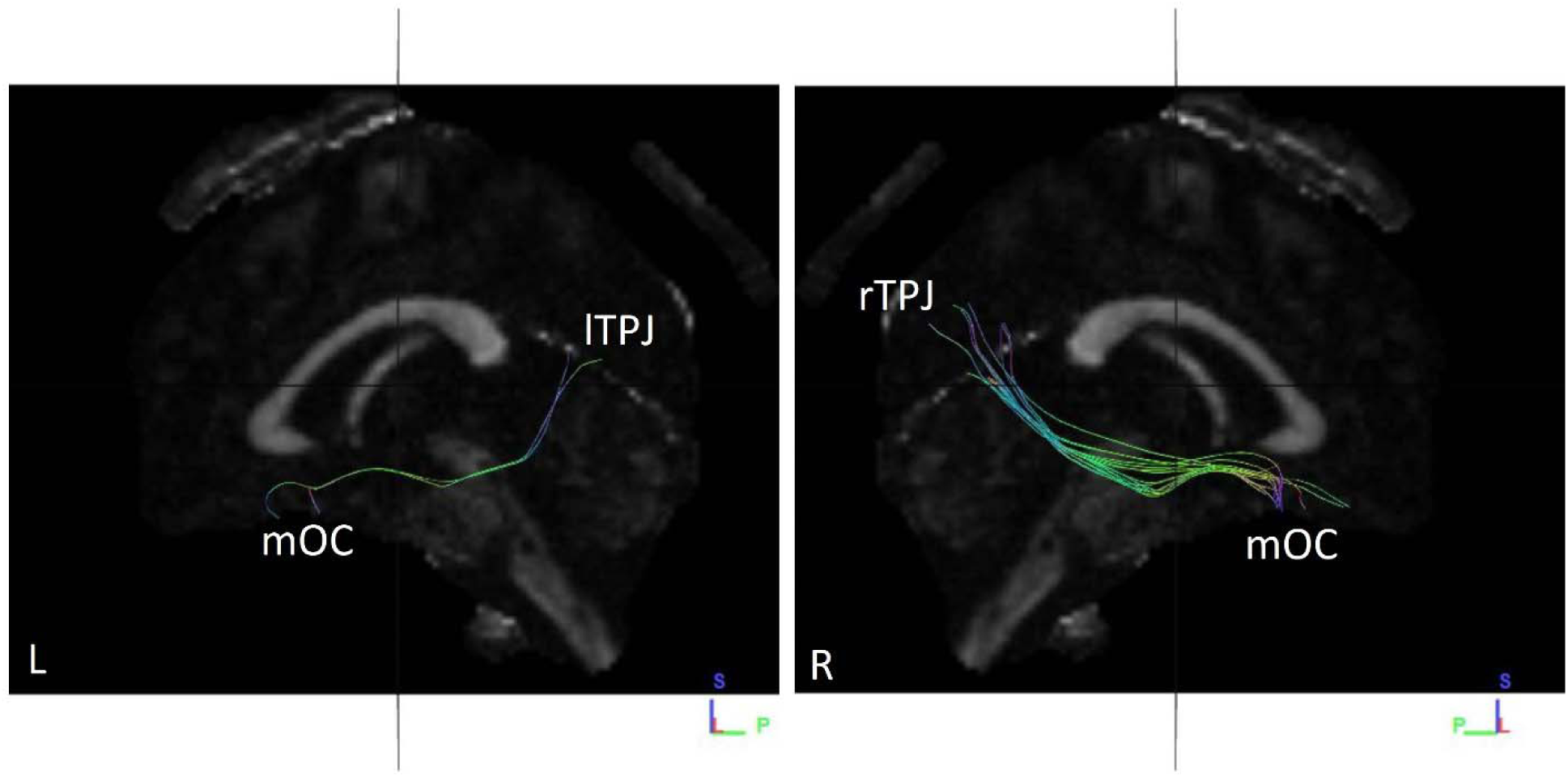
Fibres traced from mOC to lTPJ and rTPJ in subject no. 20. **(L)** Tracts observed from left sagittal view, when fibres traced from mOC to lTPJ. **(R)** Tracts observed from right sagittal view, when fibres traced from mOC to rTPJ. A significant difference in number and volume of tracts is observed between the hemispheres, with a greater number of fibres in the right hemisphere. mOC – merged Olfactory Cortex lTPJ – left Temporoparietal Junction rTPJ – right Temporoparietal Junction

**Table S1.** Sex and age category of subjects.

| **SUBJECT NUMBER** | **SEX** | **AGE GROUP (in years)** |
| --- | --- | --- |
| **1** | **F** | **20-24** |
| **2** | **F** | **20-24** |
| **3** | **F** | **20-24** |
| **4** | **F** | **25-29** |
| **5** | **F** | **25-29** |
| **6** | **F** | **25-29** |
| **7** | **F** | **25-29** |
| **8** | **F** | **25-29** |
| **9** | **F** | **35-39** |
| **10** | **F** | **40-44** |
| **11** | **M** | **20-24** |
| **12** | **M** | **25-29** |
| **13** | **M** | **25-29** |
| **14** | **M** | **25-29** |
| **15** | **M** | **30-34** |
| **16** | **M** | **30-34** |
| **17** | **M** | **30-34** |
| **18** | **M** | **35-39** |
| **19** | **M** | **35-39** |
| **20** | **M** | **35-39** |

**Table S2.** Comparison of number of tracts found in male and female subjects.

| **Female** |  | **Male** |  |
| --- | --- | --- | --- |
| **Subject Number** | **Number of tracts found** | **Subject Number** | **Number of tracts found** |
| **1** | 44 | **11** | 40 |
| **2** | 16 | **12** | 3 |
| **3** | 2 | **13** | 6 |
| **4** | 11 | **14** | 34 |
| **5** | 27 | **15** | 256 |
| **6** | 58 | **16** | 122 |
| **7** | 105 | **17** | 37 |
| **8** | 30 | **18** | 100 |
| **9** | 16 | **19** | 6 |
| **10** | 96 | **20** | 17 |

**Table S3.** Comparison of length of tracts in male and female subjects.

| **Female** |  | **Male** |  |
| --- | --- | --- | --- |
| **Subject Number** | Mean length of tracts found (mm) | **Subject Number** | Mean length of tracts found (mm) |
| **1** | 100.65 | **11** | 96.78 |
| **2** | 98.21 | **12** | 97.96 |
| **3** | 80.73 | **13** | 92.87 |
| **4** | 109.33 | **14** | 110.16 |
| **5** | 97.53 | **15** | 102.35 |
| **6** | 101.61 | **16** | 107.66 |
| **7** | 108.97 | **17** | 96.20 |
| **8** | 87.25 | **18** | 97.4 |
| **9** | 105.23 | **19** | 100.14 |
| **10** | 92.38 | **20** | 87.47 |

**Table S4.** Comparison of volume of tracts in male and female subjects.

| **Female** |  | **Male** |  |
| --- | --- | --- | --- |
| **Subject Number** | Volume of tracts<br>(mm<sup>3</sup>) | **Subject Number** | Volume of tracts<br>(mm<sup>3</sup>) |
| **1** | 5059.13 | **11** | 2997 |
| **2** | 2106 | **12** | 678.37 |
| **3** | 199.12 | **13** | 1113.75 |
| **4** | 2224.13 | **14** | 2949.75 |
| **5** | 3942 | **15** | 7779.38 |
| **6** | 5460.75 | **16** | 11005.90 |
| **7** | 9959.63 | **17** | 3975.75 |
| **8** | 2824.87 | **18** | 6166.12 |
| **9** | 2814.75 | **19** | 1198.12 |
| **10** | 6240.37 | **20** | 1940.62 |

**Table S5.** Distribution of number of tracts in right and left hemispheres.

| SUBJECT NUMBER | LEFT HEMISPHERE | RIGHT HEMISPHERE |
| --- | --- | --- |
| 1 | 3 | 41 |
| 2 | 13 | 3 |
| 3 | 0 | 2 |
| 4 | 8 | 3 |
| 5 | 9 | 18 |
| 6 | 20 | 38 |
| 7 | 27 | 78 |
| 8 | 5 | 25 |
| 9 | 7 | 9 |
| 10 | 15 | 81 |
| 11 | 1 | 39 |
| 12 | 0 | 3 |
| 13 | 0 | 6 |
| 14 | 6 | 28 |
| 15 | 53 | 203 |
| 16 | 22 | 100 |
| 17 | 1 | 36 |
| 18 | 7 | 93 |
| 19 | 3 | 3 |
| 20 | 2 | 15 |

**Table S6.** Comparison of length of tracts in right and left hemispheres.

| SUBJECT NUMBER | MEAN LENGTH<br>(mm) |  |
| --- | --- | --- |
|  | Left Hemisphere | Right Hemisphere |
| 1 | **105.61** | **99.9** |
| 2 | **98.77** | **102.04** |
| 3 | **0** | **81.95** |
| 4 | **120.5** | **96.82** |
| 5 | **116.26** | **89.01** |
| 6 | **113.3** | **93.83** |
| 7 | **105.19** | **104.55** |
| 8 | **94.11** | **85.87** |
| 9 | **103.72** | **106.79** |
| 10 | **101.53** | **91.87** |
| 11 | **108.85** | **92.49** |
| 12 | **0** | **97.96** |
| 13 | **0** | **89.92** |
| 14 | **87.72** | **115.99** |
| 15 | **101.92** | **101.46** |
| 16 | **104.34** | **106.45** |
| 17 | **133.51** | **95.47** |
| 18 | **85.57** | **98.18** |
| 19 | **109.93** | **90.34** |
| 20 | **95.29** | **87.59** |

**Table S7.** Comparison of volume of tracts in right and left hemispheres.

| SUBJECT NUMBER | VOLUME<br>(mm<sup>3</sup>) |  |
| --- | --- | --- |
|  | Left Hemisphere | Right Hemisphere |
| 1 | 793.12 | 4299.75 |
| 2 | 1744.88 | 759.37 |
| 3 | 0 | 384.75 |
| 4 | 1728 | 843.75 |
| 5 | 1768.5 | 2635.87 |
| 6 | 2092.5 | 3179.25 |
| 7 | 3611.25 | 6345 |
| 8 | 654.75 | 2170.12 |
| 9 | 1417.5 | 1920.37 |
| 10 | 1863 | 5146.87 |
| 11 | 249.75 | 3118.5 |
| 12 | 0 | 678.37 |
| 13 | 0 | 1255.5 |
| 14 | 1039.5 | 2099.25 |
| 15 | 3283.88 | 6480 |
| 16 | 3145.5 | 6773.62 |
| 17 | 297 | 4060.13 |
| 18 | 1174.5 | 5899.5 |
| 19 | 546.75 | 651.37 |
| 20 | 371.25 | 1768.5 |

**Table S8.** Comparison of tract number and volume in male and female subject on the basis of right and left hemisphere (H).

| TRACT NUMBER |  |  |  |  |  | TRACT VOLUME<br>(mm<sup>3</sup>) |  |  |  |  |  |
| --- | --- | --- | --- | --- | --- | --- | --- | --- | --- | --- | --- |
| Female |  |  | Male |  |  | Female |  |  | Male |  |  |
| Subject no. | Right H. | Left H. | Subject no. | Right H. | Left H. | Subject no. | Right H. | Left H. | Subject no. | Right H. | Left H. |
| 1 | **41** | **3** | **11** | **39** | **1** | 1 | **4299.75** | **793.12** | 11 | **3118.5** | **249.75** |
| 2 | **3** | **13** | 12 | **3** | **0** | 2 | **759.37** | **1744.88** | 12 | **678.37** | **0** |
| 3 | **2** | **0** | 13 | **6** | **0** | 3 | **384.75** | **0** | 13 | **1255.5** | **0** |
| 4 | **3** | **8** | 14 | **28** | **6** | 4 | **843.75** | **1728** | 14 | **2099.25** | **1039.5** |
| 5 | **18** | **9** | 15 | **203** | **53** | 5 | **2635.87** | **1768.5** | 15 | **6480** | **3283.88** |
| 6 | **38** | **20** | 16 | **100** | **22** | 6 | **3179.25** | **2092.5** | 16 | **6773.62** | **3145.5** |
| 7 | **78** | **27** | 17 | **36** | **1** | 7 | **6345** | **3611.25** | 17 | **4060.13** | **297** |
| 8 | **25** | **5** | 18 | **93** | **7** | 8 | **2170.12** | **654.75** | 18 | **5899.5** | **1174.5** |
| 9 | **9** | **7** | 19 | **3** | **3** | 9 | **1920.37** | **1417.5** | 19 | **651.37** | **546.75** |
| 10 | **81** | **15** | 20 | **15** | **2** | 10 | **5146.87** | **1863** | 20 | **1768.5** | **371.25** |

**Table S9.** Comparison of tract number and volume in the right and left hemisphere of the subject on the basis of sex. SN = Subject number F= Female M= Male

| TRACT NUMBER |  |  |  |  |  |  |  | TRACT VOLUME<br>(mm<sup>3</sup>) |  |  |  |  |  |  |  |
| --- | --- | --- | --- | --- | --- | --- | --- | --- | --- | --- | --- | --- | --- | --- | --- |
| Right Hemisphere |  |  |  | Left Hemisphere |  |  |  | Right Hemisphere |  |  |  | Left Hemisphere |  |  |  |
| SN | F | SN | M | SN | F | SN | M | SN | F | SN | M | SN | F | SN | M |
| 1 | 41 | 11 | 39 | 1 | 3 | 11 | 1 | 1 | 99.9 | 11 | 92.49 | 1 | 793.12 | 11 | 249.75 |
| 2 | 3 | 12 | 3 | 2 | 13 | 12 | 0 | 2 | 102.04 | 12 | 678.37 | 2 | 1744.88 | 12 | 0 |
| 3 | 2 | 13 | 6 | 3 | 0 | 13 | 0 | 3 | 81.95 | 13 | 1255.5 | 3 | 0 | 13 | 0 |
| 4 | 3 | 14 | 28 | 4 | 8 | 14 | 6 | 4 | 96.82 | 14 | 2099.25 | 4 | 1728 | 14 | 1039.5 |
| 5 | 18 | 15 | 203 | 5 | 9 | 15 | 53 | 5 | 89.01 | 15 | 6480 | 5 | 1768.5 | 15 | 3283.88 |
| 6 | 38 | 16 | 100 | 6 | 20 | 16 | 22 | 6 | 93.83 | 16 | 6773.62 | 6 | 2092.5 | 16 | 3145.5 |
| 7 | 78 | 17 | 36 | 7 | 27 | 17 | 1 | 7 | 104.55 | 17 | 4060.13 | 7 | 3611.25 | 17 | 297 |
| 8 | 25 | 18 | 93 | 8 | 5 | 18 | 7 | 8 | 85.87 | 18 | 5899.5 | 8 | 654.75 | 18 | 1174.5 |
| 9 | 9 | 19 | 3 | 9 | 7 | 19 | 3 | 9 | 106.79 | 19 | 651.37 | 9 | 1417.5 | 19 | 546.75 |
| 10 | 81 | 20 | 15 | 10 | 15 | 20 | 2 | 10 | 91.87 | 20 | 1768.5 | 10 | 1863 | 20 | 371.25 |

**Table S10.**
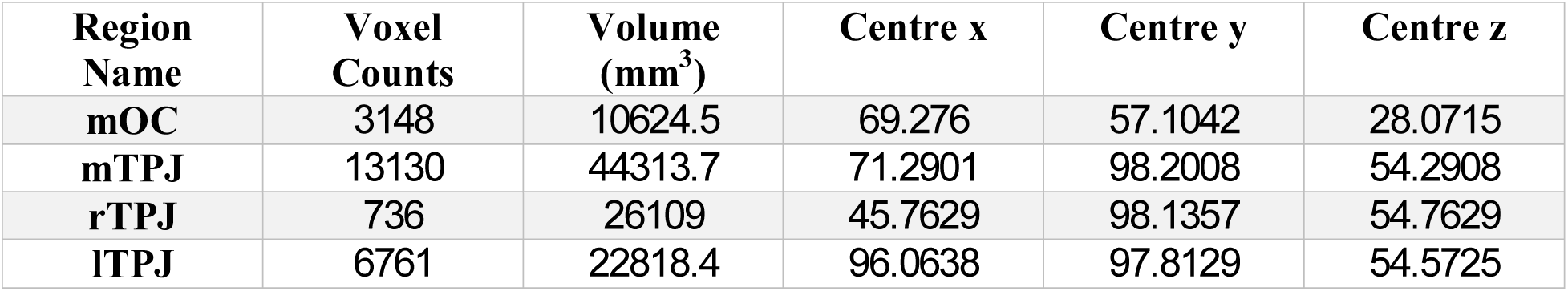
Regional statistics.

**Graph S1.**
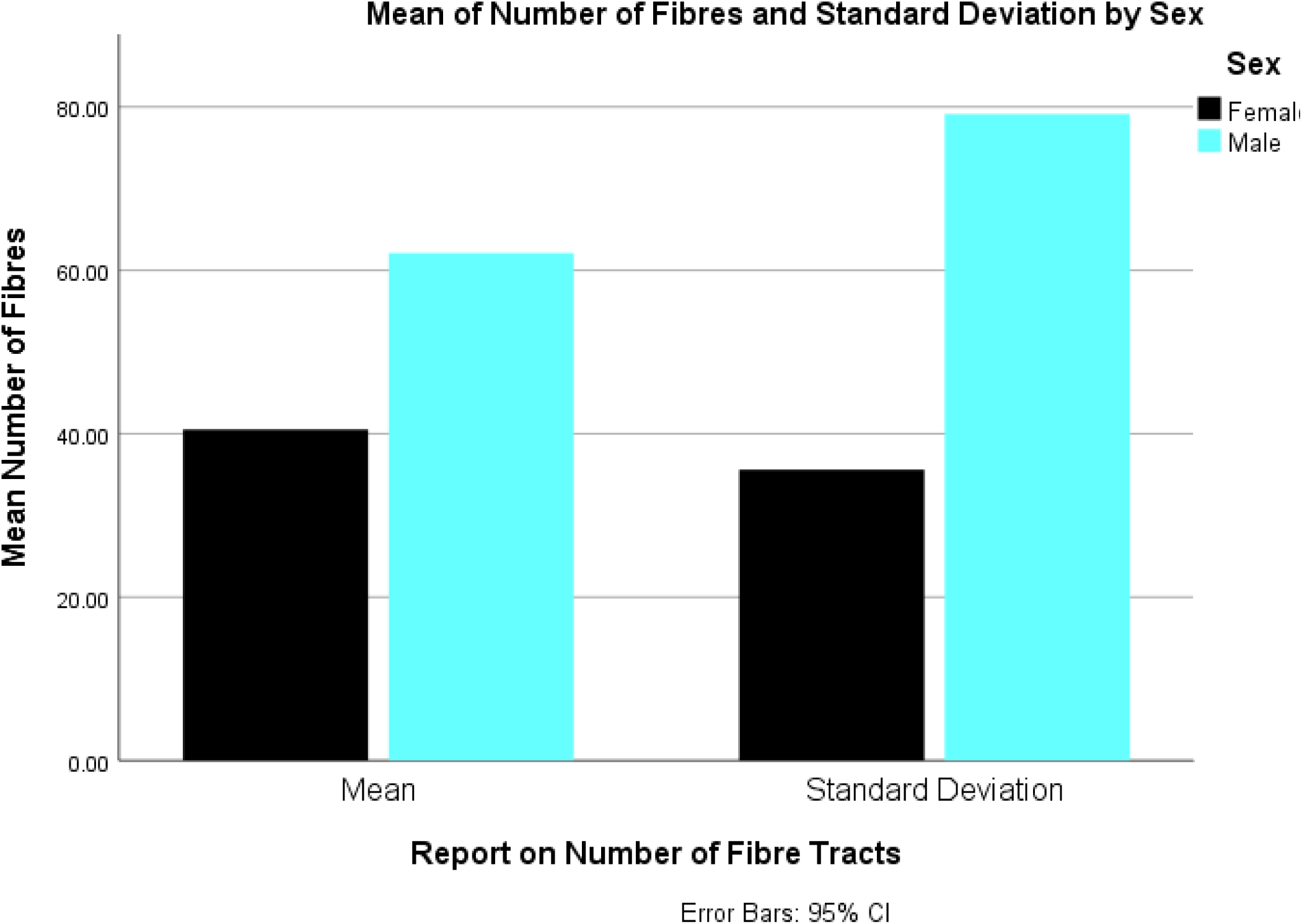
Mean number of fibres and standard deviation in men and women. This graph displays the results of a comparative analysis between the number of fibres in male and female subjects. Here, Females show, mean = 40.50 Standard deviation = 35.60 Males show, mean = 62.10 Standard deviation = 79.12

**Graph S2.**
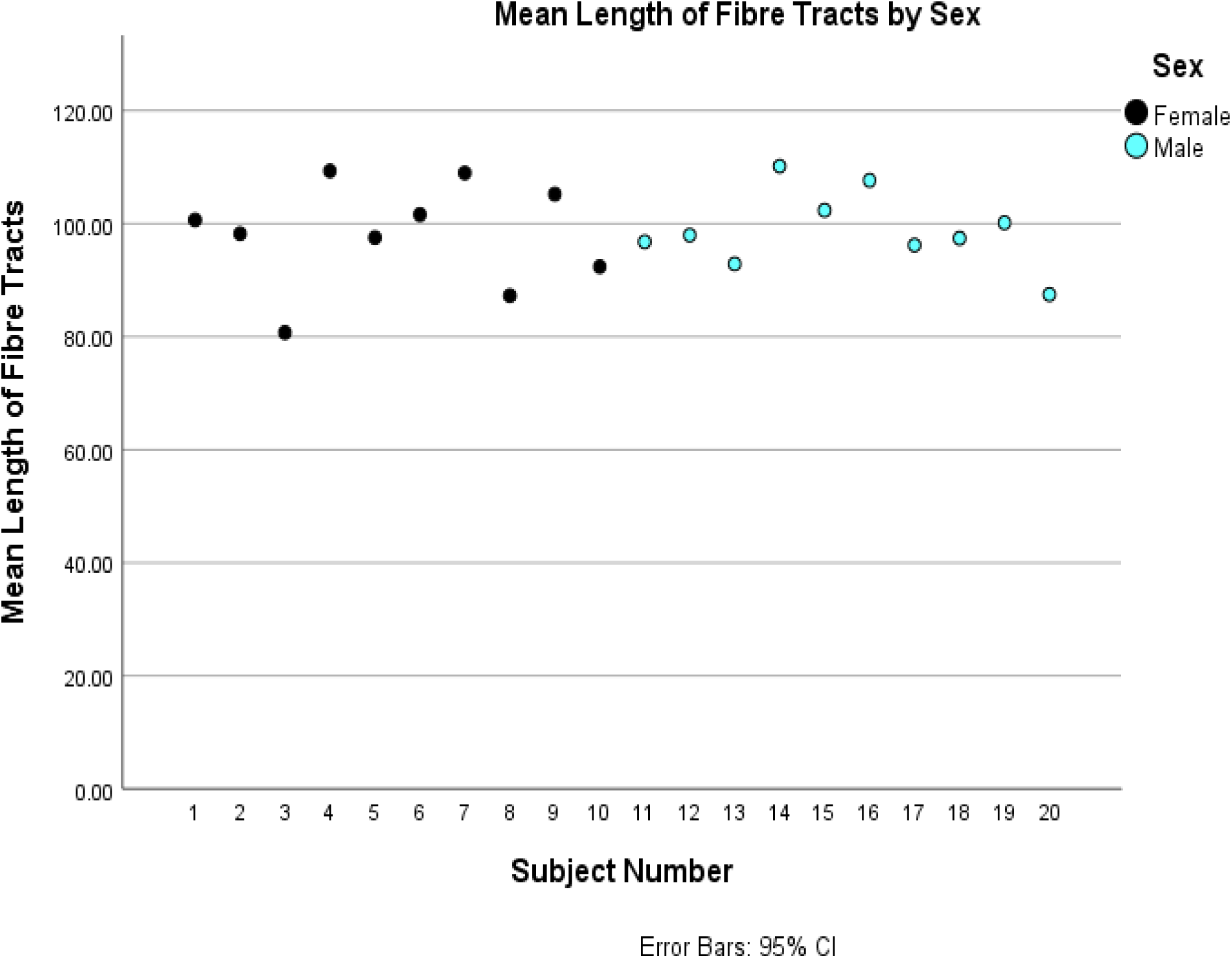
Mean length of fibres in subjects by sex. Here the mean length (mm) of all subjects have been displayed. There is no significant difference observed when male and female subjects are compared.

**Graph S3.**
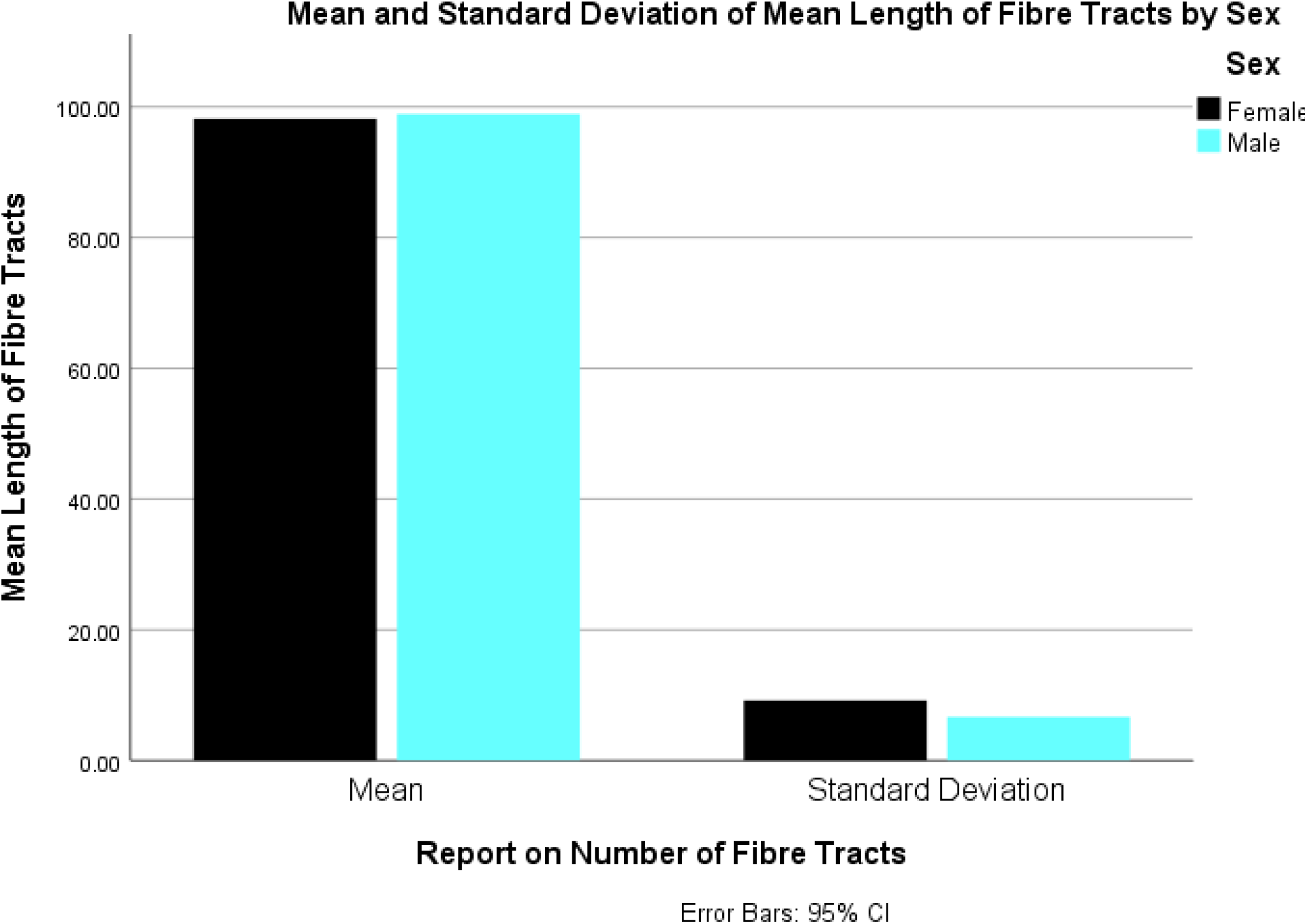
Mean of lengths of fibre tracts and their standard deviation in men and women. This graph displays the results of a comparative analysis between the mean length of fibre tracts in male and female subjects. Here, Female, mean length = 98.19mm Standard deviation = 9.23mm Male, mean length = 98.90mm Standard deviation = 6.66mm

**Graph S4.**
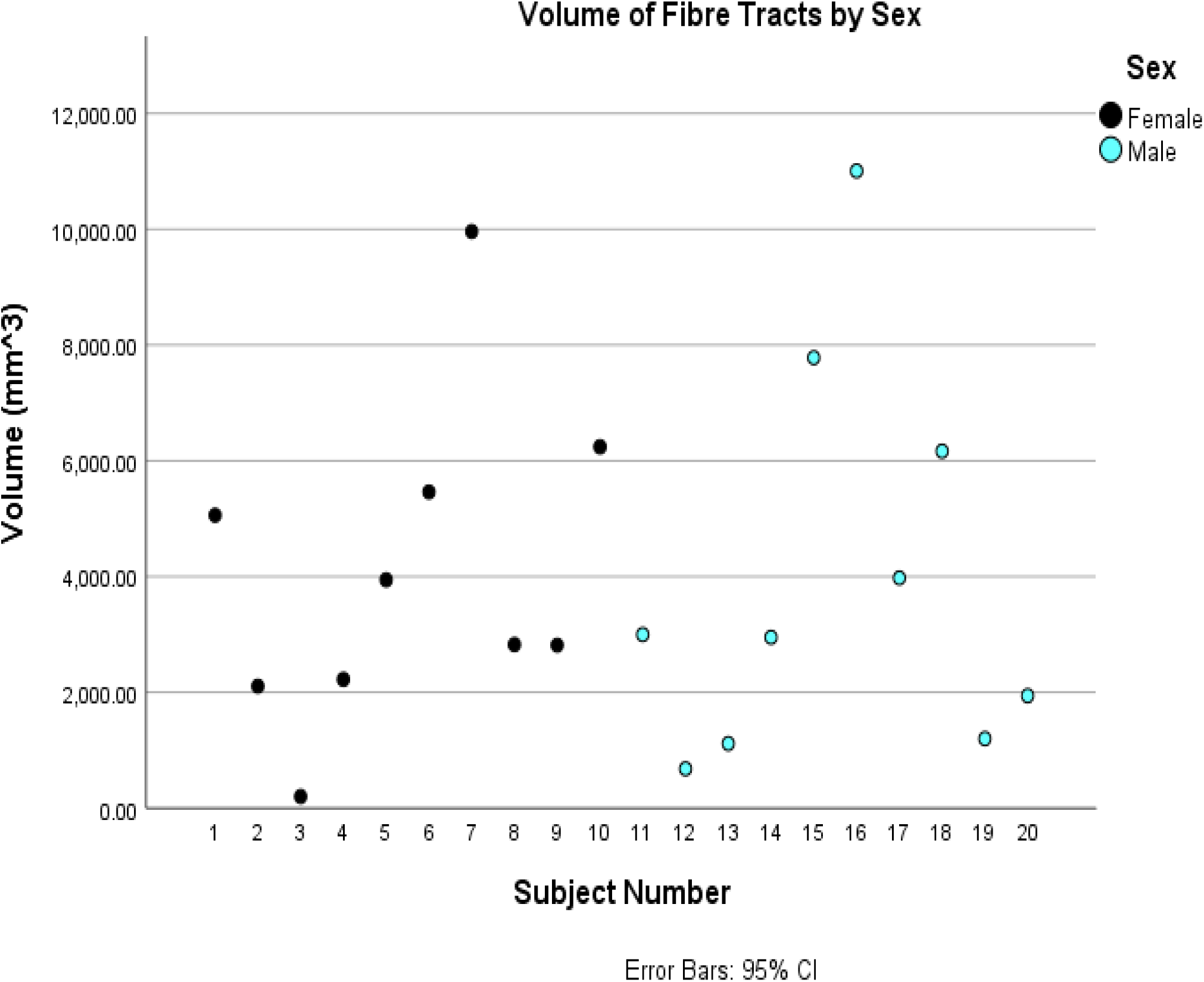
Volume of fibre tracts in subjects by sex. Here the volume (mm^3^) of all subjects have been displayed. There is no significant difference observed when male and female subjects are compared.

**Graph S5.**
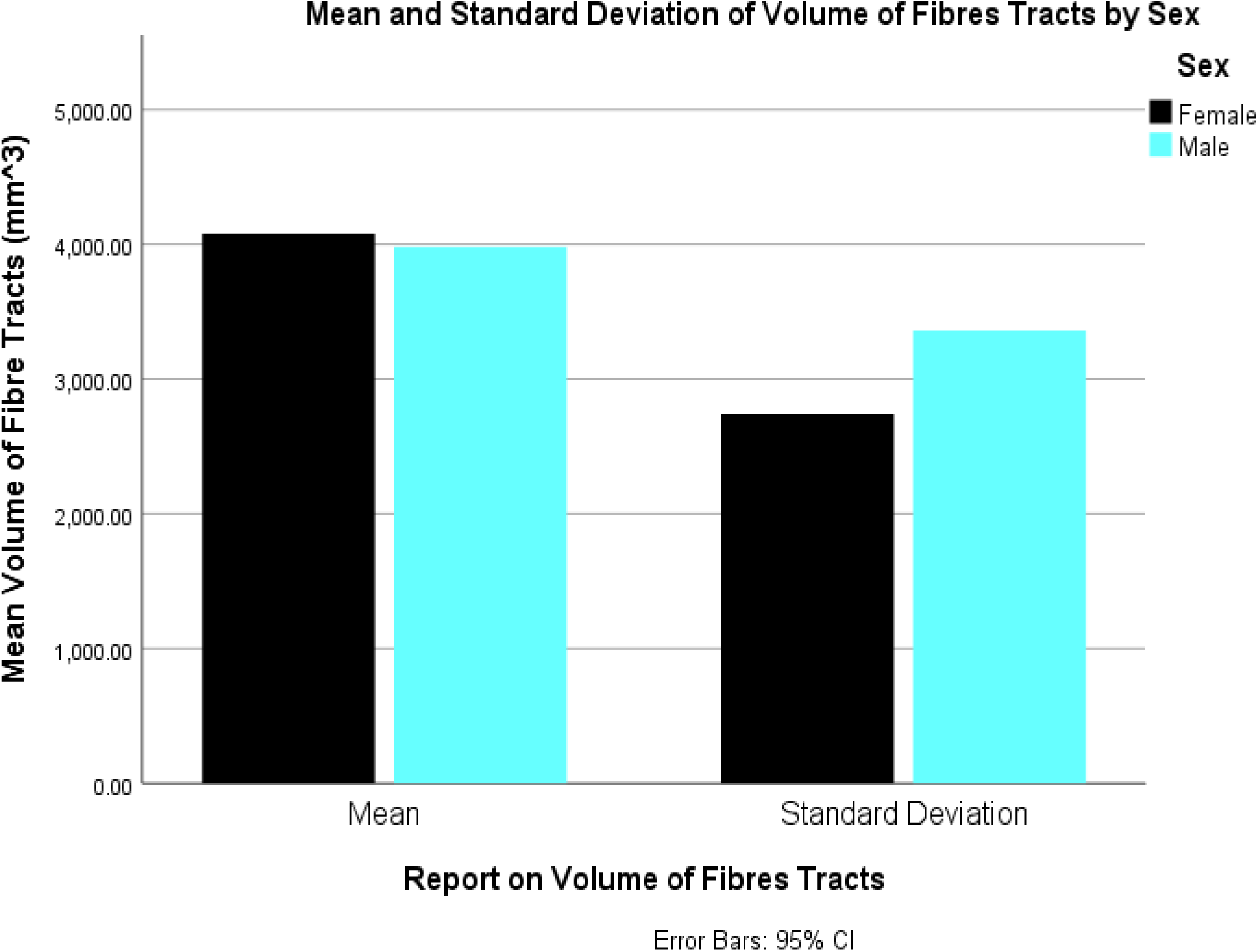
Mean of volume of fibre tracts and their standard deviation in men and women. This graph displays the results of a comparative analysis between the volume of fibre tracts in male and female subjects. Here, Female, mean length = 4083.07 mm^3^ Standard deviation = 2742.66 mm^3^ Male, mean length = 3980.4760 mm^3^ Standard deviation = 3360.21 mm^3^

**Graph S6.**
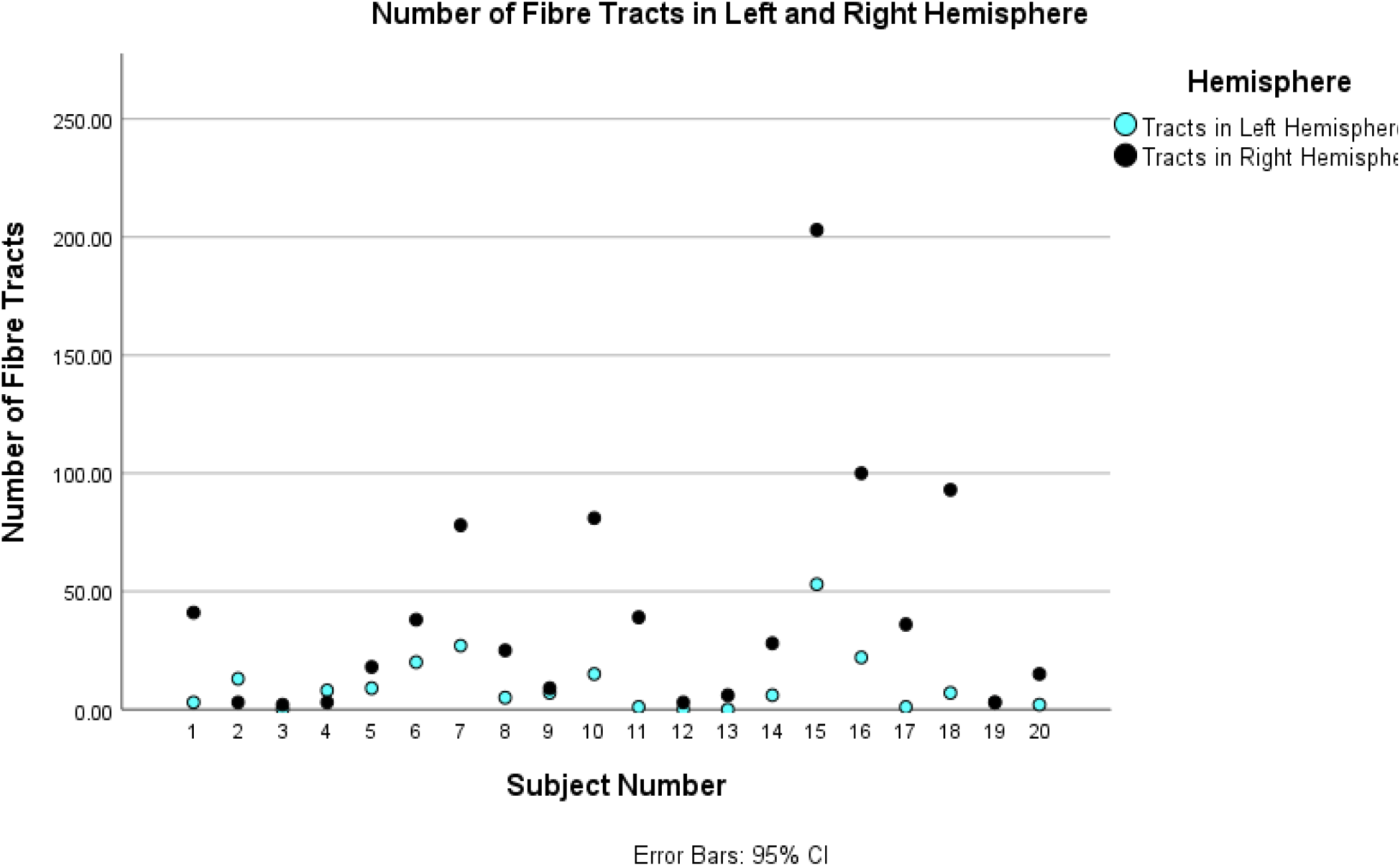
Number of fibre tracts observed in left and right hemisphere. In subject number 2 and 4, we see that a greater number of fibres are found in the left hemisphere, in the remaining 18 subjects, a greater number of fibres have been found in the right hemisphere as compared to left.

**Graph S7.**
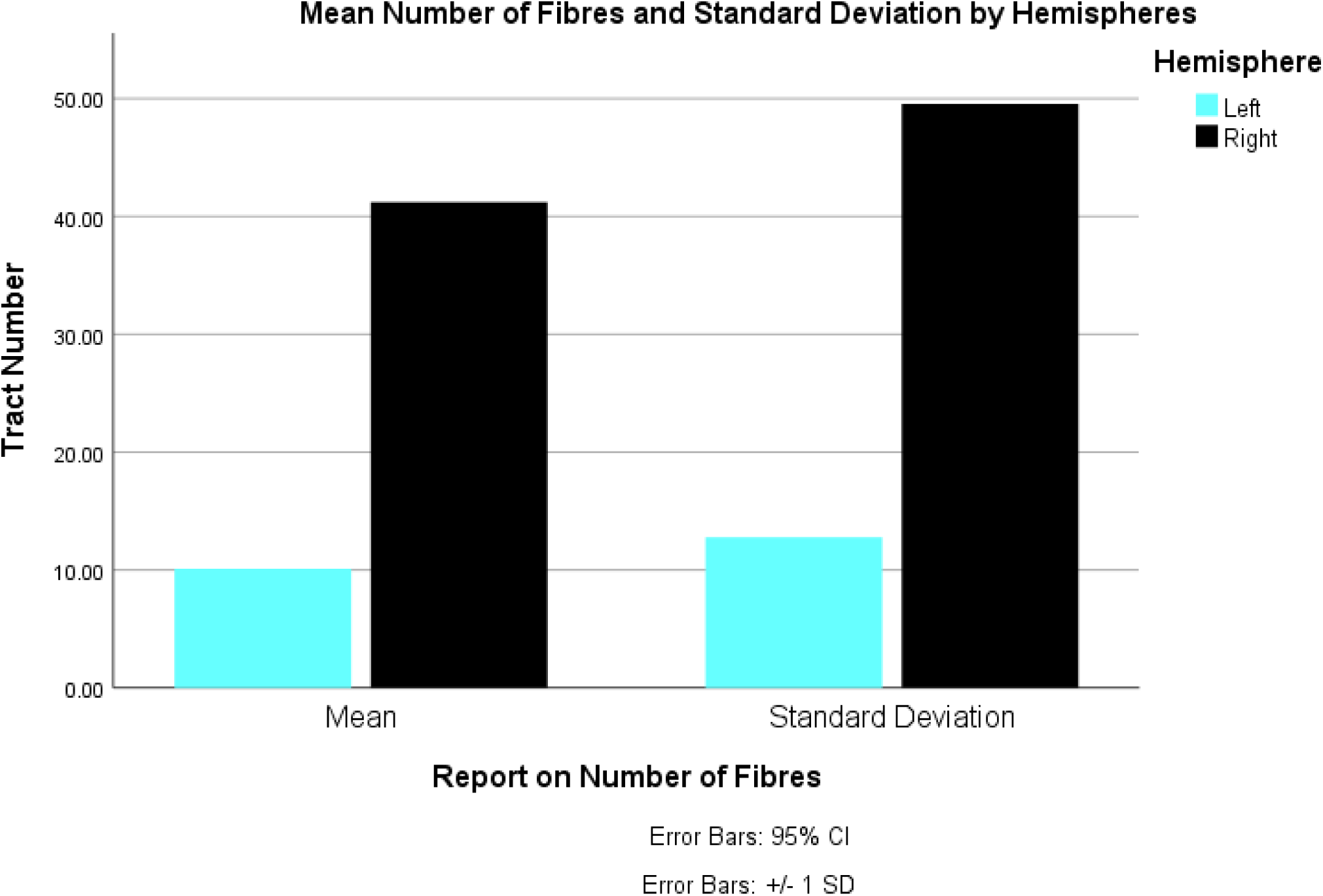
Mean of number of fibre tracts and their standard deviation in left and right hemispheres. This graph displays the results of a comparative analysis between the volume of fibre tracts in the two hemispheres of the subjects. Here, Left, mean length = 10.10 Standard deviation = 12.78 Right, mean length = 41.20 Standard deviation = 49.53

**Graph S8.**
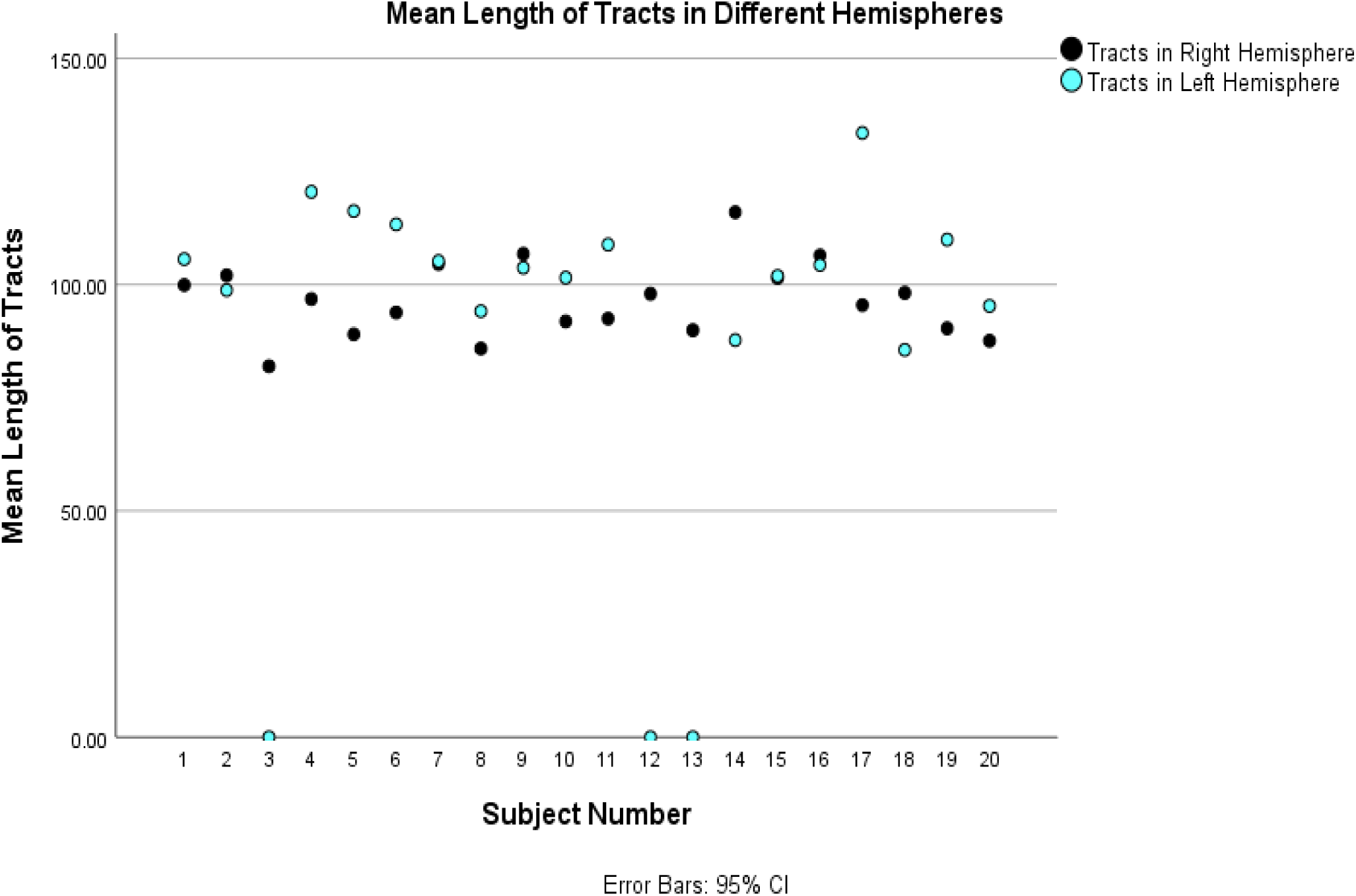
Mean length of tracts in different hemispheres. This graph displays the mean length of tracts observed in all the subjects, and compares them on the basis of hemispheres. Here we see that, 8 of the subjects show more average length of fibres in the right hemispheres while 12 shows more average length in the left hemisphere.

**Graph S9.**
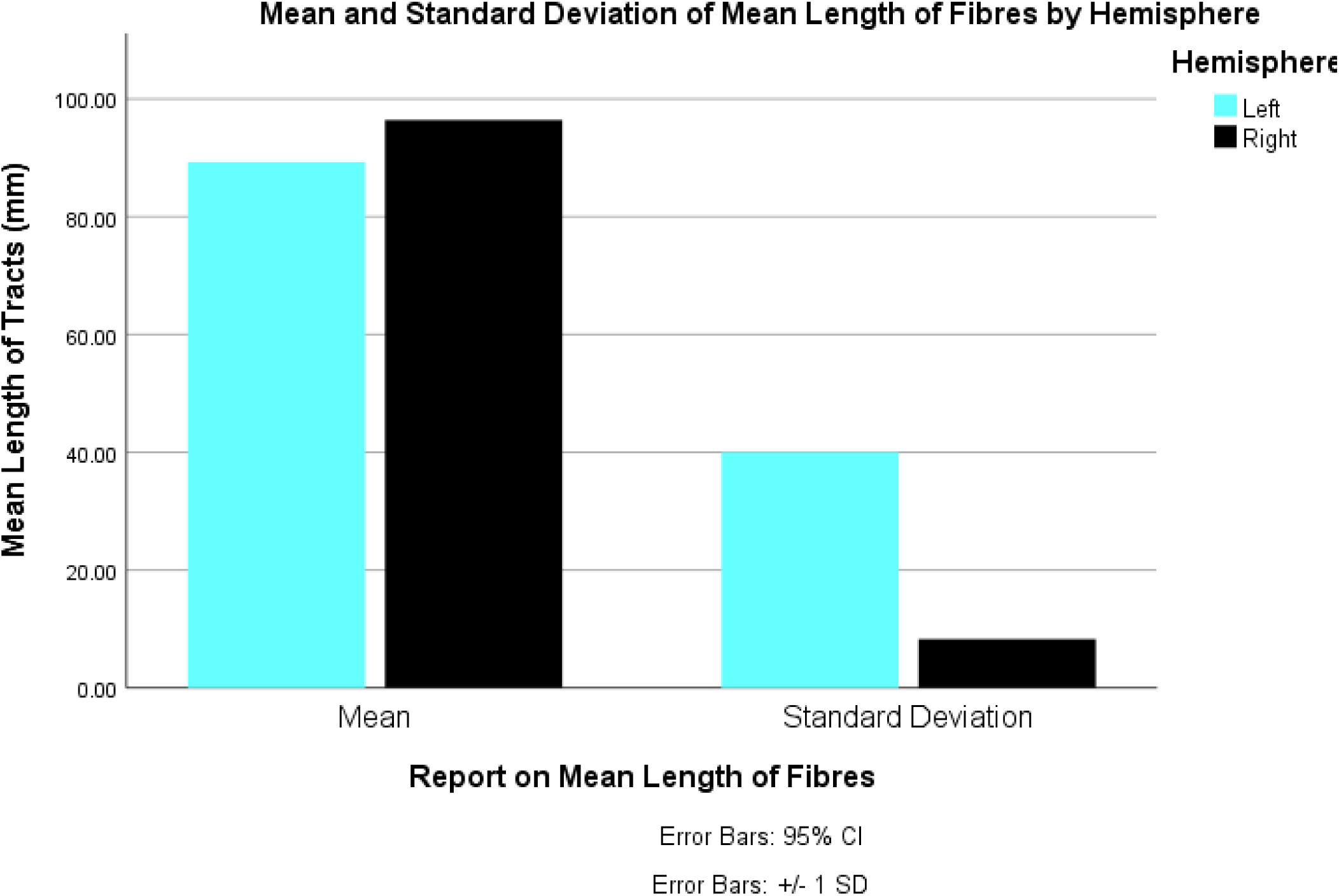
Mean and standard deviation of mean length of fibre tracts in different hemispheres. This graph displays the results of a comparative analysis between the mean length of fibre tracts in the two hemispheres of the subjects. Here, Left, mean length = 89.30 mm Standard deviation = 39.99 mm Right, mean length = 96.42 mm Standard deviation = 8.29 mm

**Graph S10.**
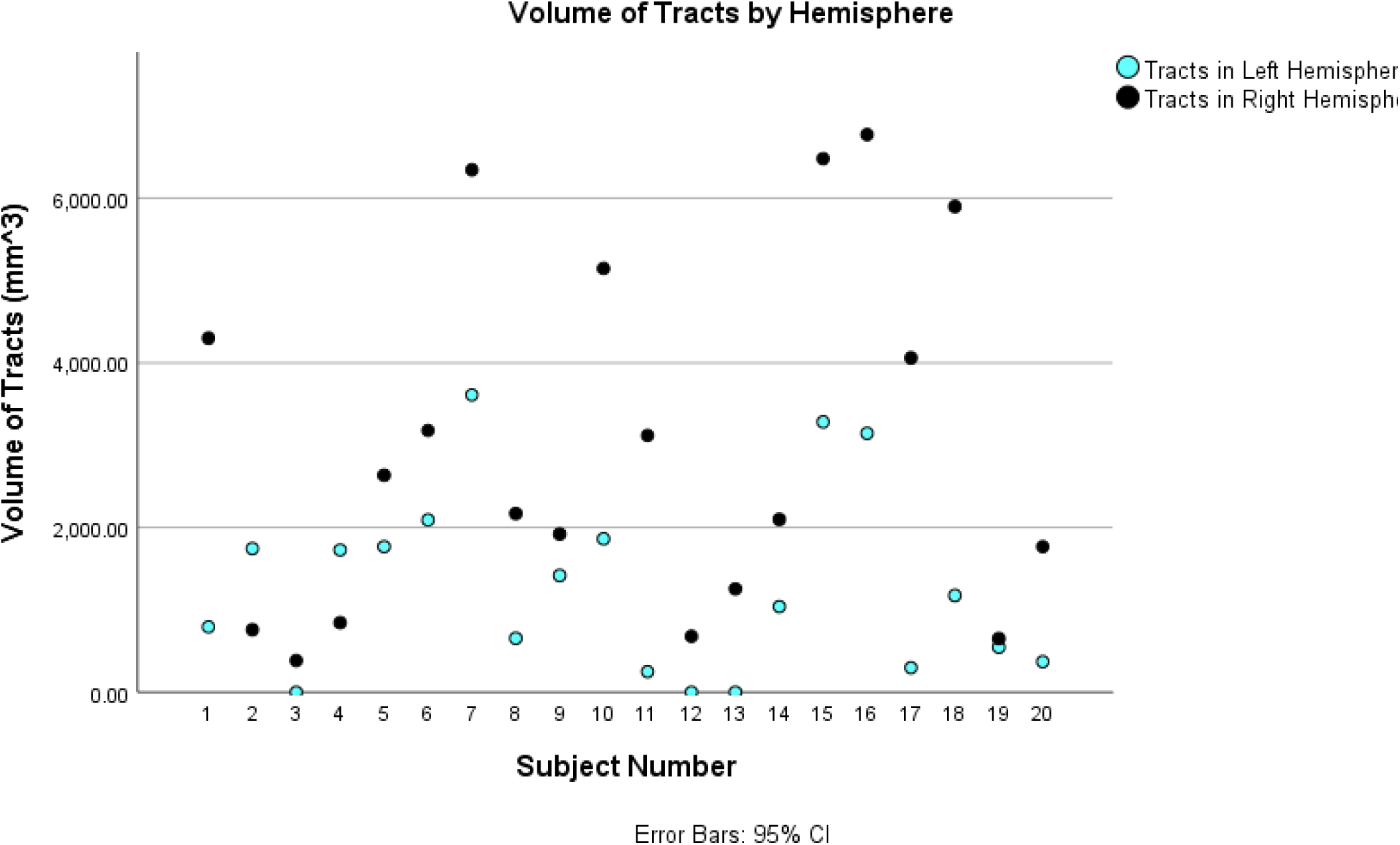
Volume of Tracts by Hemisphere. This graph displays the volume of tracts observed in all the subjects, and compares them on the basis of hemispheres. Here we see that, 18 of the subjects show more volume of fibres in the right hemispheres.

**Graph S11.**
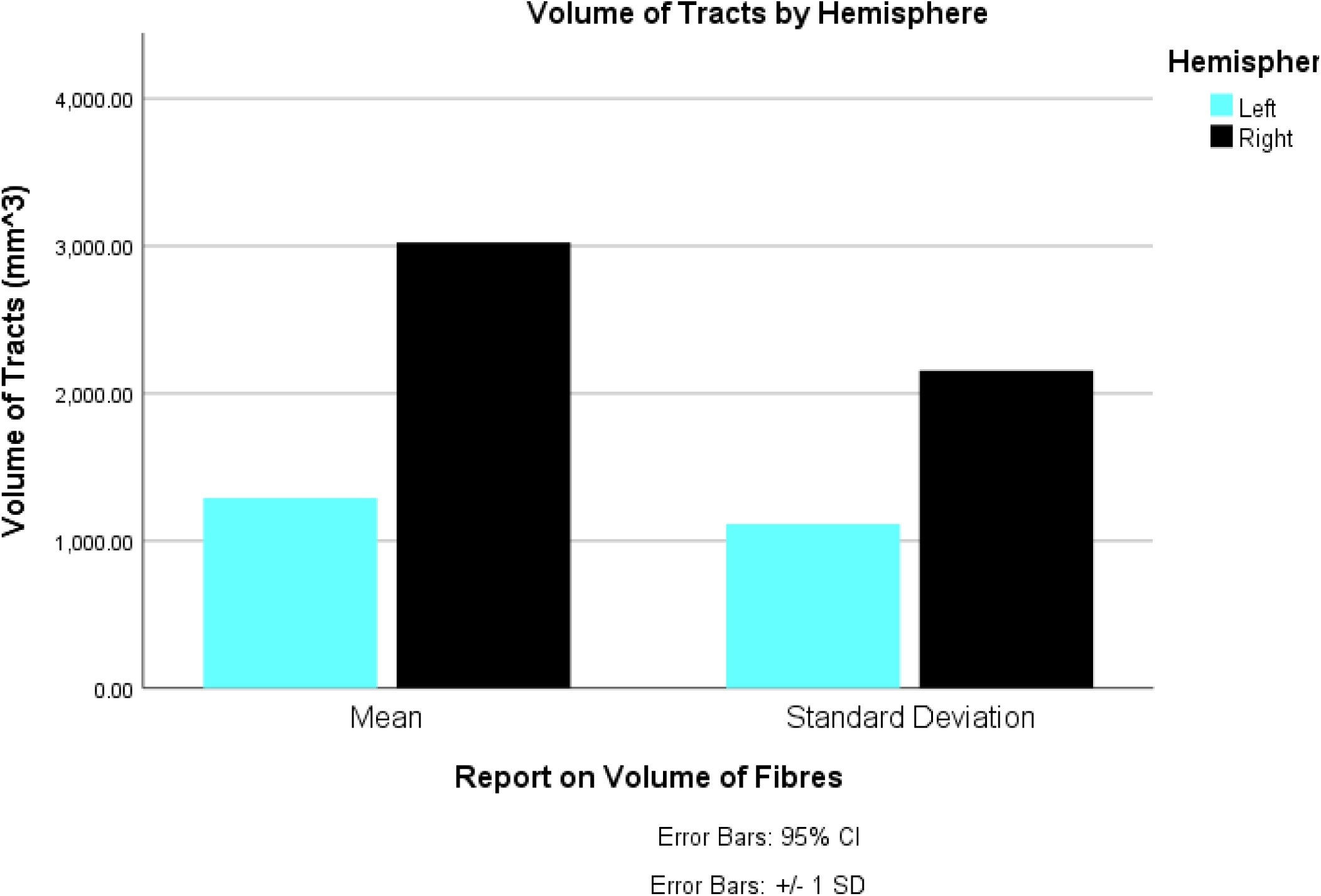
Mean and standard deviation of volume of fibre tracts in different hemispheres. This graph displays the results of a comparative analysis between the volume of fibre tracts in the two hemispheres of the subjects. Here, Left, mean length = 1289.08 mm^3^ Standard deviation = 1114.65 mm^3^ Right, mean length = 3023.49 mm^3^ Standard deviation = 2154.28 mm^3^

**Graph S12.**
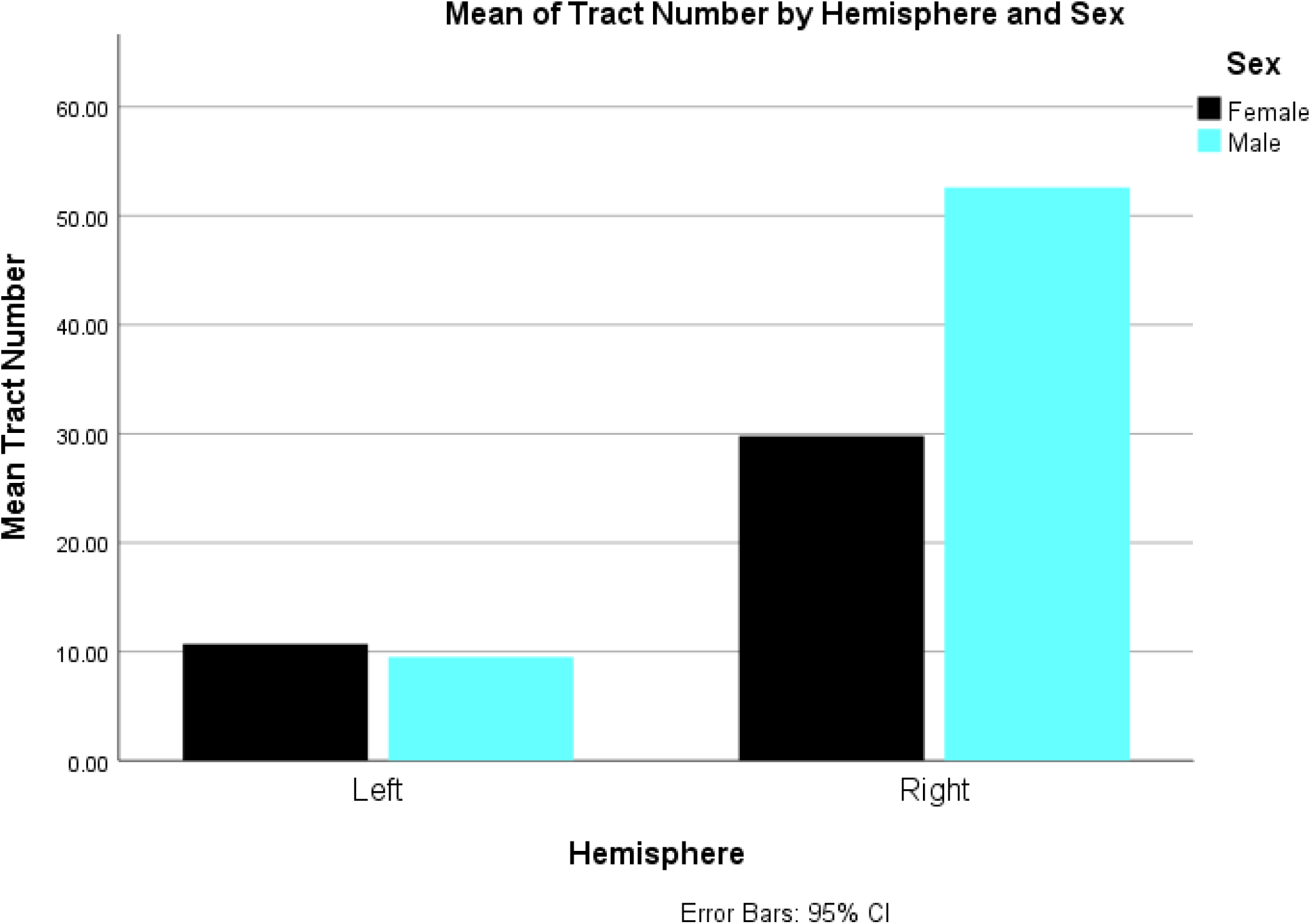
Mean of tract number by hemisphere and sex. This graph displays the results of a comparative analysis between the mean of fibre tract numbers in the two hemispheres between male and female subjects. Here, Females in left hemisphere show = 10.70 fibres Males in left hemisphere show = 9.50 fibres Females in right hemisphere show = 29.8 fibres Males in right hemisphere show = 52.60 fibres

**Graph S13.**
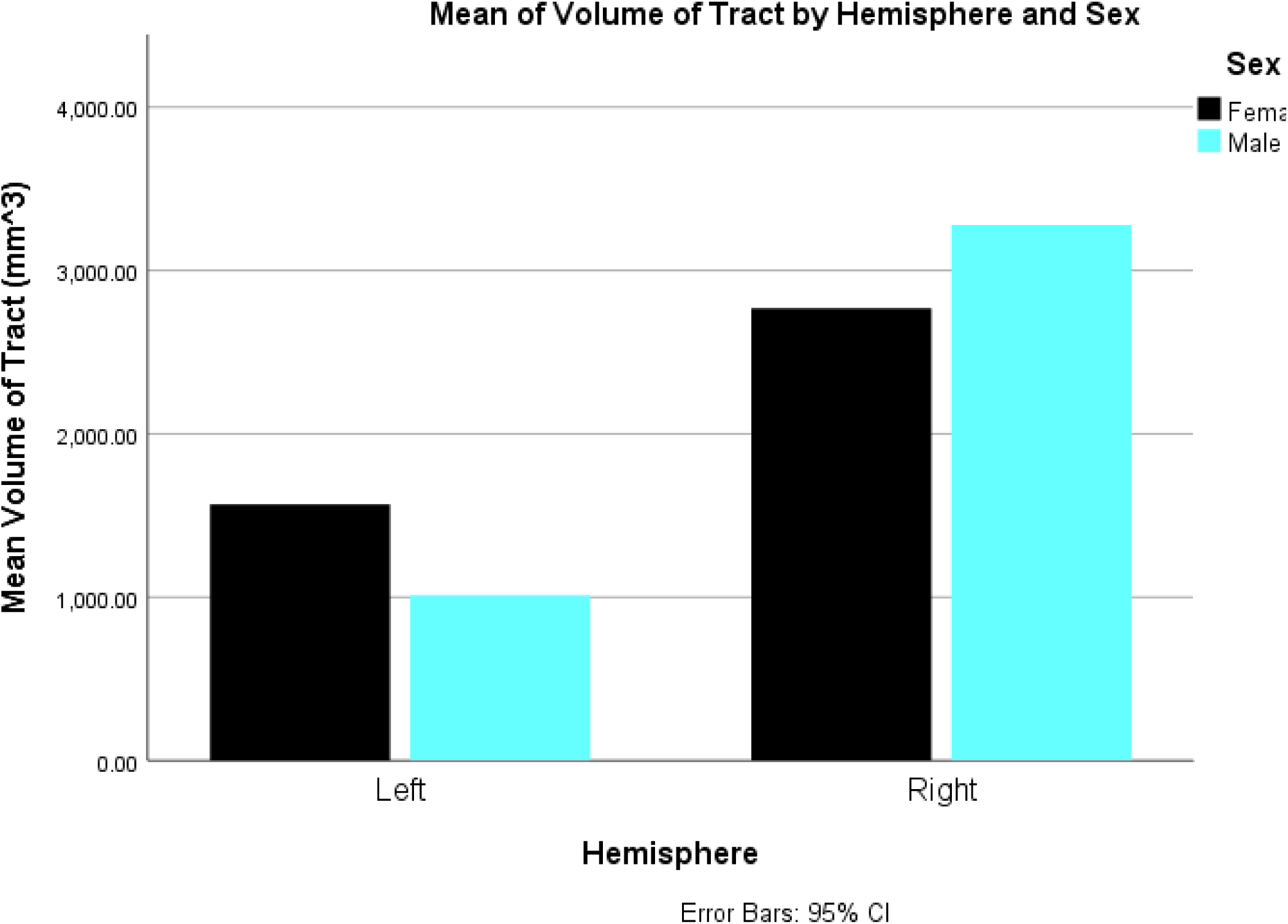
Mean of volume of tract by hemisphere and sex. This graph displays the results of a comparative analysis between the mean of volume of fibre tracts in the two hemispheres between male and female subjects. Here, Females in left hemisphere show = 1567.35 mm^3^ Males in left hemisphere show = 1010.81 mm^3^ Females in right hemisphere show = 2768.51 mm^3^ Males in right hemisphere show = 3278.47 mm^3^

**Graph S14.**
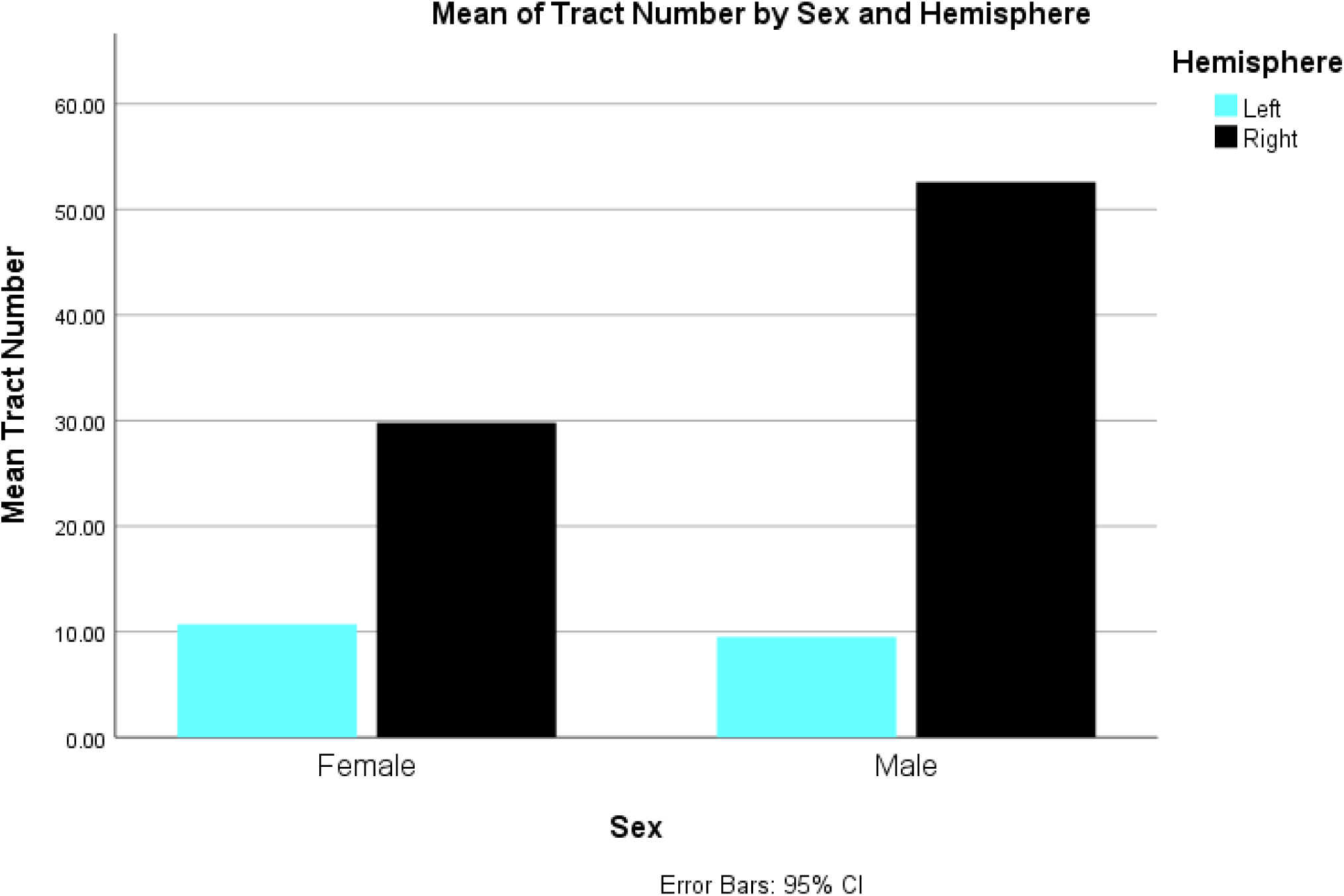
Mean of tract number by sex and hemisphere. This graph displays the results of a comparative analysis between the mean of number of fibre tracts of the sexes in different hemispheres. Here, Females in left hemisphere show = 10.70 fibres right hemisphere show = 29.80 fibres Males in left hemisphere show = 9.50 fibres right hemisphere show = 52.60 fibres

**Graph S15.**
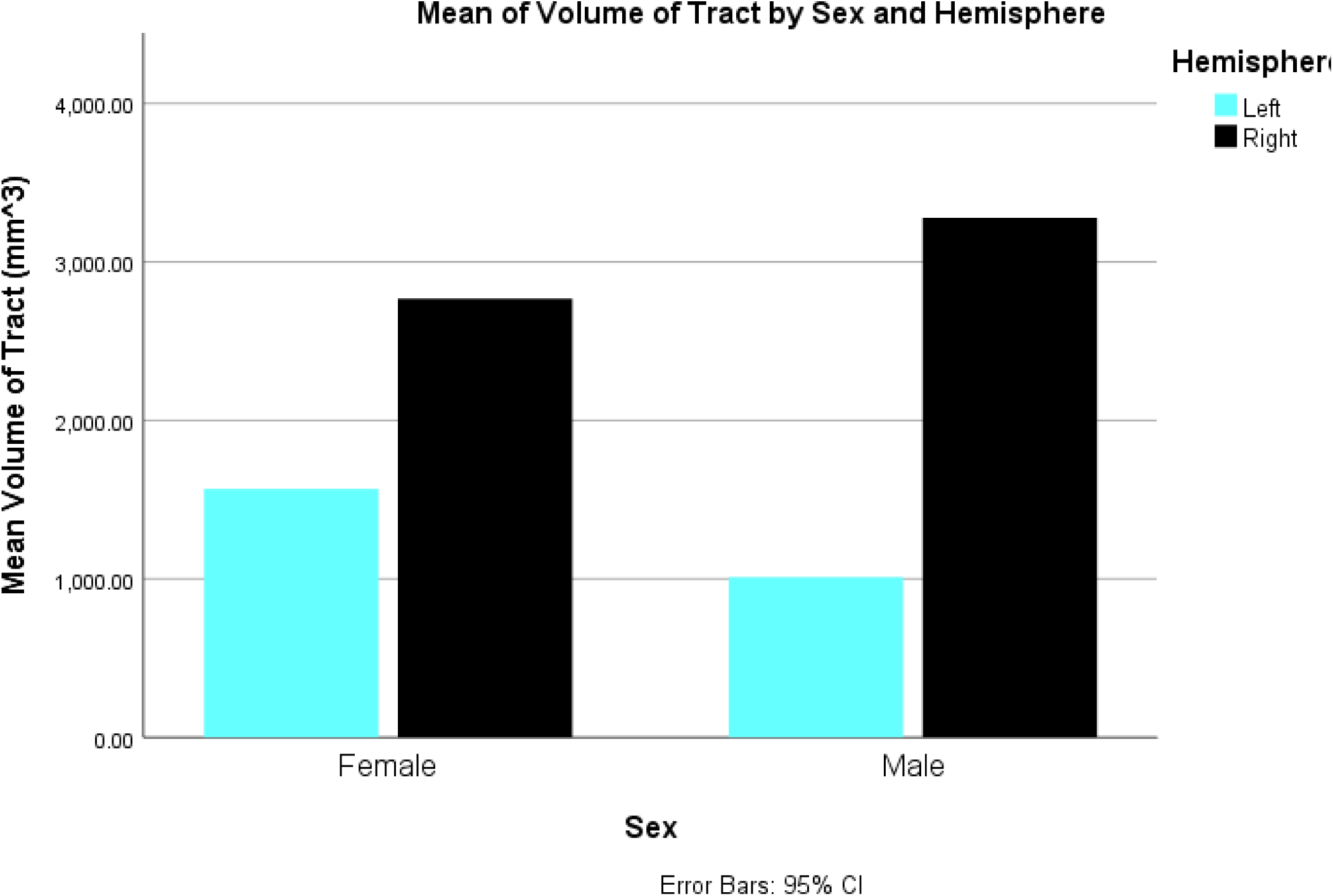
Mean of volume of tract by sex and hemisphere. This graph displays the results of a comparative analysis between the mean of volume of fibre tracts of the sexes in different hemispheres. Here, Females in left hemisphere show = 1567 mm^3^ right hemisphere show = 2768.51 mm^3^ Males in left hemisphere show = 1010.81 mm^3^ right hemisphere show = 3278.47 mm^3^

**Graph S16.**
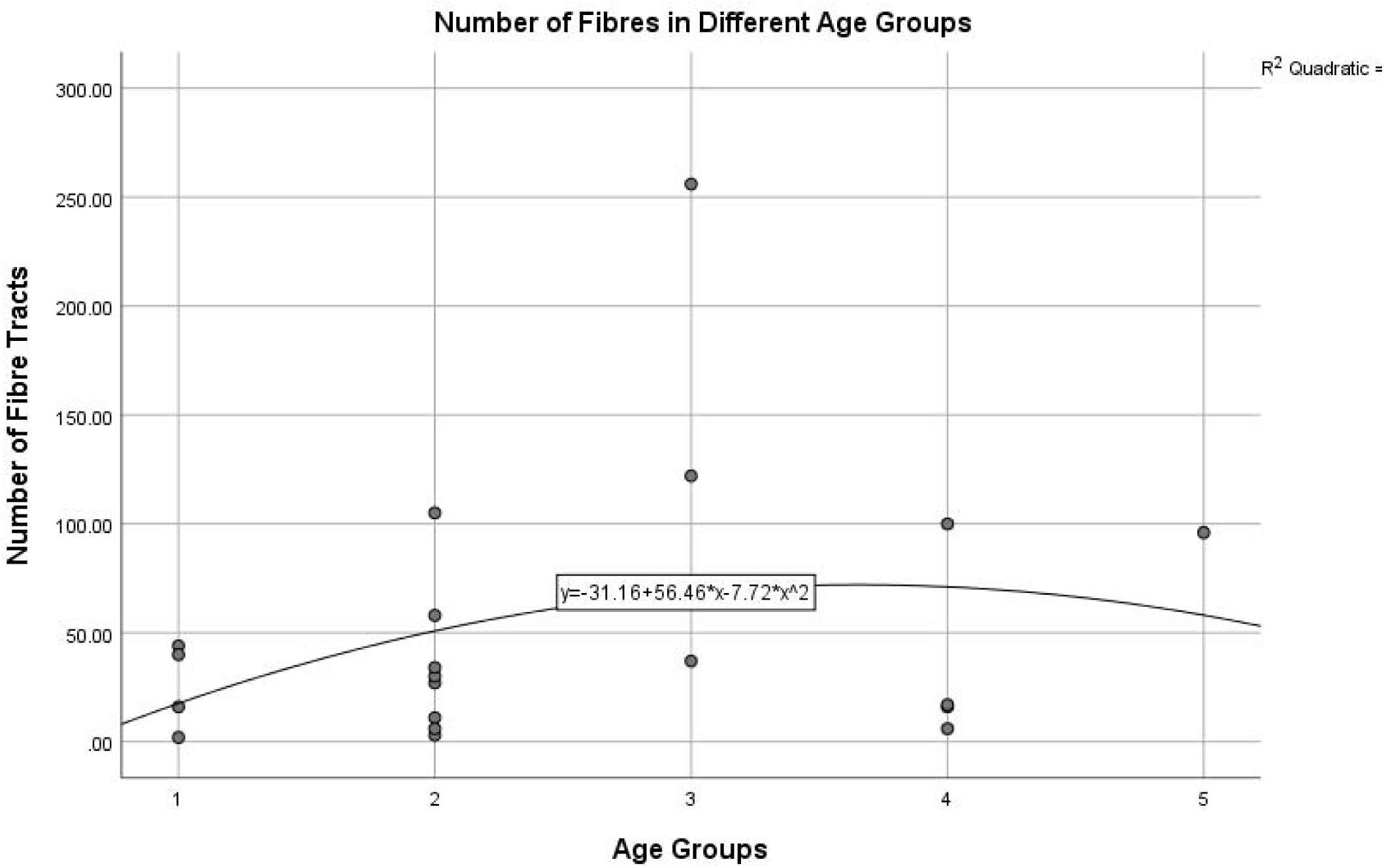
Number of Tracts in Different Age Groups. In this scatter plot, we see the number of fibre tracts divided in different age groups. Here age group, 1 = 20-24 years 2 = 25-29 years 3 = 30-34 years 4 = 35-39 years 5 = 40-44 years A gradual increase and then decrease is spotted in the pattern of distribution, as signified by the quadratic curve.

**Graph S17.**
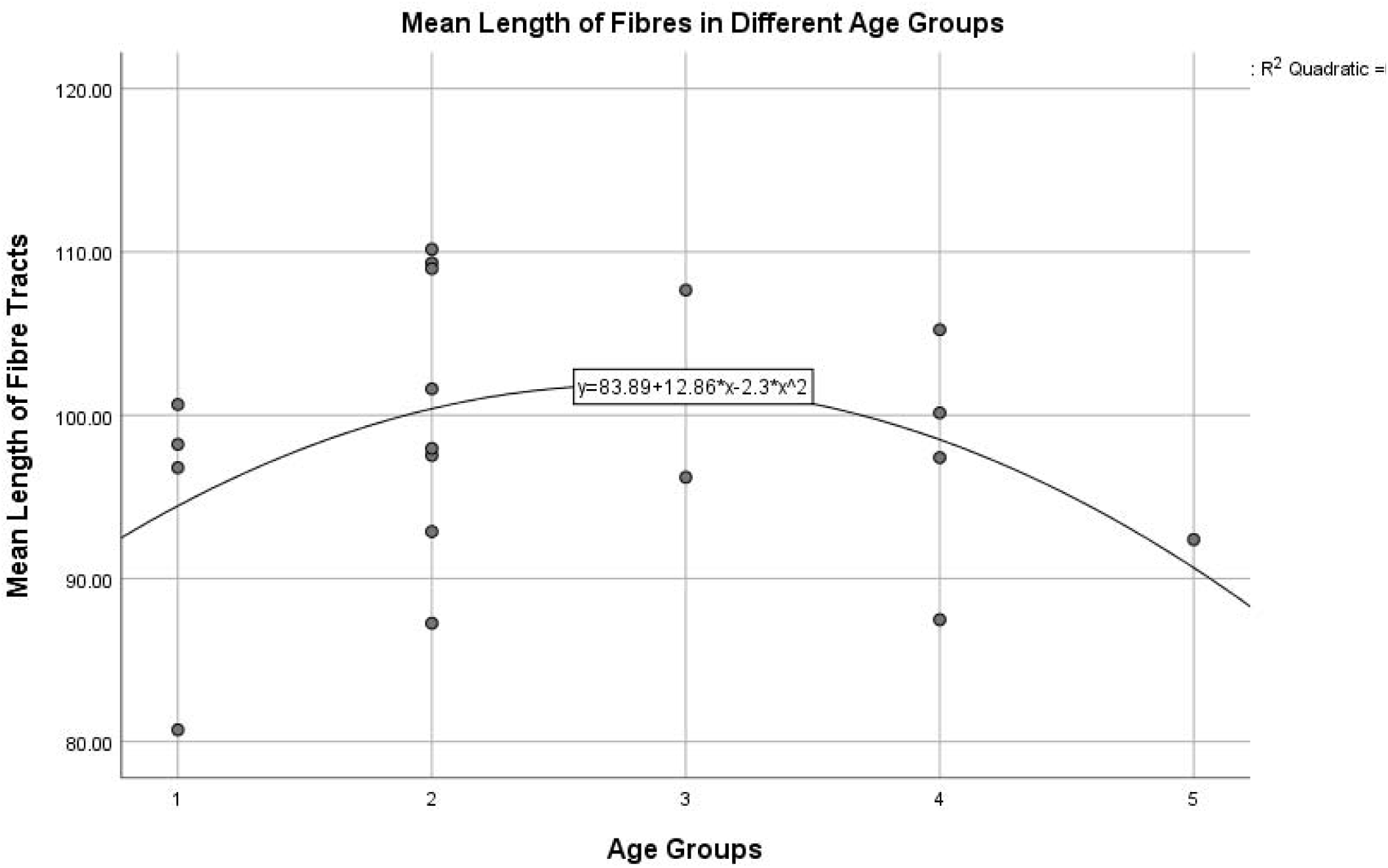
Mean Length of Tracts in Different Age Groups. In this scatter plot, we see the mean length of fibre tracts divided in different age groups. Here age group, 1 = 20-24 years 2 = 25-29 years 3 = 30-34 years 4 = 35-39 years 5 = 40-44 years A gradual increase and then decrease is spotted in the pattern of distribution, as signified by the quadratic curve.

**Graph S18.**
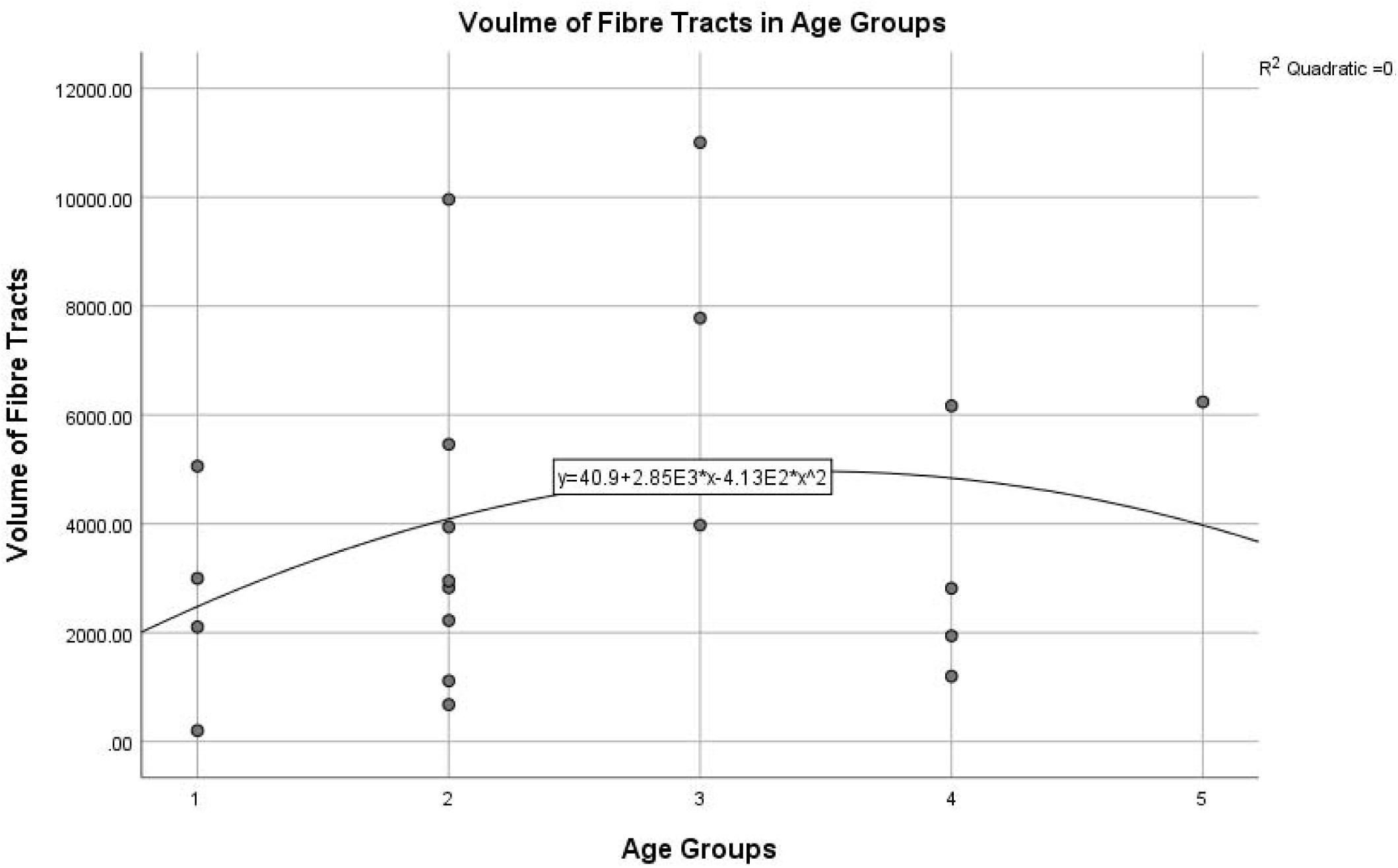
Volume of Tracts in Different Age Groups. In this scatter plot, we see the volume of fibre tracts divided in different age groups. Here age group, 1 = 20-24 years 2 = 25-29 years 3 = 30-34 years 4 = 35-39 years 5 = 40-44 years A gradual increase and then decrease is spotted in the pattern of distribution, as signified by the quadratic curve.

